# Harnessing Uniform Design to Enhance AI-Driven Predictions of Physicochemical Properties of Short Peptides

**DOI:** 10.1101/2025.03.10.642308

**Authors:** Zhihui Zhu, Huapeng Liu, Yongfu Guo, Mouzheng Xu, Xuechen Li, Haojin Zhou, Jiaqi Wang

## Abstract

Short peptides hold significant promise in drug discovery and materials science due to their biocompatibility, multifunctionality, and ease of synthesis. However, accurately predicting their physicochemical properties, a prerequisite for application development, remains a challenge. This study presents an innovative approach integrating uniform design (UD) with artificial intelligence (AI) to enhance prediction of key physicochemical properties, including aggregation propensity (AP), hydrophilicity (logP), and isoelectric point (pI). Using UD, we generate 31 distinct peptide datasets, with a consistent amino acid occupation fraction of 5% at each position, thereby creating unbiased training data for AI models. The performance of each AI model is rigorously evaluated using various testing schemes, and optimal sample sizes are determined for accurate prediction of each property. Additionally, Shapley Additive Explanations (SHAP) analysis identifies aromaticity, logP, net charge, and pI as the primary factors affecting peptide aggregation. This work provides comprehensive datasets on the physicochemical properties of all tetrapeptides, develops robust AI-based predictive models, and elucidates the relationships between key physicochemical characteristics and self-assembly behavior. By integrating experimental design, AI modeling, and peptide domain knowledge, our approach facilitates the discovery and optimization of functional peptides, offering new opportunities for peptide-based therapeutic applications.

## 1. Introduction

Peptides, essential biomolecules, have attracted considerable interest in both academia and industry due to their biocompatibility^1^, multifunctionality^2^, and therapeutic potential^3^. Their ability to self-assemble into functional nanostructures has further expanded their applications across diverse fields, including drug delivery, tissue engineering, bioimaging, antimicrobial therapy, and more^4^. In drug delivery, self-assembled peptide nanostructures serve as efficient carriers for therapeutic agents, enhancing efficacy while minimizing side effects^5,6^. In tissue engineering, they function as bioscaffolds that promote cell proliferation and tissue regeneration^7,8^. Additionally, peptide self-assemblies can be engineered for non-invasive in vivo bioimaging, enabling real-time visualization of biological processes^9^. These peptides also exhibit antimicrobial properties, offering novel strategies against antibiotic-resistant bacteria^10^. Despite these diverse applications, a major challenge remains in accurately predicting peptide self-assembly behavior and elucidating its intrinsic correlations with other physicochemical properties.

In recent years, the integration of artificial intelligence (AI, all abbreviations and acronyms are listed in the Supplementary file of “List of Abbreviations and Acronyms”) with molecular dynamics (MD) simulations has emerged as a prominent approach for predicting the self-assembly behavior of peptides^4,11,12^. MD simulations generate high-quality data under consistent conditions, which can be leveraged as training datasets to develop AI models for predicting self-assembly sequences. For example, Batra et al.^13^ combined Monte Carlo tree search algorithms with coarse-grained MD (CGMD) simulations to explore potential self-assembly sequences within the complete sequence space of pentapeptides. They employed the CGMD-generated data to train AI models, which were then used to predict the aggregation tendencies of other pentapeptide sequences. Additionally, Wang et al.^14^ developed a combined framework of CGMD simulation and Transformer-based deep learning (DL) to predict the aggregation properties of pentapeptides, decapeptides, and their mixtures. The coupled CGMD-DL approach provided valuable insights into the aggregation behaviors of these peptides, and a transferability study was conducted to expedite the discovery and design of self-assembled peptide systems. Chen et al.^15^ utilized variational autoencoder and Metropolis-Hastings sampling algorithms to generate potential peptide inhibitor sequences. By integrating generative DL with MD simulations, they designed target-specific peptidase inhibitors. These approaches represent a powerful synergy between advanced computational algorithms and data-driven modeling, significantly enhancing the exploration of the vast peptide sequence space and offering new perspectives and sequence libraries for future biological research and development.

However, the sampling methods and the representativeness of the training data in the aforementioned studies have not been sufficiently justified. It is important to note that the accuracy of AI models, not only those predicting peptide properties, is highly dependent on the quality and representativeness of the training data^16^. As peptide length increases, the diversity of possible sequences within the complete sequence space grows exponentially, incurring substantial challenges for accurate predictions. Specifically, the training data may fail to adequately capture the intricate interplay between peptide sequences and their various physicochemical properties^16^. Inadequate sampling of training data can introduce bias toward specific amino acid sequences or properties, thereby limiting the model’s generalization ability^17^.

Consequently, the implementation of a robust experimental design, which has been largely overlooked in previous research, is essential for enhancing the generalization and accuracy of AI models in predicting the physicochemical properties of peptides.

This study introduces an innovative methodology that integrates uniform design (UD) and AI to optimize predictive models for short peptide properties (**Figure 1**). By strategically selecting peptide sequences, we aim to ensure that the data used for training AI models adequately represents the overall distribution of the complete peptide sequence space. To achieve this, 31 sets of UD samples are generated, maintaining an amino acid occupation frequency of 0.05 (equivalent to 1/20) at each position, thereby providing balanced and representative training data for AI models, including Random Forest (RF)^18^, Support Vector Machine (SVM)^19^, and Transformer models^20^. The performance of the AI models is evaluated using both fixed and non-fixed testing datasets (see Experimental Section for details), and the optimal sample size for predicting each physicochemical property, including aggregation propensity (AP), hydrophilicity (logP), and isoelectric point (pI), is determined.

**Figure 1.**
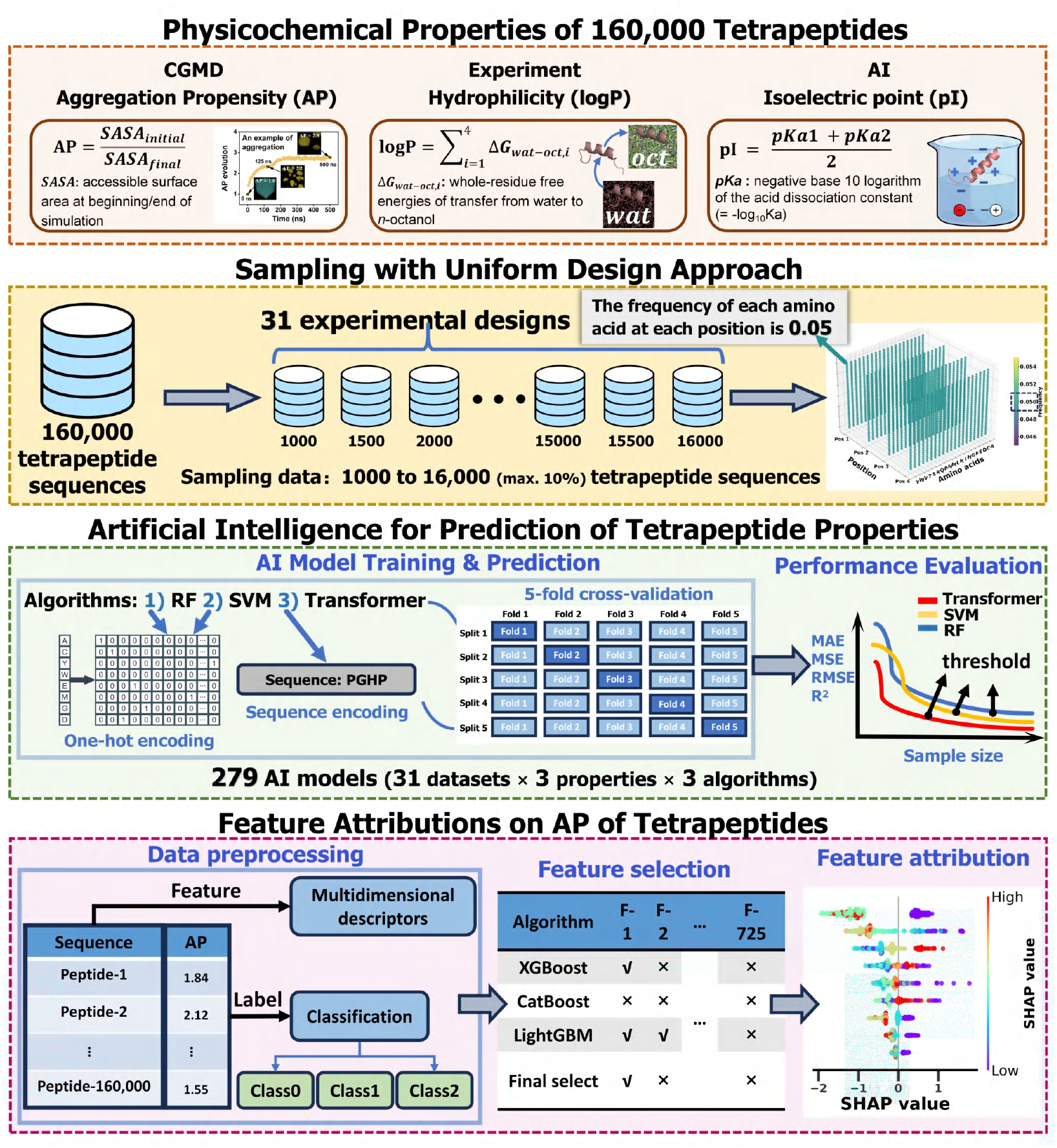
Workflow of integrated uniform design (UD) and artificial intelligence (AI) framework for predicting the physico-chemical properties of tetrapeptides. a) Generation of physicochemical properties for 160,000 tetrapeptides, including AP, logP, and pI, using CGMD simulations, experimental methods, and AI predictions, respectively. b) Sampling of training datasets using the UD approach, ensuring that the frequency of each amino acid at each position is 0.05. This process results in 31 experimental design datasets, with sample sizes incrementing by 500 from 1,000 to 16,000. c) AI model training to predict tetrapeptide properties (AP, logP, and pI) using RF, SVM, and Transformer models. For RF and SVM models, tetrapeptide sequences are preprocessed using one-hot encoding, while the Transformer model employs inputs of peptide sequences directly. Five-folds cross-validation is embedded during training. Model performance is evaluated using metrics of MAE, MSE, RMSE, and R^2^. Learning curve analysis is conducted to assess data utilization efficiency and identify performance inflection points. d) Feature attribution analysis for the AP of tetrapeptides. Multidimensional descriptors are extracted as features, and labels are classified into three categories to build interpretative models. SHAP analysis is adopted to elucidate the contribution of individual features to the predictive outcome, thereby enhancing model interpretability and identifying critical factors influencing tetrapeptide self-assembly behavior.

To gain deeper insights into the factors influencing the aggregation behavior of tetrapeptides, this study conducts an extensive analysis of the AP values of 160,000 tetrapeptide sequences. We employ tree-based machine learning (ML) models, including XGBoost^21^, LightGBM^22^, and CatBoost^23^, in conjunction with Shapley Additive Explanations (SHAP) analysis^24*−*26^ to identify the critical physicochemical characteristics and their interactions that affect the AP of tetrapeptides. The insights derived from these analyses may potentially be extrapolated to peptides of varying lengths, thereby enhancing the experimental design of self-assembling peptides across extended sequence space.

In summary, this research generates high-quality physiochemical property data for all 160,000 tetrapeptides (Supplementary Data 1) and develops accurate predictive models. Through the proposed integration of UD and AI, this work provides valuable insights into the relationship between physicochemical properties and aggregation behaviors. The selected tetrapeptide samples form a solid foundation for future AI-driven predictions of additional properties, such as peptide-protein binding affinities, enzymatic selectivity, and electronic band gap, etc., holding significant potential for applications in peptide drug developments, catalysts, and semiconductor technologies.

## 2. Results and Discussion

### 2.1 Distribution of UD Sampled Properties

To evaluate the effectiveness of the UD sampling method, we analyze the distribution patterns of AP, logP, and pI across the sampled sequences with varying sizes (1000, 4000, 8000, 12000, 16000) and compare them with the entire tetrapeptide sequence space (Figure 2). The AP values of the 160,000 tetrapeptides conform to a normal distribution with a mean of 1.86 (Figure 2a), while logP values exhibits an approximately normal distribution centered at -1.62 (Figure 2b). In contrast, the pI values display a more complex, multimodal distribution compared to AP and pI (Figure 2c). The sampled distributions of AP and logP are highly consistent with the overall distributions (Figures 2d-e). Although the distribution of pI values exhibits greater complexity, the sampled distribution increasingly aligns with the overall distribution as the sample size increases (Figure 2f). Further analyses show that the proportions of AP, logP, and pI intervals in the sampled data from 31 experimental design sets closely approximate those in the total dataset of 160,000 sequences (Figure 2g-i, Figure S1a-c).

**Figure 2.**
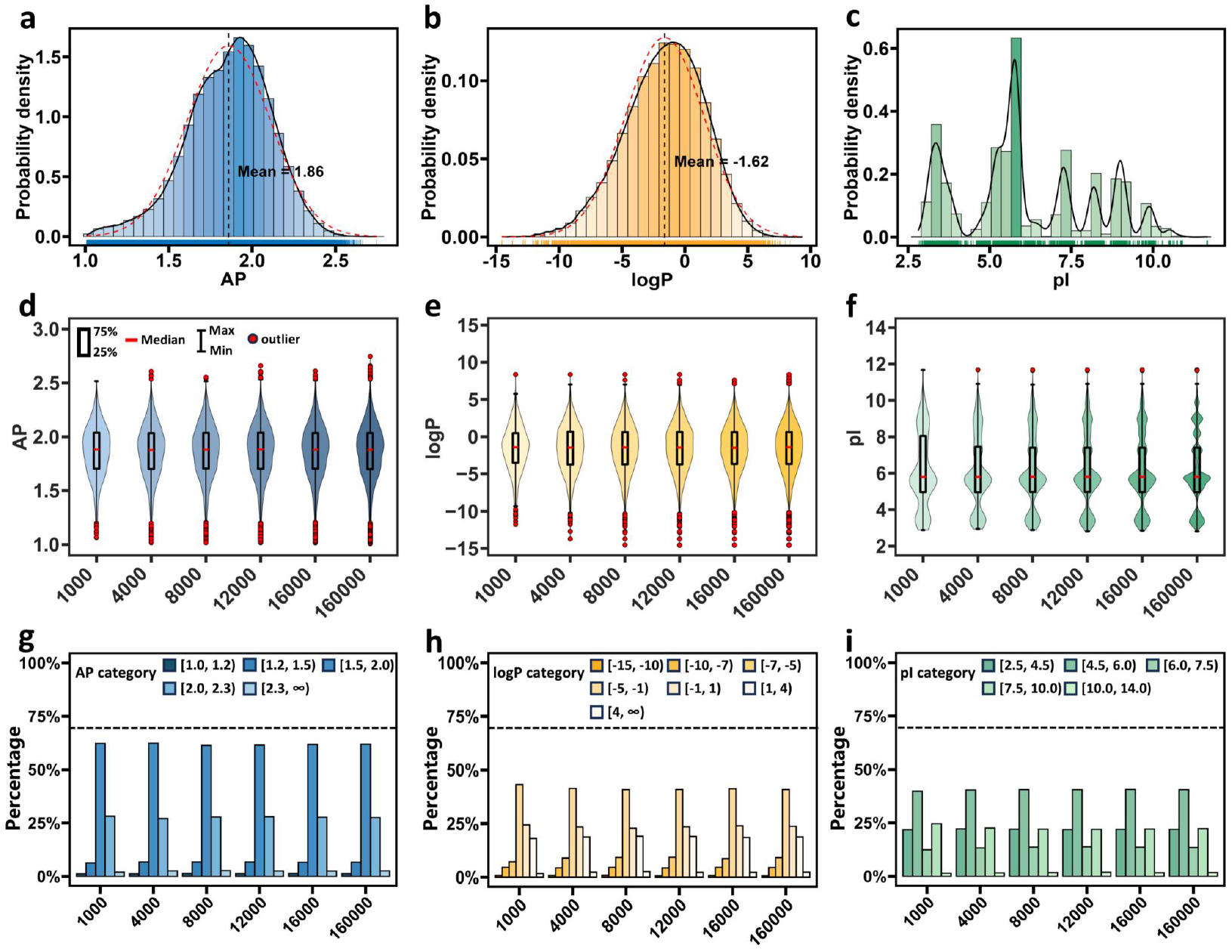
Comparison of distribution patterns of UD sampling data and overall data. a-c) Histograms and density plots of aggregation propensity (AP), hydrophobicity (logP), and isoelectric point (pI) of 160,000 tetrapeptide sequences. The red dashed lines represent the fitted normal distribution, while the black solid lines indicate the actual density distribution of the data. The color gradient in the histograms reflects density, with darker colors corresponding to higher density. The carpet plots at the bottom of each chart display the distribution of individual data points. d-f) Violin plots comparing the distribution of of AP, logP, and pI properties between UD samples (sizes of 1000, 4000, 8000, 12000, and 16000) with the overall population data. g-i) Proportions of sampled data relative to the overall dataset, after segmenting AP, logP, pI into defined intervals.

These comparisons demonstrate that the sampled data from various sizes can adequately represent the characteristics of the entire dataset. This finding underscores the ability of the UD method to mitigate sampling bias and guarantee comprehensive coverage of sequence diversity by maintaining a uniform frequency of 0.05 for each amino acid at each position (Figure S1d). This approach, in turn, lays a robust foundation for AI models to achieve superior accuracy and generalization in predicting tetrapeptide properties and beyond.

### 2.2 Performance Evaluations of AI models

The Transformer model consistently exhibits exceptional performance across all attributes (AP, logP, pI), as evaluated by metrics of MAE, MSE, RMSE, and R^2^ (Figure 3a-d). Results for both fixed and non-fixed testing datasets demonstrate that the Transformer model is more efficient at utilizing smaller training datasets effectively compared to the RF and SVM models. Notably, the Transformer model achieves significant performance (R^2^ = 0.91) gains with training datasets of approximately 3,500 to 4,500 data points (Table S1 and S2). Beyond this range, performance gains plateau, indicating that additional data contribute minimally to further improvement. Thus, the performance convergence threshold for the Transformer model in predicting tetrapeptide properties can be set at around 3,500 to 4,500 data points, corresponding to only 2.19% to 2.81% of the entire tetrapeptide sequence space.

**Figure 3.**
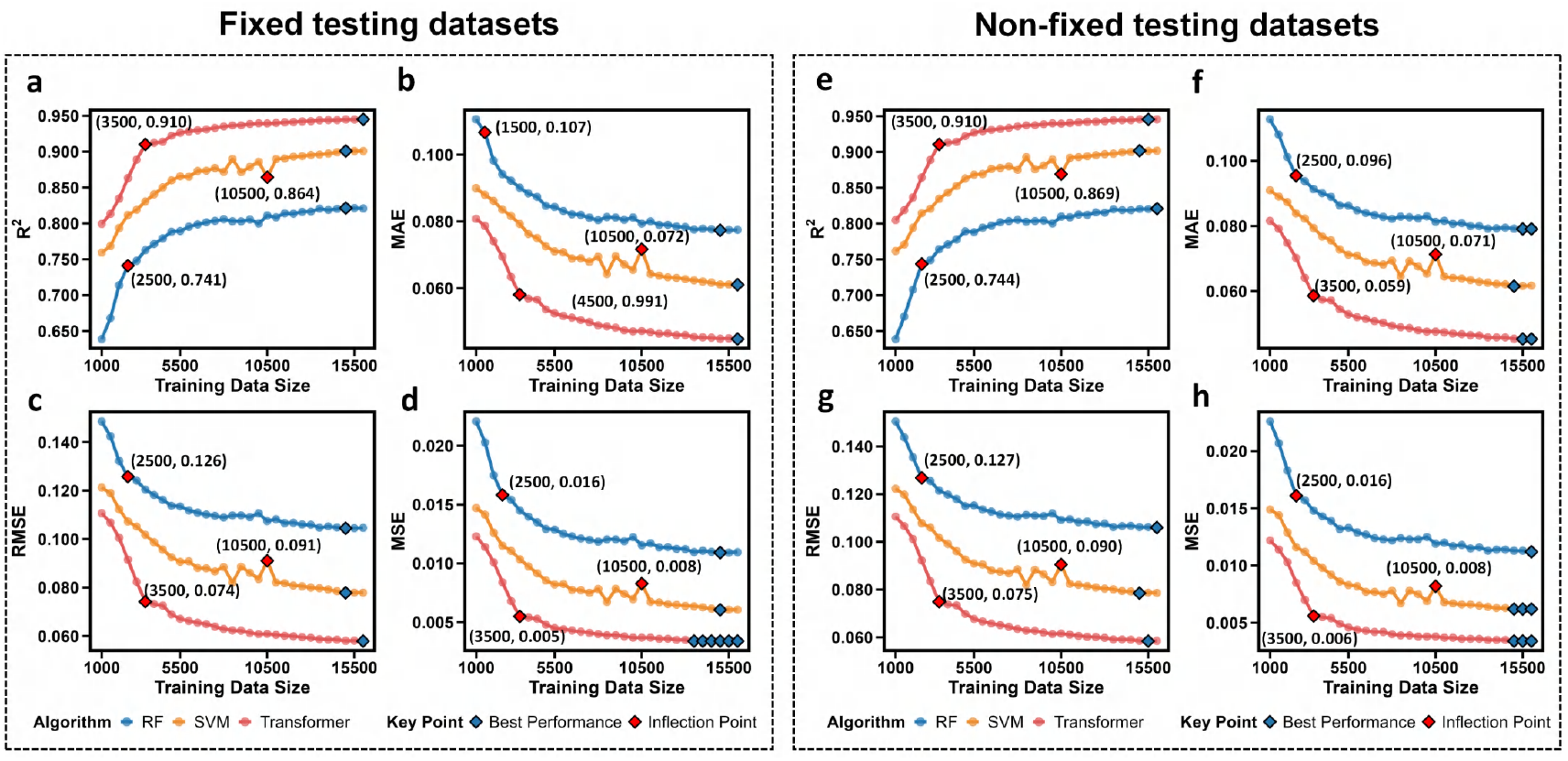
Learning curves of model performance. a-d) Performance metrics (MAE, MSE, RMSE, R^2^) of the RF, SVM, and Transformer models in predicting AP using a fixed testing dataset, evaluated across varying training set sizes. Blue diamonds mark the relative best performance points for each model, while red diamonds highlight inflection points where the performance trends change significantly. The inflection points are determined by the extreme value points of the second derivative, indicating significant curvature change in the learning curve. e-h) Performance metrics (MAE, MSE, RMSE, R^2^) of the RF, SVM, and Transformer models in predicting AP using a non-fixed test set, evaluated across varying training set sizes.

Conversely, the RF model reaches its performance inflection point with a smaller training dataset (about 2,000 to 2,500 data points) but exhibits relatively inferior performance compared to the Transformer model. Meanwhile, the SVM model slightly outperforms the Transformer in predicting the logP property (Figure S2) but underperforms in predicting the AP and pI (Figure 3 and Figure S3, respectively). These trends are consistent across both fixed and non-fixed testing sets. This synergy between the UD sampling strategy and the advanced Transformer model architecture, coupled with the accurate identification of performance inflection points, enables Transformer to demonstrate superior learning efficiency and predictive capability on medium-sized data sets.

### 2.3 Error Analysis of AI model

To elucidate the complex interplay among diverse error sources, sampling methods, tetrapeptide properties, and model architecture, we conduct a comprehensive error analysis (Figure 4) on the RF, SVM, and Transformer model trained with 4500 sequences, representing the convergence threshold for Transformer model performance. The model performance with different sizes, properties, and model type can be found in Figure S4 to Figure S12.

**Figure 4.**
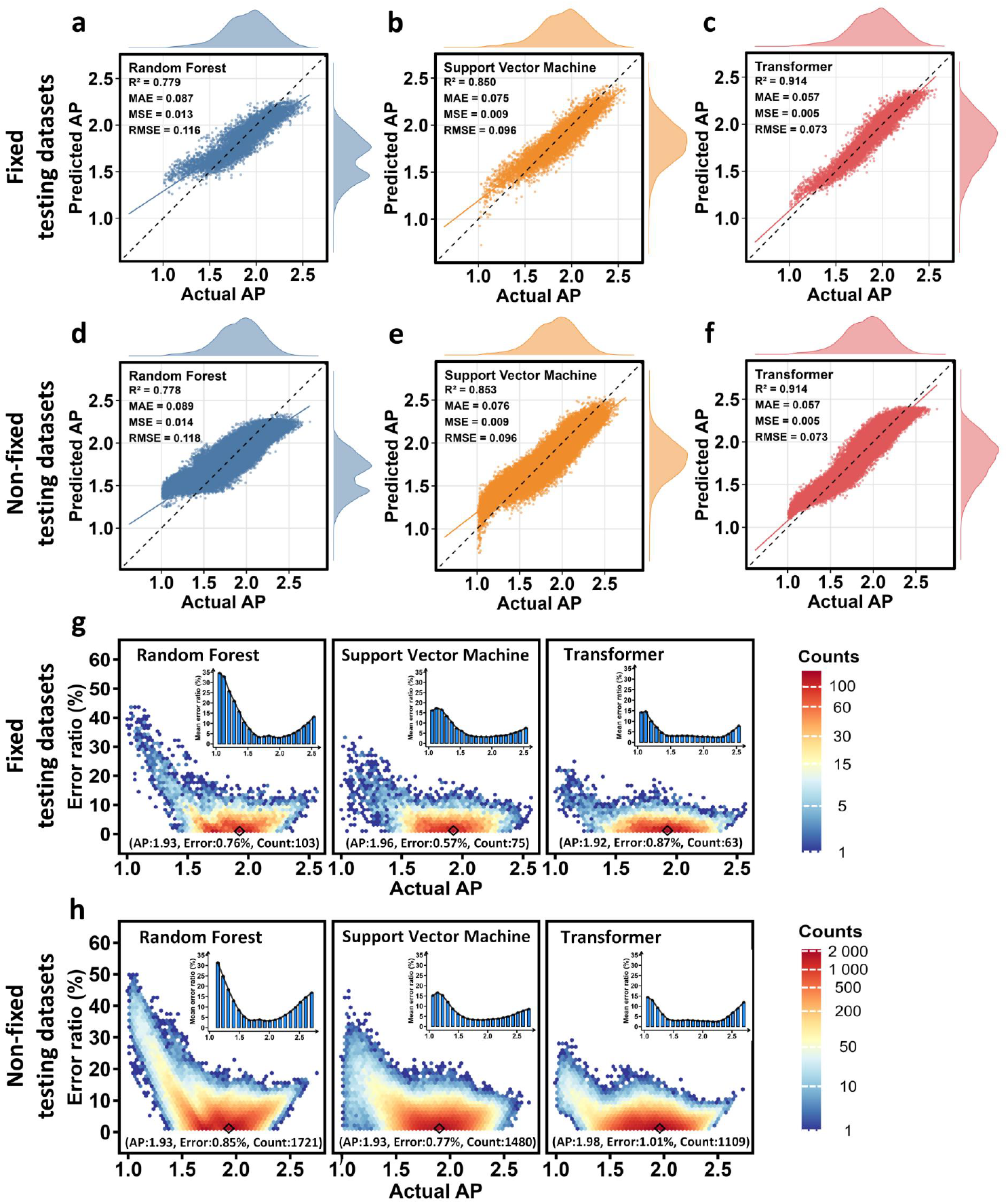
Cross-model prediction error analysis of Aggregation Propensity (AP). a-f) The relationship between the actual and predicted AP values of the RF, SVM, and Transformer models, evaluated using fixed and non-fixed testing dataset, with a training set of 4,500. Each plot includes a linear regression line (solid line) and an ideal prediction line (dashed line). Density plots are attached to the edges of the plots to illustrate the distribution of predicted and actual values. g-h) The relationship between actual AP values and corresponding prediction errors, visualized using hexagonal density plots for fixed and non-fixed testing dataset with the same training data. The prediction error is calculated as a percentage of the absolute difference between the actual and predicted AP value versus the actual AP values. The color gradient from blue to red indicates increasing point density, with the highest density points marked by black diamonds and corresponding coordinates labeled below. Histograms show the average prediction error rates for the RF, SVM, and Transformer models across the AP value range.

Regarding the AP, all models exhibit elevated error rates in the lower AP range (1 < AP < 1.5) and the high AP range (AP > 2.3, Figure 4a-f). This phenomenon may be attributed to the inherent scarcity of tetrapeptides at extreme AP values, which limits model learning in these intervals. In the intermediate range (1.5 < AP < 2), the error ratio stabilizes at around 2.5% (Figure 4g-h), while the SVM and RF models show marginally higher error ratios of 3% to 5% (Figure 4g-h). This stable error ratio suggests that the models can effectively capture the features of the tetrapeptide sequences in this region. This capability may be related to the higher density of samples in this intermediate range, where abundant data improve model performance and reduce both bias and variance in the errors.

In the prediction of logP, the model exhibits an unusually high error rate in the logP interval from -1 to 1 (Figure S13). This phenomenon is related to an inherent limitation of the error calculation method: when the logP value approaches zero, it acts as a denominator, thereby inflating the error proportion at extreme logP values (−15 < AP -10 and AP > 4, representing only 2.95% of the samples). Significant differences in model performance emerge in these sparse regimes, i.e., the RF model has an average error rate of 30-40%, while the Transformer has an average error rate of 10-20%, and the SVM model approaches an error rate of zero (Figure S13). This discrepancy may reflect differences in how each model handles sparse data regions, particularly when predicting extreme logP values, where SVM demonstrates unexpectedly high accuracy.

The pI predictions exhibit stable error ranges across interval of 5 to 9 (Figure S14), with mean error ratios generally below 10% for all models. However, in the low pI range (from 2.5 to 4.5), the RF and SVM models show higher error rates. The Transformer model also displays a marked increase in error within the range of 4 to 4.5 (Figure S14). These inconsistencies can be attributed to the multimodal complexity of the pI distribution and the varying abilities of the models to capture this complexity.

In summary, the Transformer model, leveraging its self-attention mechanism, adeptly captures long-term dependencies and intricate features, achieving minimal prediction error within data-dense intervals. Despite elevated errors in extreme intervals, the Transformer consistently outperforms SVM and RF models. While SVM excels at predicting logP values, it underperforms in predicting AP and pI. This disparity likely stems from the complex interplay between tetrapeptide properties and model features, where logP values exhibit more linear correlations with tetrapeptide sequences following one-hot encoding, favoring SVM, whereas AP and pI involve more complex nonlinear relationships that require the enhanced feature extraction abilities of the Transformer. By integrating UD sampling, multi-model comparisons, and detailed error analyses, this study reveals the intricate challenges in predicting tetrapeptide sequence properties, highlighting the importance of selecting appropriate algorithms based on specific properties and data characteristics. These findings provide valuable insights for future peptide property prediction studies and guide the optimization of sampling strategies and model architectures to enhance performance.

### 2.4 Feature Impact on Aggregation

In this study, three integrated learning tree models, i.e., XGBoost, LightGBM and CatBoost, combined with SHAP analyses are employed to systematically evaluate the key features influencing the aggregation of tetrapeptides. All tree models exhibit superior performance in multiclass classification (Table S3, Figure S15). We pay special attention to the features affecting the ideal AP range (class 1: 1.5 < AP < 2) for achieving self-assembly, with analyses focusing on their total effect (SHAP value), main effect (SHAP main value, SHAP_main_) and interactive effect (SHAP interactive effect, SHAP_inter_). The results of the SHAP analyses for other classes (class 0: 1 < AP < 1.5 and class 2: AP > 2) are presented in Figures S16 to S33.

#### 2.4.1 SHAP Total Effect

The assessment of model performance reveals that all three models exhibit high accuracy, demonstrating their robust discriminative capabilities in multi-category classification tasks. Through a thorough analysis of the SHAP dependence results from the three models, we identify four principal features via intersection analysis: aromaticity, pI, net charge, and logP. Although the specific ranking of these features varies across the models (Figure 5), there is unanimous agreement that aromaticity exerts the most significant impact on the AP of tetrapeptides, corroborating prior findings by Wang et al.^17^

**Figure 5.**
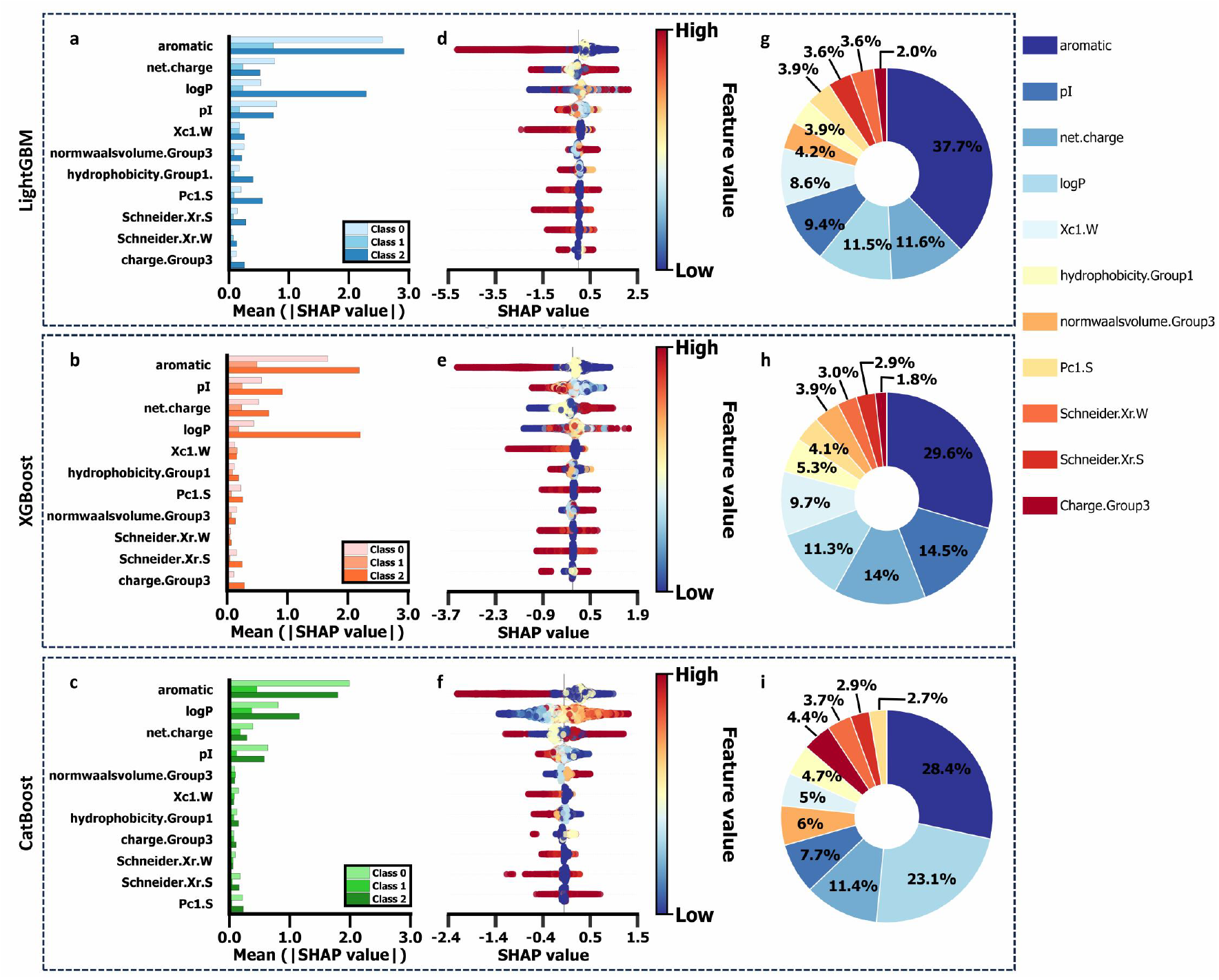
Cross-model prediction error analysis of Aggregation Propensity (AP). a-f) The relationship between the actual and predicted AP values of the RF, SVM, and Transformer models, evaluated using fixed and non-fixed testing dataset, with a training set of 4,500. Each plot includes a linear regression line (solid line) and an ideal prediction line (dashed line). Density plots are attached to the edges of the plots to illustrate the distribution of predicted and actual values. g-h) The relationship between actual AP values and corresponding prediction errors, visualized using hexagonal density plots for fixed and non-fixed testing dataset with the same training data. The prediction error is calculated as a percentage of the absolute difference between the actual and predicted AP value versus the actual AP values. The color gradient from blue to red indicates increasing point density, with the highest density points marked by black diamonds and corresponding coordinates labeled below. Histograms show the average prediction error rates for the RF, SVM, and Transformer models across the AP value range.

##### Aromaticity - total SHAP effect

To gain a deeper understanding of the impact of aromatic residue quantity on the aggregation of tetrapeptides, we first analyze the overall SHAP values (Figure 6, including both main effects and interactive effects). The results indicate that, for the formation of ideal aggregates (Class 1) for achieving self-assembly, the presence of 0 to 1 aromatic residue in the tetrapeptide contributes positively. However, incorporation of additional aromatic residues (beyond one) obstructs the formation of the optimal aggregation state (Figure 5d-f). To substantiate this threshold effect hypothesis, we conduct a Mann-Whitney U test across groups categorized by the quantity of aromatic residues (0-1, 1-2, 2-3, 3-4). The analysis shows that as the quantity of aromatic residues escalated from 1 to 2, the model’s predictive contribution for Class 1 undergoes a substantial change, with SHAP values declining markedly to negative values (XG-Boost: 156.64%, LightGBM: 220%, CatBoost: 128.14%; all p-values < 0.001). The differences remained significant after post-Bonferroni correction (adjusted p < 0.00025), robustly supporting the threshold effect hypothesis (Figure 6a-c).

**Figure 6.**
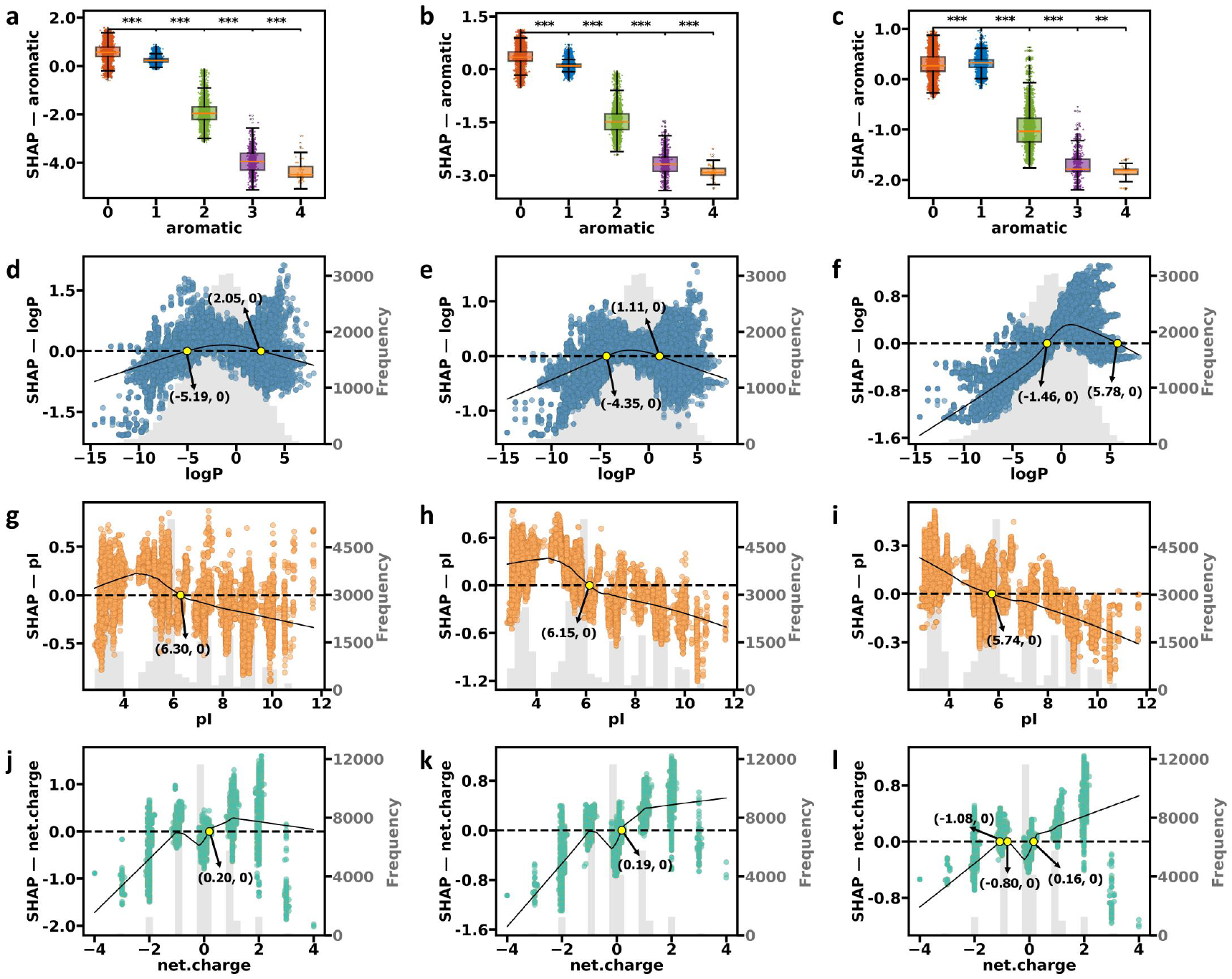
SHAP dependence plots for top features in Class 1 of aggregation. a-c) Impact of aromatic residue count on Class 1 SHAP values for LightGBM, XGBoost, and CatBoost models. Box plots show SHAP value distributions for different aromatic residue counts (0, 1, 2, 3, and 4). Boxes represent interquartile ranges, middle lines indicate medians, and whiskers extend to 1.5 times the interquartile range. Asterisks or NS (none-significance) above box plots indicate statistical significance levels of SHAP value differences between adjacent groups. Significance levels are based on Mann-Whitney U tests with Bonferroni correction: ^***^ p < 0.00025 (0.001/4), ^**^ p < 0.0025 (0.01/4), ^*^ p < 0.0125 (0.05/4), NS: p *≥* 0.0125 (not significant). d-f) SHAP dependence plots for the logP feature in Class 1 for LightGBM, XGBoost, and Cat-Boost models. Gray histograms (referring to the right axes) reflect the frequency distribution of logP values in the testing dataset. Black LOWESS (Locally Weighted Scatterplot Smoothing) fitting curves show the overall trend between logP and SHAP values. Black dashed lines indicate the SHAP value of 0 baselines, while yellow markers highlight critical values where the impact of logP on predictions shifts from positive to negative (or vice versa). g-i) SHAP dependence plots for the pI feature in Class 1 for LightGBM, XGBoost, and CatBoost models. These plots include scatter points, gray histograms, and black LOWESS curves, similar to panels d-f. Yellow points mark critical values where the impact of pI shifts. j-l) SHAP dependence plots for the net charge feature in Class 1 for LightGBM, XGBoost, and CatBoost models. These plots include scatter points, gray histograms, black LOWESS curves, and intersection annotations to comprehensively show how net charge affects model predictions.

##### logP - total SHAP effect

The effect of logP on the ideal aggregation state exhibits a distinct “window effect”. Specifically, moderate logP values facilitate class 1 outcomes, while both excessively high or low logP values are detrimental to the optimal aggregation of tetrapeptides (Figure 5, Figure 6d-f). Considering the results across the three models, LightGBM and XGBoost models provide more consistent intervals. Therefore, maintaining logP values between -5 and 2 is deemed most favorable for tetrapeptides to achieve desirable aggregation state (Figure 6d-e), while the CatBoost model suggests an interval of -1.5 to 6 (Figure 6d-e). The moderate hydrophilicity enhances intermolecular attraction between peptides in aqueous environment, prompting tetrapeptide molecules to form a hydrophobic core that promotes self-assembly. However, excessively low logP values may lead to solubility issues (i.e., precipitation), while extremely high logP values may result in solutions instead of self-assemblies.

##### pI - total SHAP effect

The relationship between the pI and AP of tetrapeptides highlights the delicate balance of charge in peptide aggregation processes. For an ideal self-assembling tetrapeptide, a lower pI value facilitates aggregate formation, exerting a beneficial effect, while a pI value approaching neutrality has minimal or negligible influence on aggregation. Conversely, a high pI value hinders the formation of Class 1 aggregates (Figure 5, Figure 6g–i). Examination of the SHAP dependence plots for pI in the LightGBM, XGBoost, and CatBoost models reveals that the inflection points are approximately situated at pI values of 6.3, 6.15, and 5.74, respectively. Below these inflection points, pI values positively influence aggregate formation, with the beneficial impact of charge interactions progressively accumulating and peaking around the inflection point. Beyond the inflection point, as pI continues to rise, its influence on the formation of ideal aggregates diminishes, while higher pI levels conversely impede the manifestation of Class 1 outcomes (Figure 6g–i). This behavior is likely linked to the dynamic alteration in the charge shielding effect: at low pI conditions (below 6.15), attractive forces between negative and positive charges are intensified, promoting the formation of stable aggregated structures. Conversely, at elevated pI values (above 6.3), the excessive concentration of negative or positive charges on the peptide chain results in solvation and neutralization of charge interactions, thereby impeding the formation of stable aggregates.

##### Net charge - total SHAP effect

Regarding net charge, high net charge values positively contribute to class 1 formation, while lower net charge values adversely affect it (Figure 5, Figure 6j-l). The SHAP dependence plot of net charge corroborates this pattern; the LOWESS smoothing curve indicates that as net charge value increases, its average SHAP effect on class 1 also rises. At a net charge value of approximately 0.20, all three models exhibit an inflection point where the SHAP value transitions from negative to positive, signifying that the positive contribution of net charge to Class 1. However, when the net charge value exceeds 3, almost all scatter points are clustered on the side with negative SHAP values, although the LOWESS curve remains positive (Figure 6j-l). This discrepancy may arise from sparse data points in this region, where local smoothing and boundary effects elevate the LOWESS curve above the actual SHAP values. The negative SHAP values at very high net charge values (greater than 3) suggest that local instability due to overcharging may occur under these extremely high charge conditions.

#### 2.4.2 SHAP Main Effect

##### Aromaticity - SHAP main effect

Upon further isolating the SHAPmain effect of the aromatic feature (Figure 7a-c), we discern a more precise influence pattern: the aromatic feature exhibits a positive impact on Class 1 prediction only in the absence of aromatic residues. However, the introduction of even a single aromatic residue results in a negative effect. This finding refines the prior threshold hypothesis, indicating that the influence of aromatic residues may be highly contingent upon the molecular environment, including the positional effects of aromatic residue sequences^27^, the characteristics of adjacent amino acid residues^28^, and the overall conformation of the peptide chain^29^. Notably, while the presence of aromatic residues obstructs the formation of Class 1 aggregates, it positively contributes to a stronger aggregation tendency associated with Class 2 (Figure S17a–c and Figure S19a–c). This phenomenon may arise from the substantial modification of the peptide’s physicochemical properties by aromatic residues through phi-phi stacking interactions, thereby influencing the self-assembly behavior.

**Figure 7.**
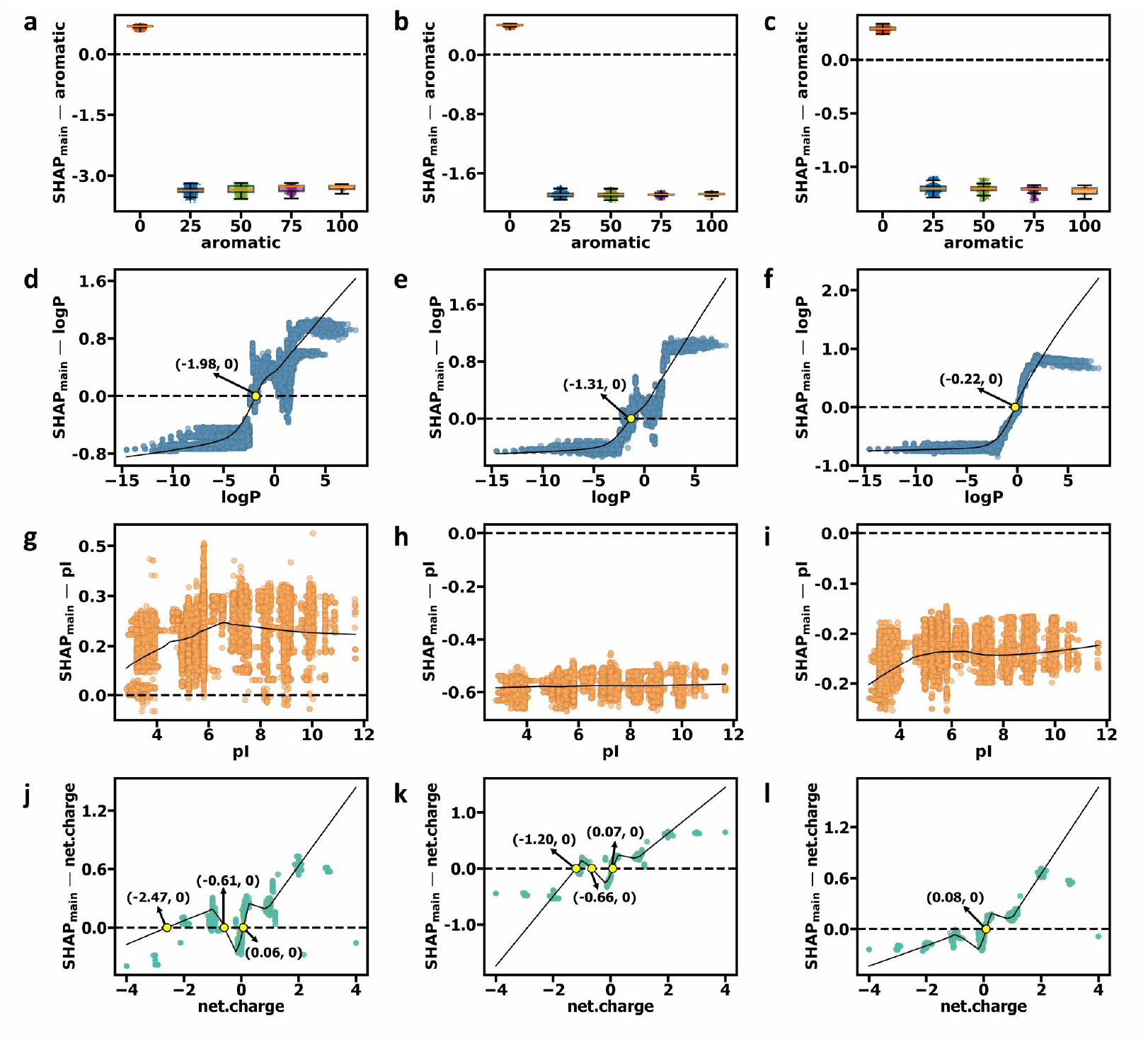
Main effect SHAP dependence plots for top features in Class 1 of AP. a-c) SHAP main effect dependence plots for the aromatic property on Class 1 results in the LightGBM, XGBoost, and CatBoost models (from left to right). d-f) SHAP main effect dependence plots for the logP feature on Class 1 results in the LightGBM, XGBoost, and CatBoost models (from left to right). g-i) SHAP main effect dependence plots for the pI feature on Class 1 results in the LightGBM, XGBoost, and CatBoost models (from left to right). j-l) SHAP main effect dependence plots for the net charge feature on Class 1 results. The black LOWESS (Locally Weighted Scatterplot Smoothing) curve in each plot depicts the overall trend between the feature values and their corresponding SHAP main effect values.

##### logP - SHAP main effect

Further analysis of the SHAPmain value dependence plot for logP in class 1 reveals that when logP is below the intersection points (−1.98 for LightGBM, -1.31 for XGBoost, -0.22 for CatBoost), the SHAP main effect value is negative. This implies that logP acts as an independent inhibitor to the model’s prediction for Class 1. Nevertheless, as logP levels increase within the favorable range of -2 to 2), a positive correlation emerges between logP and SHAP main effect values. Despite the SHAP main effect value remaining consistently positive, the total SHAP value of logP tends to be negative, with the LOWESS curve positioned below zero (Figure 7d-f). This indicates that the positive independent effect of logP is counterbalanced by a substantial negative interaction effect, resulting in an overall negative contribution of logP to class 1 prediction.

##### pI - SHAP main effect

Analysis of the SHAP main effect values for pI characteristics reveals substantial differences among the three models. The LightGBM model indicates that pI consistently provides a positive independent contribution to Class 1 prediction across all ranges, whereas the XGBoost and CatBoost models show a negative independent contribution in all ranges (Figure 7g–i). This disparity not only highlights the intricate mode of action that pI characteristics may possess but also uncovers the robust interactions that may occur between pI and other attributes.

This notable difference may arise from several factors. First, the pI feature may exhibit complex distribution characteristics, such as multi-modality, which could lead the models to adopt different strategies in interpreting its effect. Second, inherent variations in feature splitting and tree structure construction among the models might result in divergent assessments of the main effect of pI. Third, the strong interaction between pI and other features, such as net charge or logP, may be handled differently by each model during the separation of main effects. Despite these variations, the general consistency of the SHAP dependence plots provides significant insights: maintaining the pI value within the inflection point range (between 5.7 and 6.3) appears crucial for designing optimal self-assembling peptide sequences. Within this range, the peptide chain can achieve sufficient solubility while efficiently facilitating molecular interactions, thereby enhancing the formation of stable self-assembled structures.

##### Net charge - SHAP main effect

Regarding the SHAP main effect values for net charge, we observe that for most scatter points, the SHAP main effect values are above zero, further demonstrating the independent positive effect of high net charge on Class 1 (Figure 7j-l). The observed discrepancies indicate that the interactive effect between net charge and other features considerably influences the overall effect size, resulting in a negative impact on the total SHAP value. Thus, it is crucial to investigate how the interplay between net charge and other characteristics affects these patterns, especially within the high net charge spectrum.

#### 2.4.3 SHAP Interactive Effect

##### Aromatic - logP

To systematically quantify the impact of interaction effects among key features on the formation of desirable aggregation patterns in tetrapeptide sequences, we employ two complementary methods of SHAP interaction analysis: feature interaction heatmaps based on SHAP values and dependence plots of SHAP interaction effects. The heatmaps indicate that the most pronounced interaction effect is between aromatic residues and logP (interaction values of -0.12, -0.10, and -0.08 in the LightGBM, XGBoost, and CatBoost models, respectively), followed by the interaction between net charge and pI (interaction intensities of -0.05, -0.04, and -0.03), as illustrated in Figure 8a–c.

**Figure 8.**
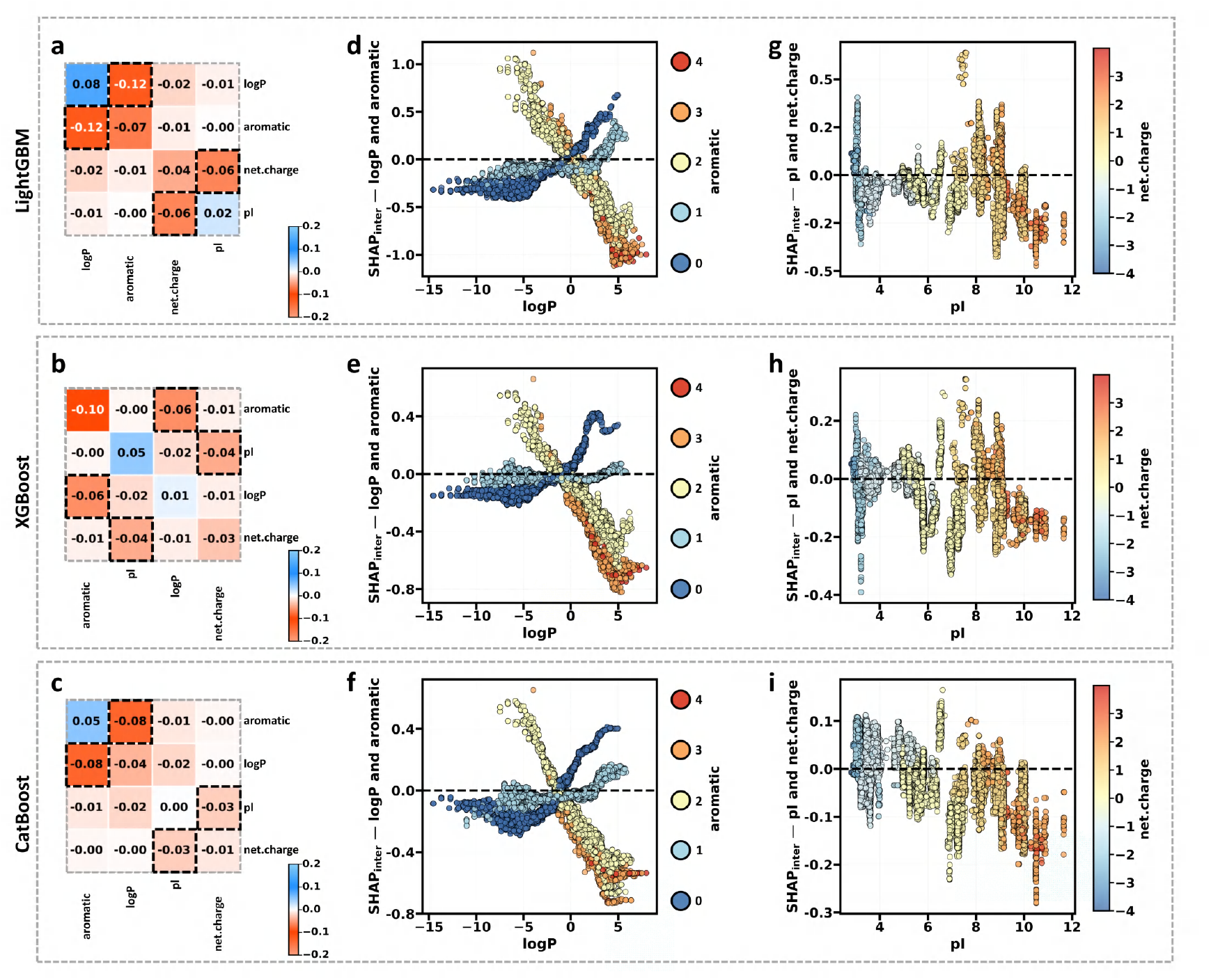
Interactive effect of SHAP plots for top features in Class 1 of AP. a-c) Heatmaps of SHAP interaction values for top features in Class 1 for LightGBM, XGBoost, and CatBoost models (from top to bottom). Off-diagonal elements quantify the magnitude and direction of pairwise feature interactions, while diagonal elements represent the strength and direction of individual feature main effects (positive in blue, negative in orange). These heatmaps depict interactions among the top 4 features, while the complete interaction heatmap of all 11 features provided in Supplementary Figure 20. d-f) SHAP interaction effect dependence plots for logP and aromatic features for LightGBM, XGBoost, and CatBoost models (from top to bottom). Black dashed lines indicate where SHAP effect values are zero. The color gradient illustrates how the interaction between logP and aromatic features affects model predictions, with vertical distribution reflecting the degree and direction of their influence. g-i) SHAP interaction effect dependence plots for pI and net charge features for LightGBM, XGBoost, and CatBoost models (from top to bottom). Color gradients illustrate shifts in interaction impacts, while the vertical distribution of points highlights the influence degree and direction of the interactions on SHAP values.

The dependence plots for the aromatic-logP interaction effect from the three models demonstrate that when logP is below -1, an increase in aromatic residues in tetrapeptides leads to a higher SHAP interaction effect. Specifically, the effect ranges from -0.5 to 0 for sequences with no aromatic residues, -0.2 to 0 for those with one aromatic residue, and 0 to 0.5 for those with two or more aromatic residues. This interaction effect diminishes as logP approaches -1, becoming negligible around -1 regardless of the number of aromatic residues (Figure 8d–f).

Conversely, when logP exceeds -1, the interaction effect is inverted. For tetrapeptides without aromatic residues, the interaction effect rises significantly with increasing logP. In contrast, the interaction effect increases more gradually for those with one aromatic residue. For tetrapeptides with two or more aromatic residues, the interaction effect is negative and decreases significantly with higher logP. This indicates that at low logP, more aromatic residues enhance aggregation, whereas at high logP, fewer aromatic residues are favorable for aggregate formation, and an excess may destabilize the aggregate structure.

##### Net charge - pI

The dependence plot of the net charge-pI interaction effect reveals a distinct color gradient trend, indicating significant variation in net charge values across different pI intervals (Figure 8g–i). This trend aligns with the established relationship between the magnitude of the pI and charge distribution^30*−*32^. Specifically, increasing pI values correlate with a progressive transition in net charge from negative to positive.

Within the pI range below 6, the system’s charge distribution is predominantly negative, resulting in negative interaction effects. This suggests that in acidic environments, negative charges impede the development of Class 1 aggregates. As the pI value increases from 6 to 8, the charge state progressively approaches neutrality, leading to an equilibrium in charge distribution and a gradual shift toward positive net charge. During this transition, the interaction effect between net charge and pI exhibits no substantial positive or negative impact.

In the elevated pI range above 8, the solution becomes increasingly alkaline, resulting in less protonation of acidic amino acid side chains and a transition toward positive charge. The plot manifests as orange and red hues; however, the interaction impact remains negative. This indicates that the interplay between high pI and high net charge may result in an uneven charge distribution, thereby influencing the stability of the aggregation morphology.

##### Interactions among other features

Our analysis indicates that, alongside primary features, secondary variables subtly influence the aggregation tendency of tetrapeptides. Variables such as Pc1.S (amphiphilic pseudo-amino acid composition for serine) and normwaalsvolume.Group3 (Normalized van der Waals Volume Group 3), though individually minor in impact, offer supplementary dimensions for refining aggregation behavior. While the individual effects of these traits are small, their combined influence may be pivotal in accurately regulating the aggregation process, particularly when optimizing self-assembly behavior to achieve specific functions.

These findings underscore the necessity of evaluating a comprehensive array of structural and physicochemical features when creating and optimizing peptide-based materials to achieve precise control over aggregation behavior.

In summary, we have established a comprehensive framework for studying the complex feature interaction network influencing the self-assembly behavior of tetrapeptides by combining multi-model ensemble methods with SHAP explanations. This methodology enhances understanding of existing data and guides the future design of specialized self-assembling peptide materials with desirable functions.

## 3. Conclusion

This study investigates various physicochemical properties of the tetrapeptide sequence space using UD sampling approach, generating a high-quality, low-bias dataset for AI model training. The sampling technique maintains an equal probability distribution (5%) for each amino acid at every position, minimizing bias in amino acid type and position. Statistical analyses confirm that UD sampling produces representative subsets highly consistent with the theoretical sequence space in terms of AP, logP, and pI distributions.

In model performance evaluations, combining UD sampling with the Transformer architecture demonstrates significant advantages and broad applicability. The technique achieves a performance inflection point at reduced training set sizes (3500 – 4500 sequences) for predicting AP, logP, and pI, highlighting the framework’s efficiency in predicting peptide properties with limited data. This reduces data acquisition and computational resource requirements while maintaining high accuracy. Detailed error analysis reveals challenges in predicting extreme values and highlights the need for enhanced sampling density or targeted tuning techniques in these regions. These findings emphasize the importance of considering data attributes, problem complexity, computational efficiency, and model capability when selecting and optimizing predictive models.

Additionally, SHAP analysis identifies key factors influencing tetrapeptide aggregation, including the threshold effect of aromaticity, the window effect of hydrophobicity, and complex interactions among features. These insights provide valuable guidance for designing self-assembling tetrapeptides.

In summary, this work presents a comprehensive framework for examining and predicting tetrapeptide characteristics. It enhances understanding of the molecular principles governing self-assembly and offers quantitative insights for designing peptides with customized aggregation properties. These findings advance the discovery of novel biomolecules and address challenges in biomedicine and materials science.

## 4. Experimental Section

### Calculations of physicochemical properties

The AP values of tetrapeptides are calculated based on the results of CGMD simulations, performed using the open-source package GROMACS^33^ and the Martini force field (version 2.2)^34*−*36^. The formula for AP is defined as:

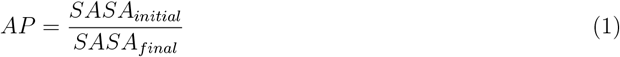

Here, SASA_*initial*_ and SASA_*final*_ denote the solvent-accessible surface area of all tetrapeptides in the simulation box at the beginning and end of the CGMD simulation, respectively^37^.For non-aggregating tetrapeptides, the AP value remains at 1. According to prior research^17,38^, an AP value ranging from 1.5 to 2 signifies favorable aggregation, facilitating optimal self-assembly behavior. However, when the AP value exceeds 2, the tetrapeptides may exhibit excessive aggregation, resulting in precipitation.

The logP value of peptides reflects hydrophilicity, indicating their tendency to partition between octanol and water. Hydrophilic peptides (larger logP values) readily interact with water molecules through hydrogen bonding, whereas hydrophobic peptides (smaller logP values) tend to avoid such interactions^39,40^. The hydrophilicities of 160,000 tetrapeptide sequences are experimentally determined and calculated using the following equation:

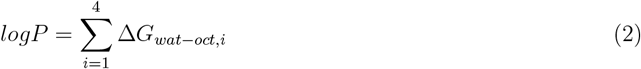

In this equation, Δ*G*_*wat−oct,i*_ (unit: kcal/mol) represents the Wimley-White whole-residue hydrophilicity of each amino acid^41^, corresponding to the free energy change associated with transferring an amino acid from the aqueous phase to *n*-octanol phase.

If Δ*G*_*wat−oct,i*_ > 0, the amino acid is more stable in the aqueous phase than in the n-octanol phase, indicating hydrophilicity and favoring interactions with water molecules. Conversely, if Δ*G*_*wat−oct,i*_ < 0, the amino acid is hydrophobic and prefers to avoid interactions with water.

The pI^42^ of a peptide refers to the pH value at which the net charge of the peptide molecule is zero, resulting from the balance between positively charged basic amino acids and negatively charged acidic amino acids. This value is calculated using an online Isoelectric Point Calculator 2.0 (IPC 2.0)^43^, which utilizes DL to predict the pI and pKa values of proteins and peptides. This tool can be accessed at http://www.ipc2-isoelectric-point.org.

### Sampling with uniform design

In this study, we sample subsets within the tetrapeptide sequence space using the UD method^44^. This approach is a type of space-filling design that aims to select experimental points that are uniformly distributed over the entire experimental domain^45^. Our experimental domain, *χ*, is defined as the Cartesian product of four factors (A, B, C, D), with each factor representing a position in a tetrapeptide sequence at the level of 20 natural amino acids. Specifically, *χ*={*A*_1_, *A*_2_, …*A*_20_} × {*B*_1_, *B*_2_, …*B*_20_} × {*C*_1_, *C*_2_, …*C*_20_} × {*D*_1_, *D*_2_, …*D*_20_}, resulting in a sequence space of 160,000 possible combinations of tetrapeptides.

In this space, each combination of factor levels (4 factors, 20 levels each) is termed a level-combination, which can be regarded as a point in the experimental domain. For example, the sequence KEAD represents a level-combination (K, E, A, D). We use the symbol “*n*” to denote the number of experimental runs, with each run producing a level-combination, i.e., a tetrapeptide sequence. A set of experimental designs can be represented as a matrix *U* = *x*_1_, *x*_2_, …, *x*_*n*_, where *x*_*j*_ ∈ *χ* and *n* equals to 1000 to 16,000 with an increment of 500.

To facilitate efficient sampling across this extensive sequence space, we employ the GenUD function of the R package UniDOE^46^ (Supplementary Code 1), which utilizes a Stochastic Optimization Adaptive Threshold Acceptance (SOAT, Figure 9) approach for the effective generation of uniform designs. Unlike traditional Threshold Acceptance (TA) algorithms^45,47^ that rely on fixed threshold sequences, the SOAT approach adapts dynamically to the search process by automatically modifying the thresholds. This flexibility enables the algorithm to escape local optima and identify the optimal design within a given design space more rapidly, resulting in a design with minimal deviation from the uniform distribution.

**Figure 9.**
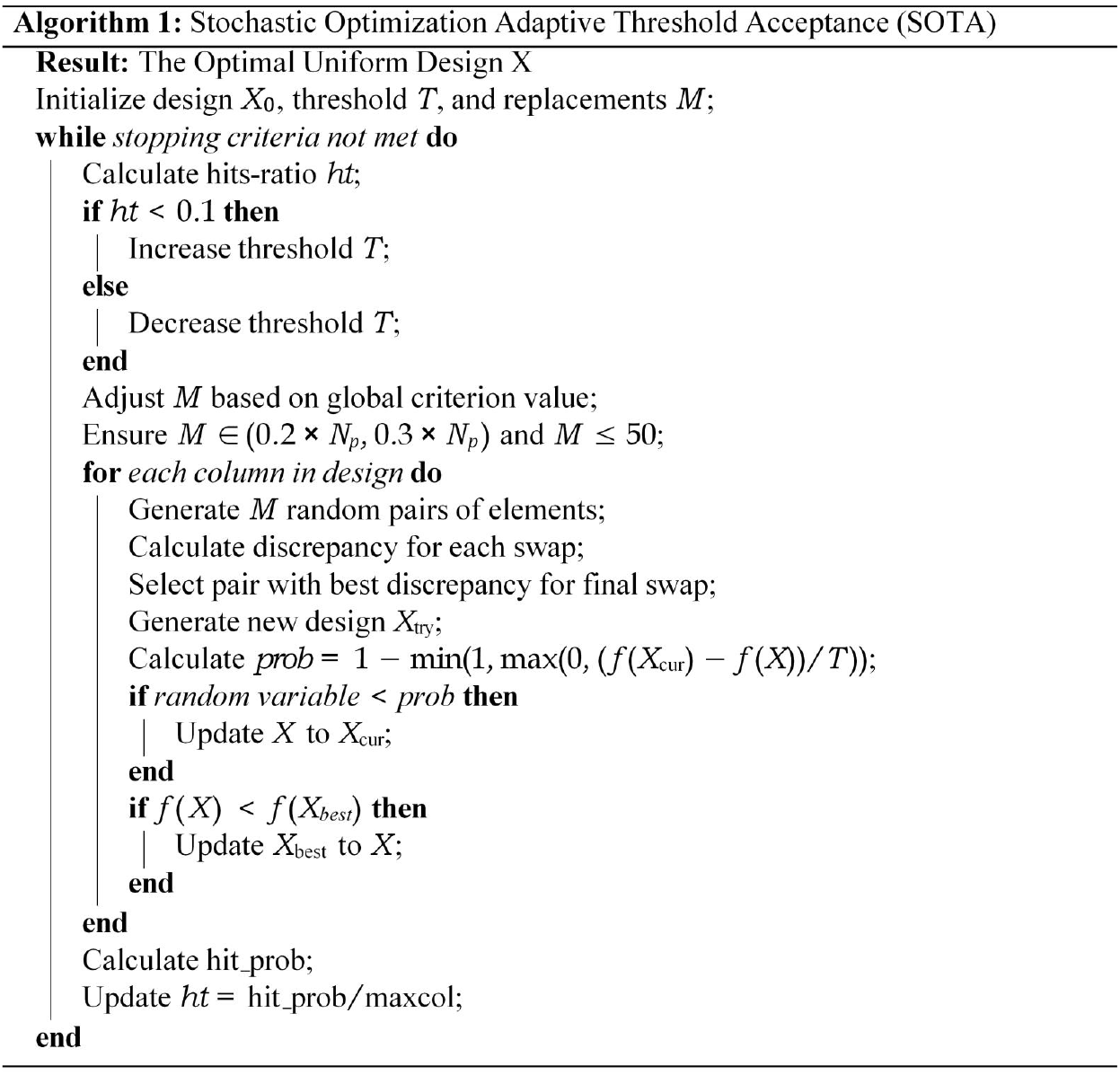
Algorithm of the Stochastic Optimization Adaptive Threshold Acceptance (SOAT) technique.

An mixture L2-Discrepancy (MD2) is established as the optimization objective to enhance the design quality within each design set U. The MD2 criterion, proposed by Zhou et al.^48^, integrates the benefits of the Centralized L2-Discrepancy (CD2) and the Wrap-around L2-Discrepancy (WD2)^49,50^. It also satisfies the evaluation criteria established by Fang et al.^51^ for eight uniformity measures employed in the assessment of experimental design construction. MD2 is particularly appropriate for our amino acid sequence space, as it preserves the uniformity of design points while accounting for the periodic boundary requirements of the experimental domain. The equation for MD2 is as follows:

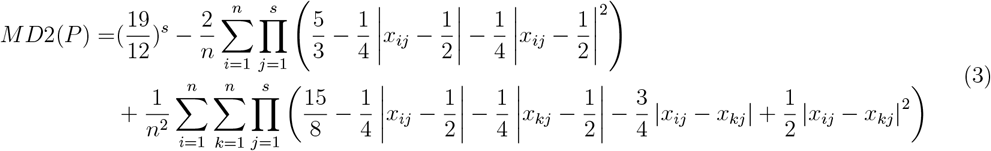

Where *n* is the number of experimental runs, *s* (= 4) is the factor of the experiment.

Subsequently, we optimize MD2 using the SOAT technique, as illustrated in Figure 9. The algorithm consists of an internal for-loop and an external while-loop, which work in tandem to identify the optimal design by continuously adjusting the threshold and swapping design points. The fundamental concept of this method is to globally seek the optimal design by probabilistically accepting inferior designs to escape local optima while preserving design uniformity.

This sampling technique maximizes the coverage of the design space with a minimal number of samples, significantly reducing the number of peptide sequences to be evaluated while ensuring the representativeness and uniformity of the selected sequences. This method markedly enhances the efficiency of exploring the tetrapeptide sequence space and yields high-quality training data for the subsequent prediction of physicochemical properties using AI.

### Training and testing of AI models

This study employs two classical ML algorithms, i.e., RF and SVM, and one DL algorithm, i.e., Transformer, to predict three properties of tetrapeptides, namely AP, logP, and pI (Supplementary Code 2).

In the data pre-processing phase for the RF and SVM models, we apply one-hot encoding to transform the tetrapeptide sequences into 80-dimensional numerical vectors. To enhance model performance, we employ a random search approach to randomly select 50 parameter combinations within the specified hyperparameter space and conduct a 5-fold cross-validation on the training dataset. The optimal hyperparameter configurations are ultimately selected based on the Root Mean Square Error (RMSE) of the validation set (see Supplementary Table 4).

Regarding the Transformer model, we modify the open-source code available at https://github.com/Zihan-Liu-00/DL-for-Peptide. Specifically, we retain only the essential code relevant to the regression task of the Transformer model and refine it further for tetrapeptide property prediction. We use RMSE other than MSE as the optimization objective to enhance sensitivity to larger errors. To address potential instability and convergence challenges associated with varying dataset sizes (1,000 to 16,000 sequences), we implement a linear warm-up learning rate strategy. Training begins with an initial learning rate of 0.001, which gradually increases to 0.2 over the designated warm-up phases, facilitating steady convergence across different data scales. Additionally, we employ a stochastic gradient descent (SGD) optimizer^52^ with a momentum of 0.9 to enhance convergence speed and minimize oscillations during training. Collectively, these optimization techniques improve the adaptability and stability of the Transformer model for tetrapeptide property prediction tasks, enabling effective learning across various dataset scales and properties.

During training, 80% of the data is allocated for model training, while the remaining 20% is reserved for model validation. The training datasets are deposited as Supplementary Data 2. To thoroughly assess the generalization capability and predictive stability of the models, we implement two distinct testing set selection strategies following the identification of optimal hyperparameters for each model: fixed testing datasets (Supplementary Data 3) and non-fixed testing datasets (Supplementary Data 4). In the fixed testing dataset framework, 10,000 tetrapeptide sequences, absent from the training data, are randomly selected from the entire tetrapeptide sequence space. In the non-fixed testing dataset framework, the residual data (144,000 to 159,000 sequences, depending on the number of training data) within the complete sequence space serve as the testing set for the respective model. A total of 279 AI models [31 training datasets (Supplementary Data 2) × 3 properties × 3 algorithms] are trained and subject to performance assessment. Model performance is quantitatively assessed using multidimensional metrics^53^, including Mean Absolute Error (MAE), RMSE and MSE, and the Coefficient of Determination (R^2^). The full performance metrics of the models on the training, validation, and testing sets are detailed in Supplementary Data 5 to 9.

### Tree-based model

This work employs three integrated tree-based models, XGBoost, LightGBM, and CatBoost, to investigate the principal variables influencing the AP values of tetrapeptides. These models belong to the Gradient Boosting Decision Tree (GBDT) family^54,55^, which offers robust nonlinear modeling capabilities for efficiently handling large datasets, making them well-suited for analyzing the full space of tetrapeptide sequences. XGBoost regulates model complexity through regularization, effectively mitigating overfitting. It enhances gradient and error updates by employing second-order Taylor expansion, thereby substantially increasing training speed and model accuracy22. LightGBM adopts a histogram-based decision tree construction technique that integrates leaf-wise growth with gradient-based one-sided sampling (GOSS). This approach markedly improves processing efficiency, particularly on large, high-dimensional datasets^22^. CatBoost effectively mitigates overfitting due to its symmetric tree architecture and distinctive feature combination approach. These characteristics make it especially advantageous for scenarios with suboptimal data quality or noise^23,56^.

We categorize the AP values into three classes based on a detailed understanding of peptide self-assembly behavior: Class 0 (AP *∈* [1, 1.5)), Class 1 (AP *∈* [1.5, 2)), and Class 2 (AP *∈* [2, 2.5]). Class 1 represents the optimal aggregation state, exhibiting the most promising characteristics for achieving self-assembly.

For feature extraction, we adopt the R package protr (version 1.7-4)^57^ to convert tetrapeptide sequences into multidimensional descriptors, encompassing the physicochemical attributes and structural characteristics of the tetrapeptides. Additionally, the R package peptides (version 2.4.6)^58^ extracts supplementary properties, including net charge, hydrophobicity, Boman’s index, and the proportion of each amino acid type, thereby enriching the descriptor set. Experimentally determined logP values and AI-predicted pI values are also incorporated into the feature set. The final integration of 726 descriptors provides a comprehensive array of input features for training tree-based ML models (Supplementary Code 3, Supplementary Data 10).

A multi-model voting strategy is used to guarantee the selection of the most significant features. The average importance score for each feature is calculated using the built-in feature importance scoring functions of the XGBoost, LightGBM, and CatBoost models, in conjunction with five-fold cross-validation.

A dynamic threshold (mean importance plus one standard deviation) is established to identify features deemed significant in each model. Ultimately, features that are recognized as significant by at least two models are selected, ensuring consistency and relevance across models. This strategy substantially reduces the feature set and eliminates redundancies.

To address the data imbalance problem, we implement a stratified sampling technique during the data preprocessing phase to ensure a balanced distribution of target variable categories. We divide the entire dataset of 160,000 tetrapeptide sequences into an 80% training set and a 20% testing set. During the hyperparameter tuning phase, we conduct a random search over 50 hyperparameter combinations within the hyperparameter space and assess the performance of each combination using stratified five-fold crossvalidation. The optimal hyperparameter settings are selected based on negative log loss metric to enhance the models’ robustness against imbalanced data. The hyperparameter search space for each model is outlined in Table S5. To thoroughly evaluate model performance, we compute six metrics: accuracy, precision, recall, F1 score, ROC AUC, and PR AUC. Additionally, we generate confusion matrices, ROC curves, and PR curves to visualize the classification efficacy of the models^59,60^, and the prediction results for tree models are shown in Supplementary Data 11.

### SHAP analysis

To elucidate the complex mechanisms that govern the self-assembly of tetrapeptides, we implement SHAP analysis^25^ (Supplementary Code 3), a sophisticated approach rooted in game theory. This approach provides an in-depth evaluation of model predictions and elucidates the diverse influences of physicochemical properties and structural characteristics on AP. By employing SHAP analysis, we identify critical features and gain a comprehensive understanding of their influence on model predictions, including their overall impact, main effects, and interaction effects.

The SHAP framework proposes an additive decomposition of a model’s prediction *f* (*x*) for any instance, succinctly expressed as: *f* (*x*) = *φ*_0_ + ∑_*i*_ *φ*_*i*_ + ∑_*i<j*_ *φ*_*ij*_, where *φ*_0_ represents the base value, corresponding to the model’s prediction in the absence of any features. The terms *φ*_*i*_ denote main effect values, reflecting the average contribution of each feature *i* across various feature backgrounds, while the terms *φ*_*ij*_ represent interaction effect values, capturing the pure synergistic impact of features *i* and *j*, distinct from their individual contributions. This decomposition provides a comprehensive perspective on feature impacts, encompassing both individual effects and complex interactions. Through this rigorous SHAP analysis, we bridge the gap between complex ML predictions and interpretable scientific insights, providing rational design strategies in peptide-based nanomaterials and advancing our comprehension of the fundamental rules governing molecular self-assembly.

## Supporting information

List of Abbreviations and Acronyms

## Supporting Information

Supporting Information including the supplementary data, code, results are available at the Github repository (https://github.com/JiaqiBenWang/UD-AI-Peptide) or from the author. The supporting information and the supplementary file of “List of Abbreviations and Acronyms” are available from the Wiley online library.

## Acknowledgements

Dr. Jiaqi Wang acknowledges the funding support from the National Natural Science Foundation of China (Grant No. 52101023), the Basic Research Program of Jiangsu (Grant No. BK20241816), and the Research Development Fund (RDF) of Xi’an Jiaotong-Liverpool University (RDF-24-01-013). Dr. Haojin Zhou acknowledges the funding support from the Research Development Fund (RDF) of Xi’an Jiaotong-Liverpool University (RDF-23-01-073). The authors also acknowledge the support of the high-performance computing platform at Xi’an Jiaotong-Liverpool University.

## Table of Contents

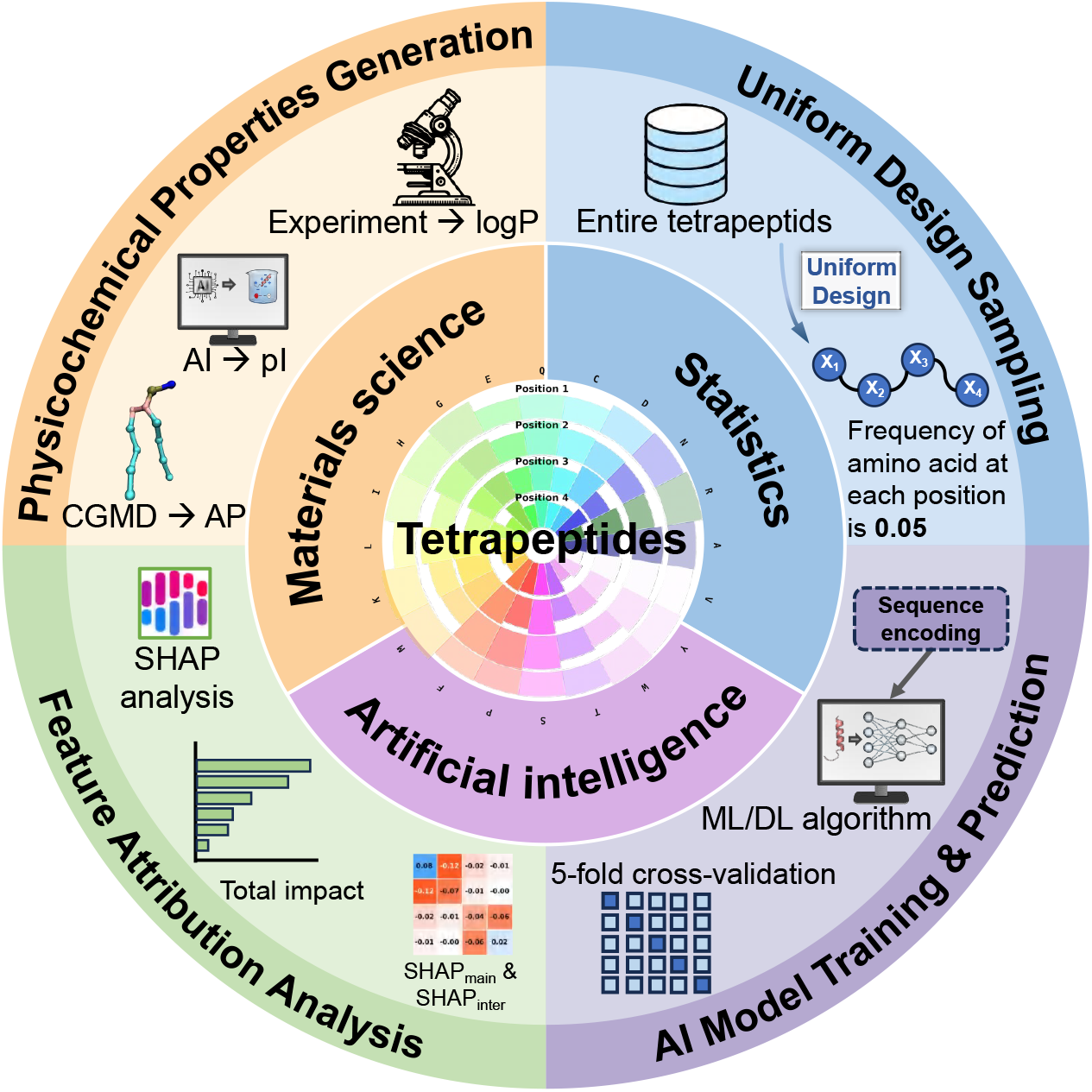

ToC Entry

