## Supplemental Information for "Harnessing Uniform Design to Enhance AI-Driven Predictions of Physicochemical Properties of Short Peptides"

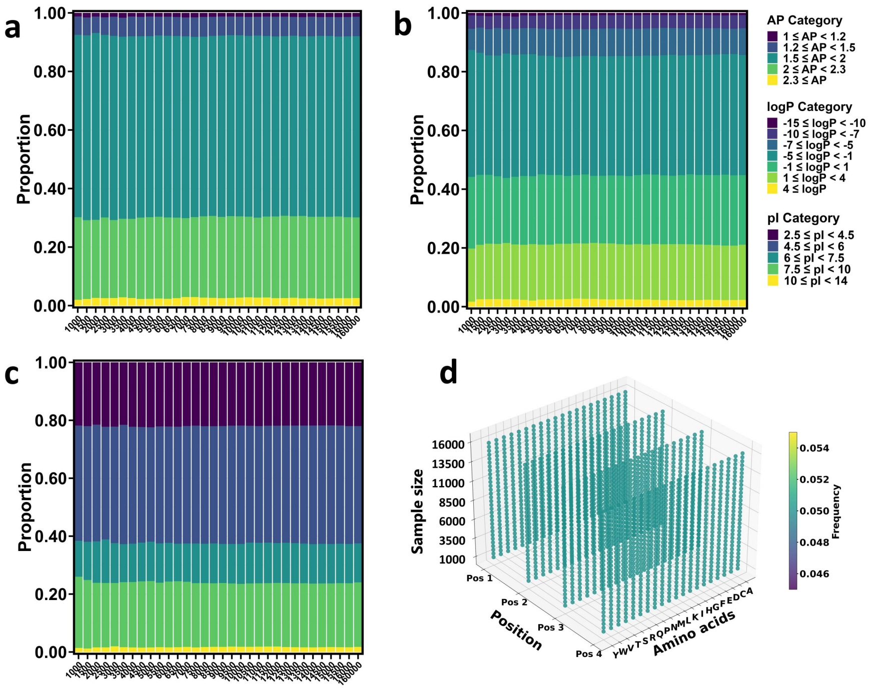


**Figure S1. a-c)** UD sampling of 1000 to 16000 samples (with each increase of 500 data) compared to the population data at different intervals of AP, logP, and pI attributes. **d)** The frequency of 20 amino acids at different positions in the tetrapeptide sequence for 1000 to 16000 tetrapeptide sequence data (increasing 500 at a time) under different sample numbers.

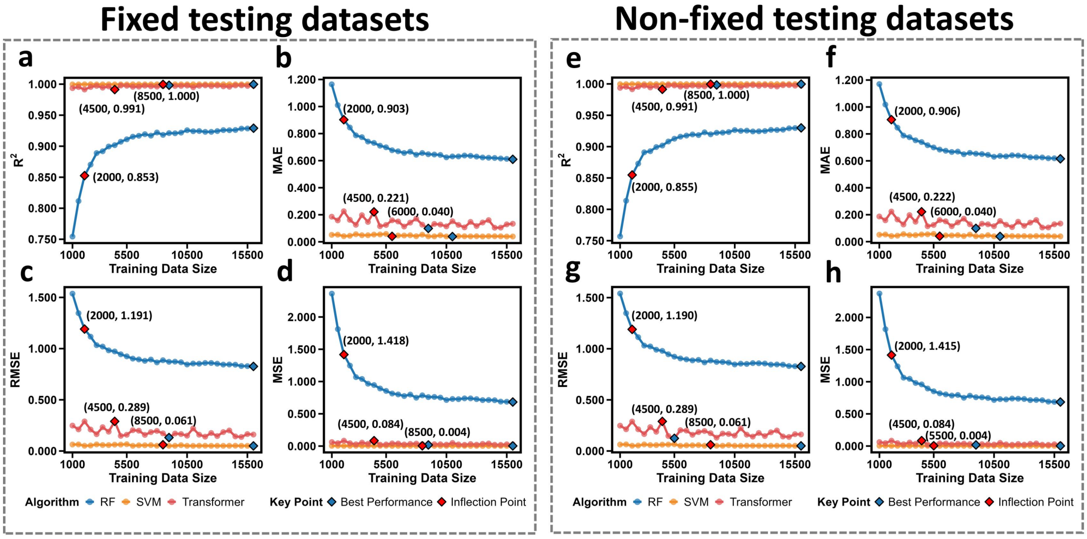


**Figure S2. a-d)** Performance metrics (MAE, MSE, RMSE, $R^2$) of the RF, SVM, and Transformer models in predicting the logP using a fixed test set, evaluated on models trained with varying training set sizes. **e-h)** Performance metrics (MAE, MSE, RMSE, $R^2$) of the RF, SVM, and Transformer models in predicting logP using a non-fixed test set, evaluated on models trained with different training set sizes.

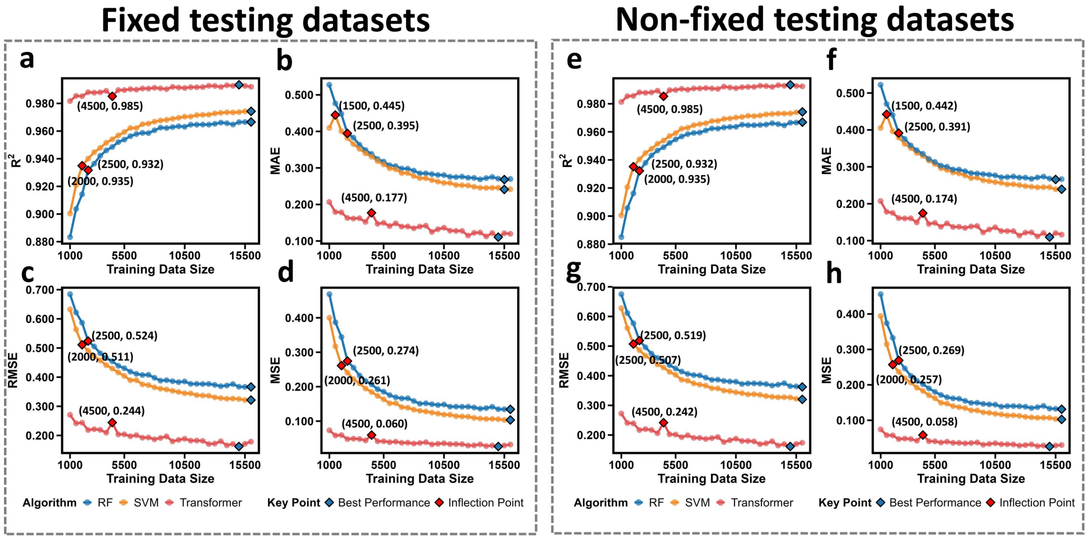


**Figure S3. a-d)** Performance metrics (MAE, MSE, RMSE, R²) of the RF, SVM, and Transformer models in predicting the pI using a fixed test set, evaluated on models trained with varying training set sizes. **e-h)** Performance metrics (MAE, MSE, RMSE, R²) of the RF, SVM, and Transformer models in predicting pI using a non-fixed test set, evaluated on models trained with different training set sizes.

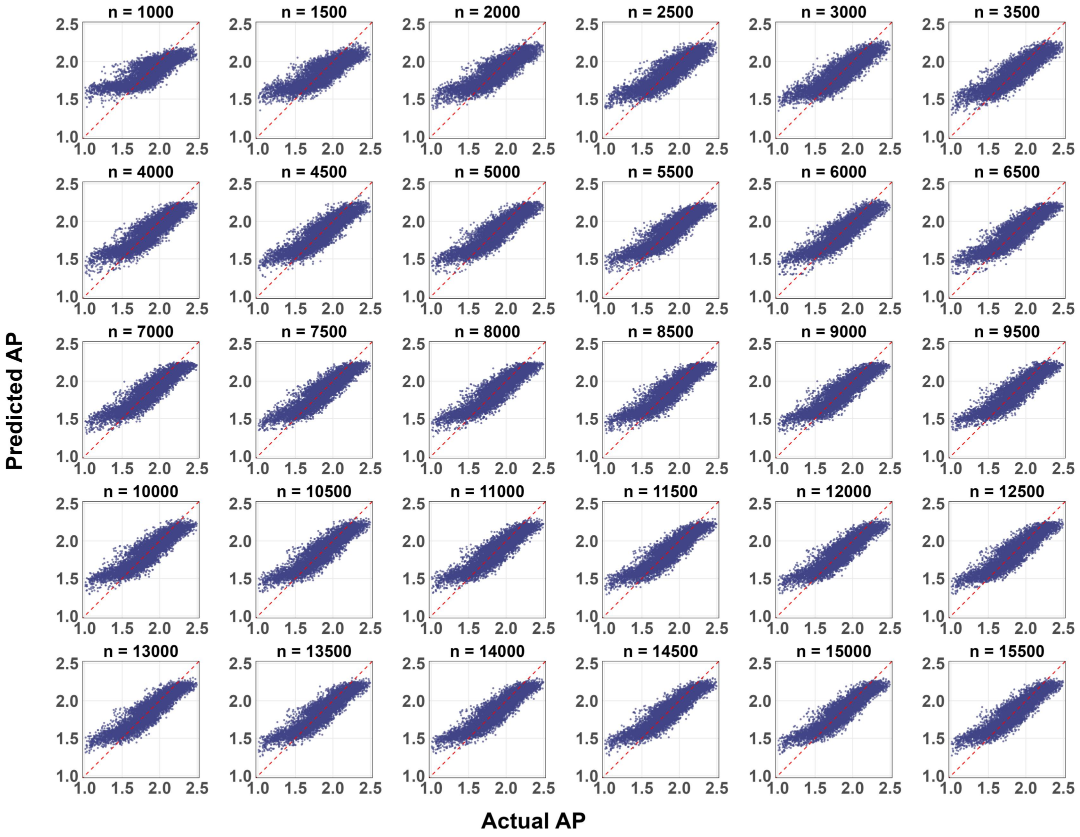


**Figure S4.** Comparison of actual versus predicted AP values from Random Forest models evaluated using an independent test set of 10,000 samples. The variable n represents the size of the training set (sampled dataset).

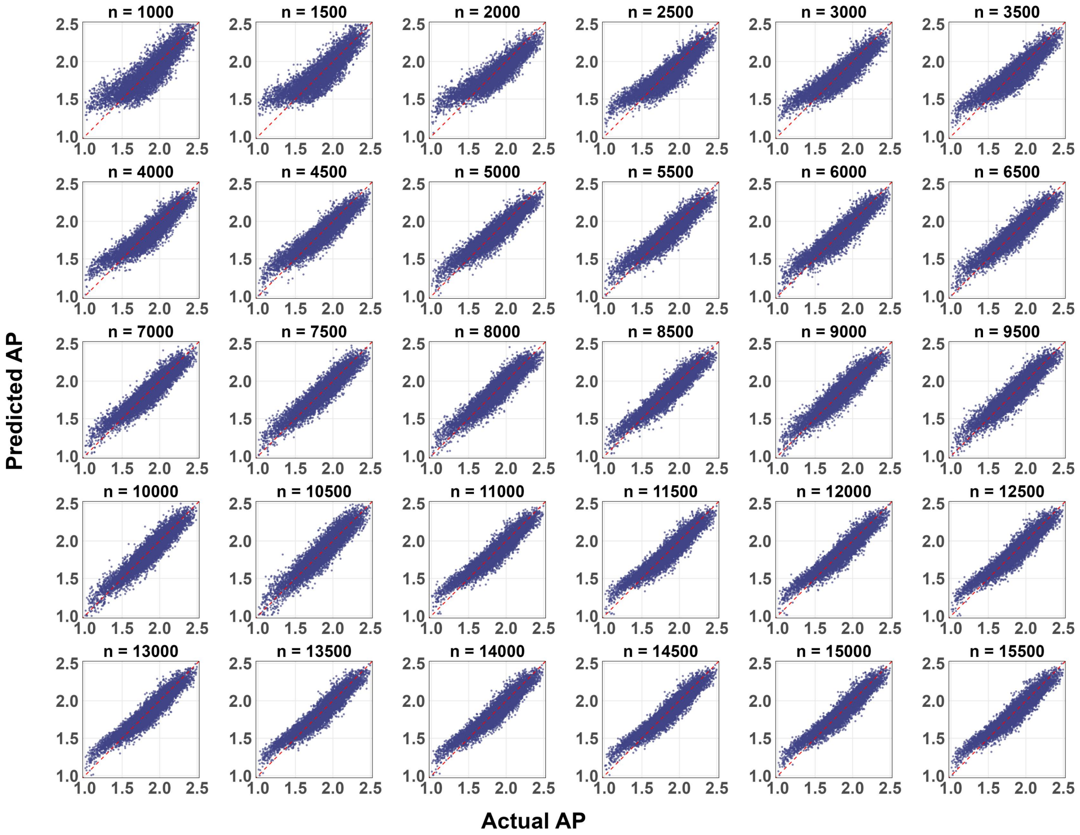


**Figure S5.** Comparison of actual versus predicted AP values from Support Vector Machine models evaluated using an independent test set of 10,000 samples. The variable n represents the size of the training set (sampled dataset).

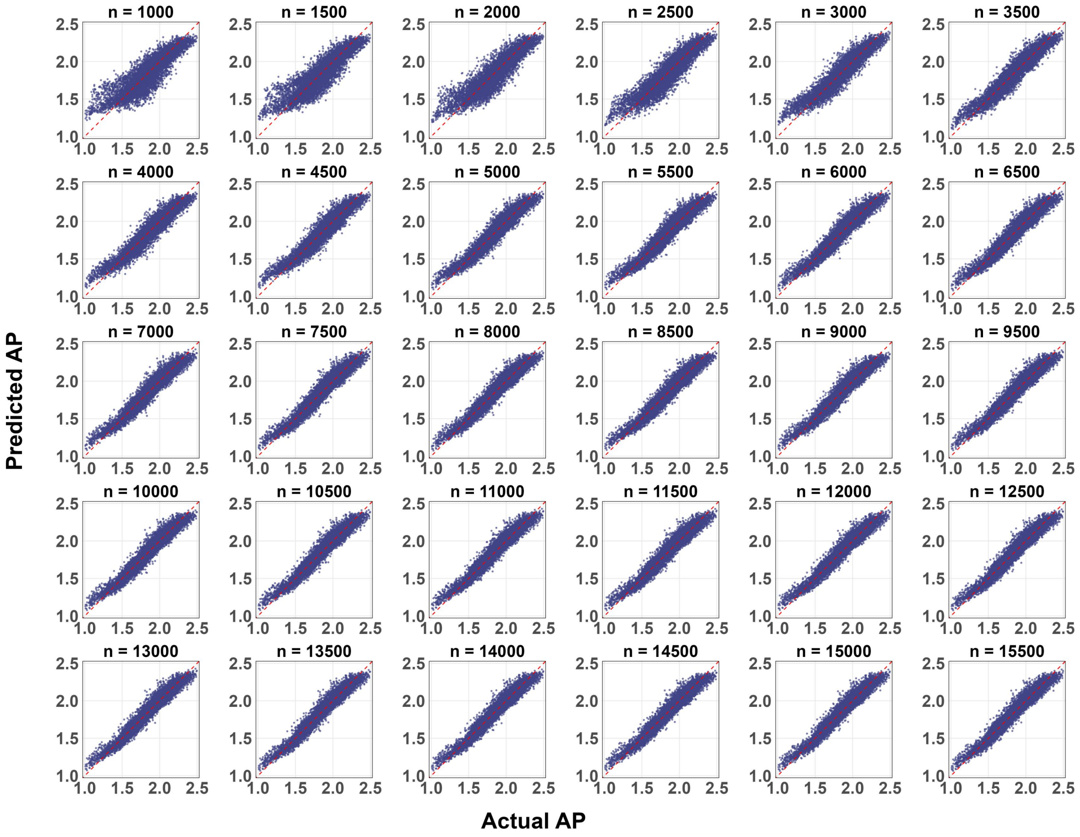


**Figure S6.** Comparison of actual versus predicted AP values from Transformer models evaluated using an independent test set of 10,000 samples. The variable n represents the size of the training set (sampled dataset).

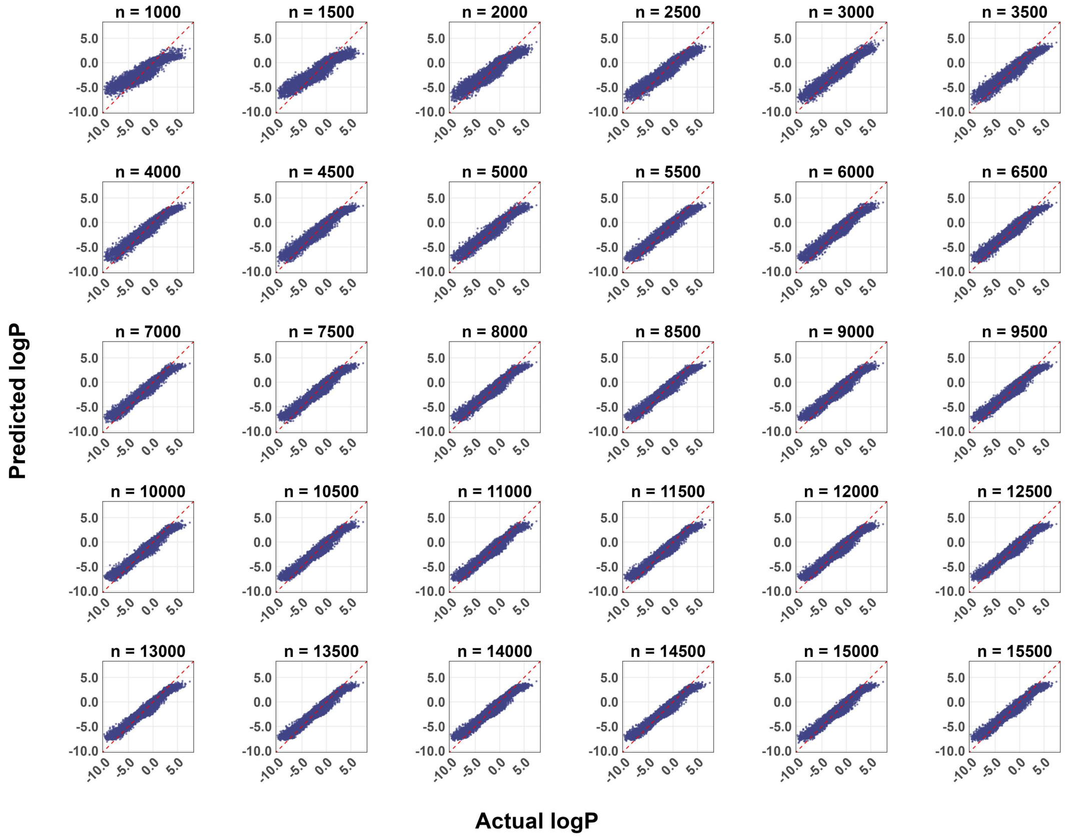


**Figure S7.** Comparison of actual versus predicted logP values from RF models evaluated using an independent test set of 10,000 samples. The variable n represents the size of the training set (sampled dataset).

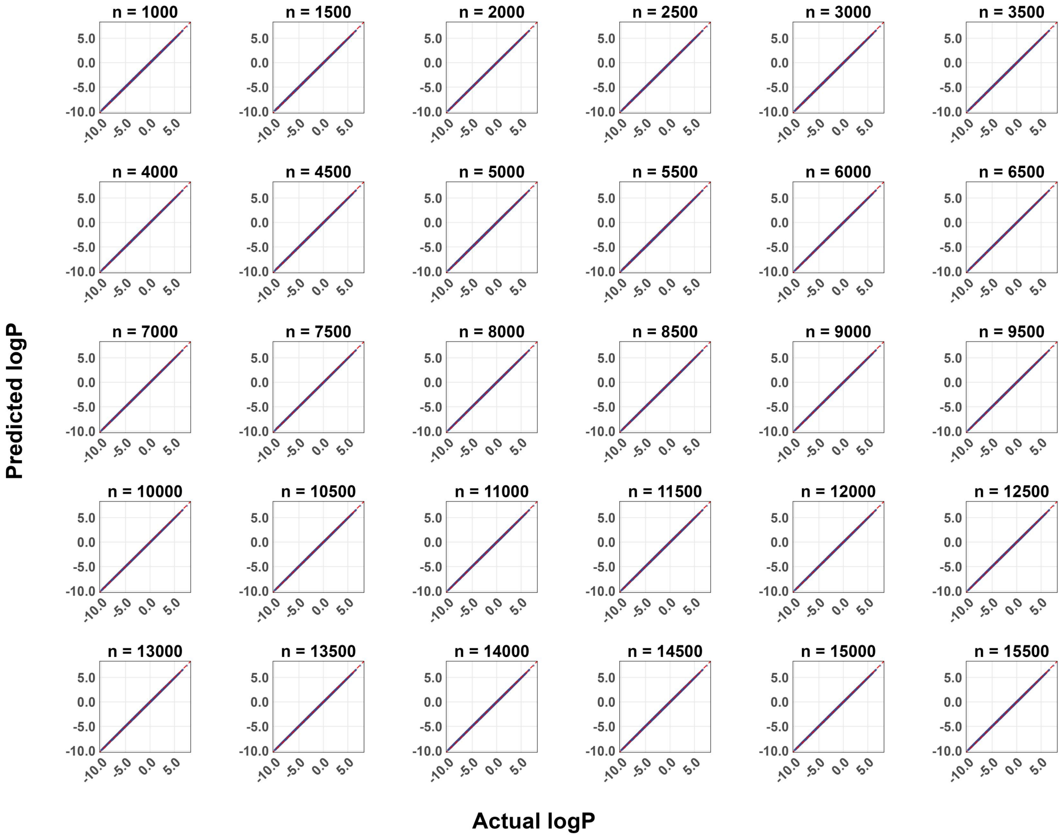


**Figure S8.** Comparison of actual versus predicted logP values from SVM models evaluated using an independent test set of 10,000 samples. The variable n represents the size of the training set (sampled dataset).

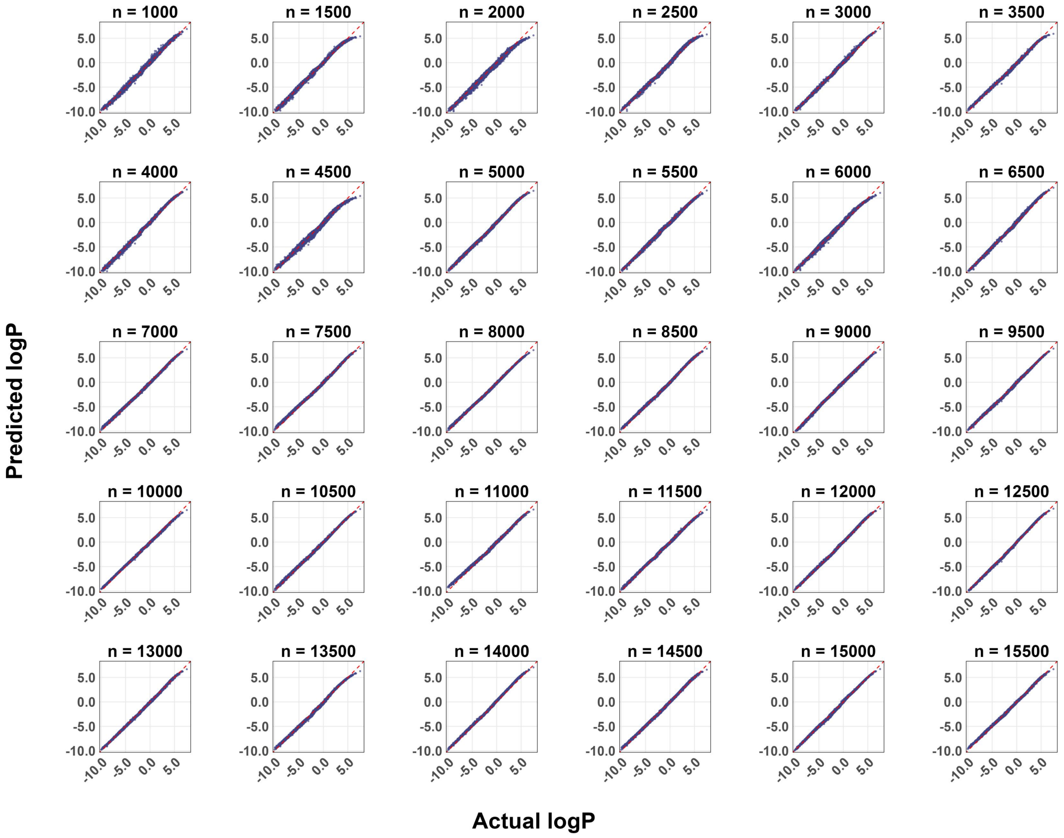


**Figure S9.** Comparison of actual versus predicted logP values from Transformer models evaluated using an independent test set of 10,000 samples. The variable n represents the size of the training set (sampled dataset).

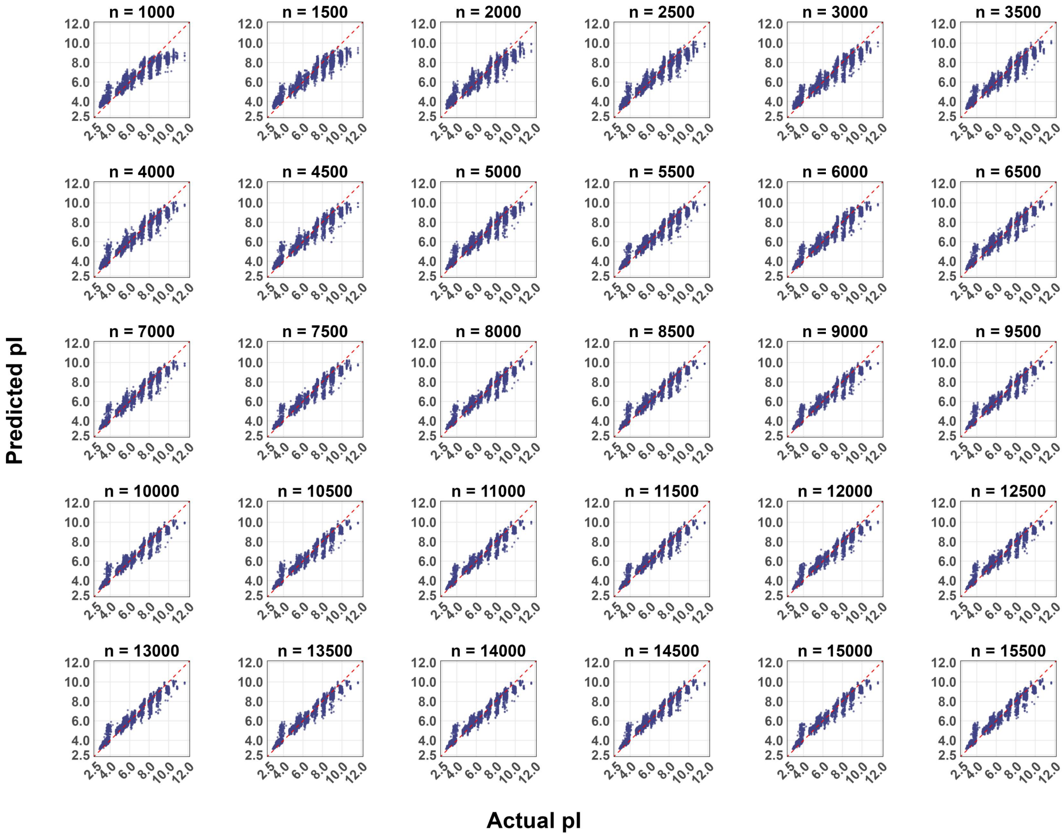


**Figure S10.** Comparison of actual versus predicted pI values from RF models evaluated using an independent test set of 10,000 samples. The variable n represents the size of the training set (sampled dataset).

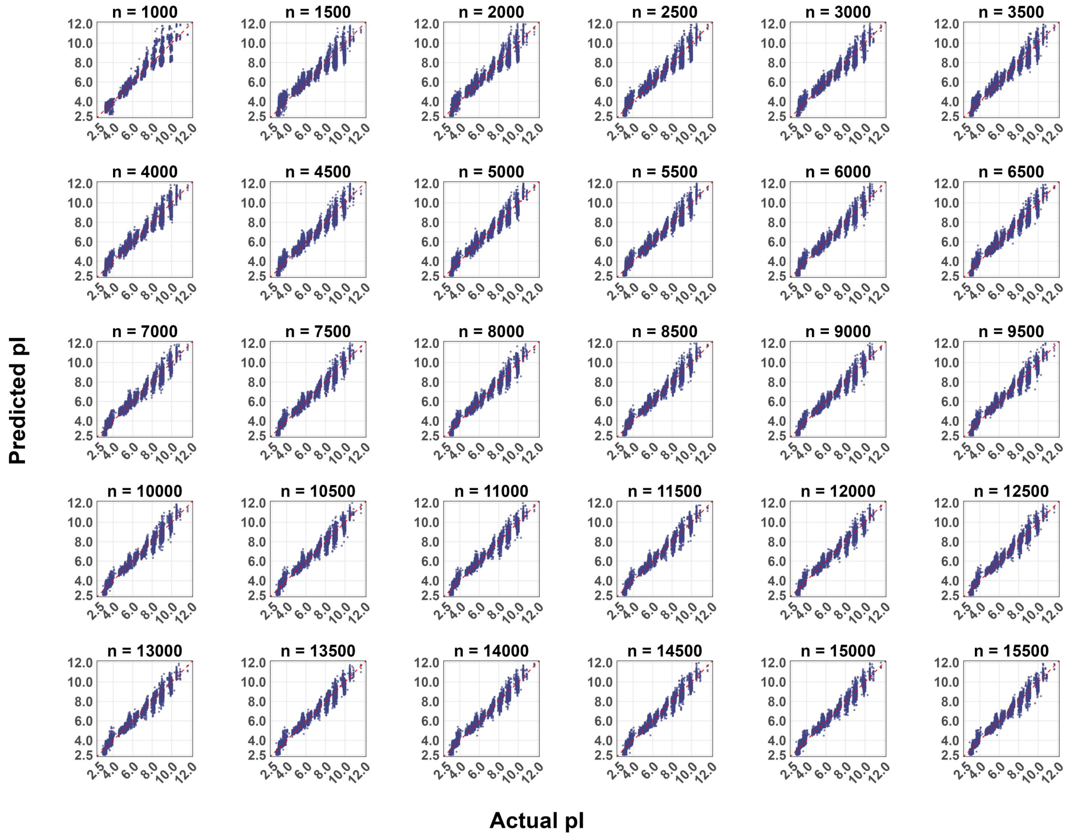


**Figure S11.** Comparison of actual versus predicted pI values from SVM models evaluated using an independent test set of 10,000 samples. The variable n represents the size of the training set (sampled dataset).

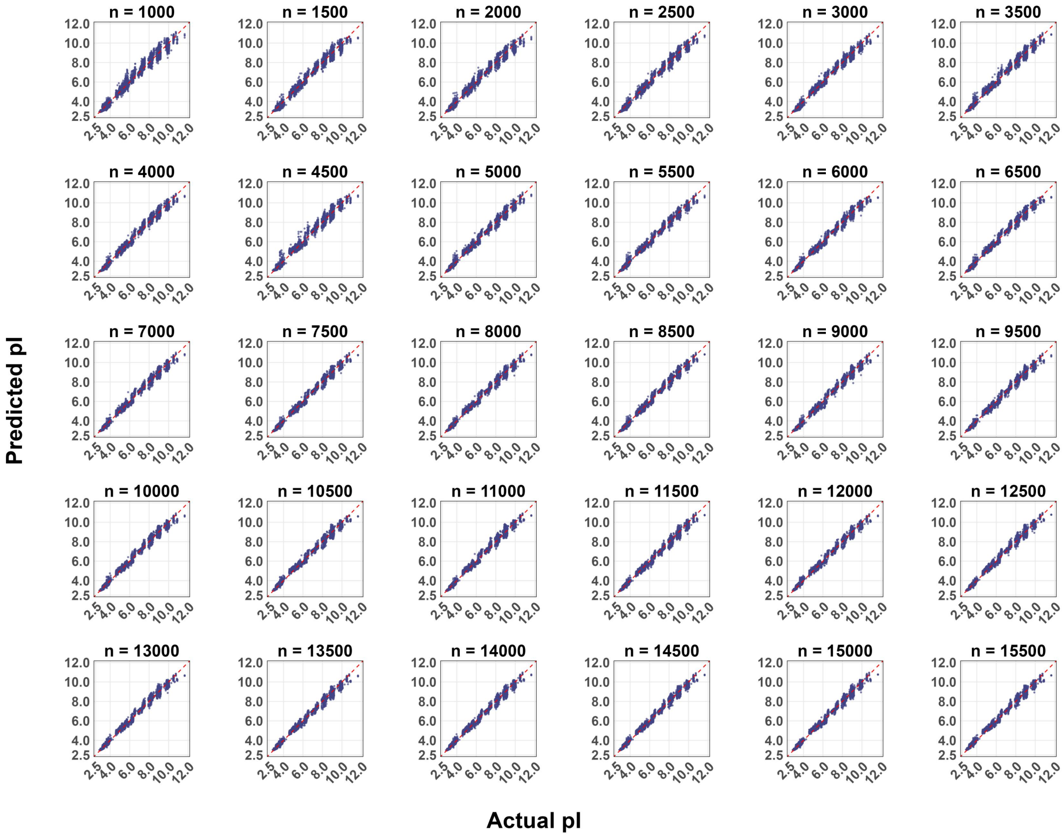


**Figure S12.** Comparison of actual versus predicted pI values from Transformer models evaluated using an independent test set of 10,000 samples. The variable n represents the size of the training set (sampled dataset).

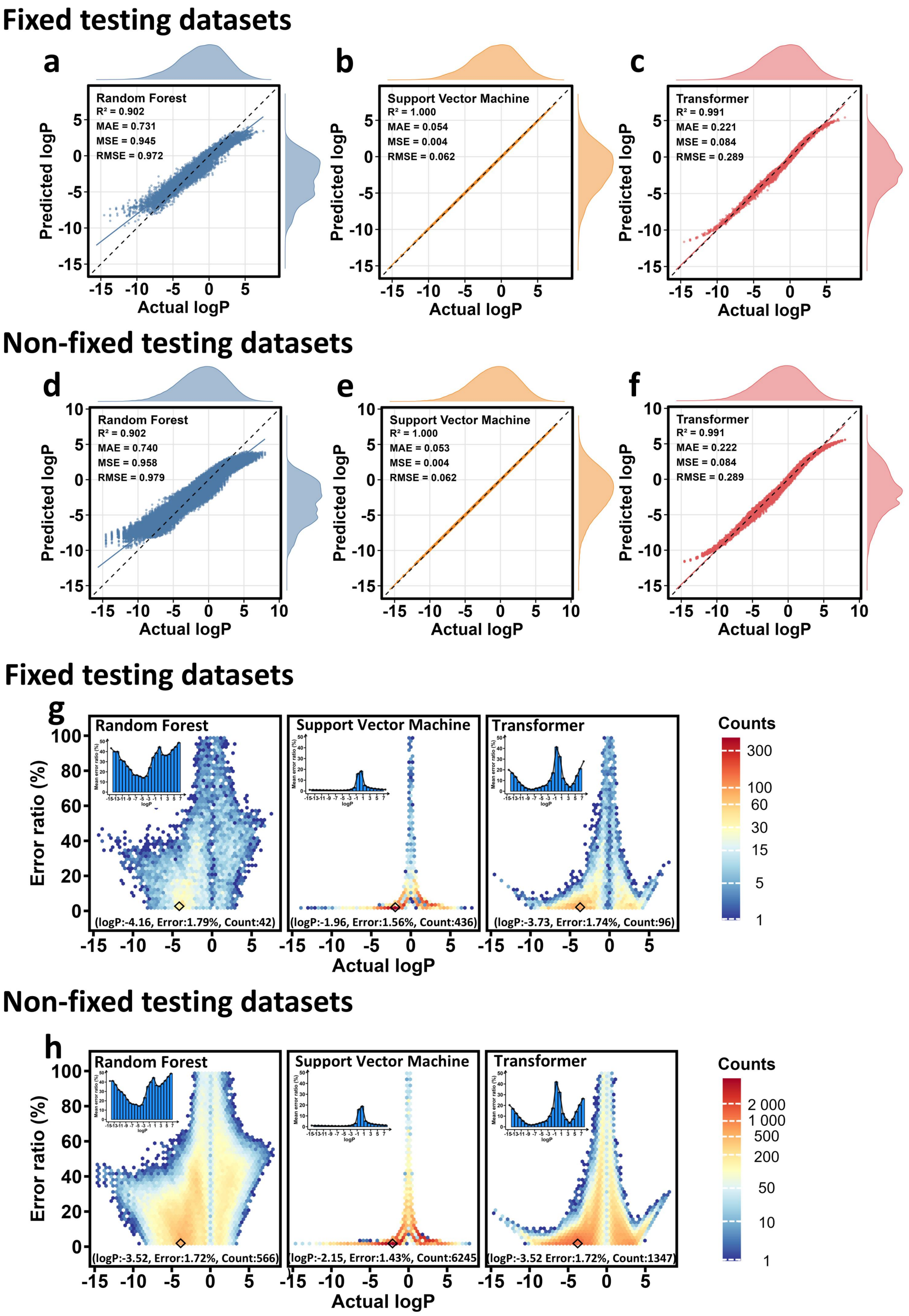


**Figure S13.** Cross-model prediction error analysis of logP. **a-f)** The relationship between the actual and predicted logP values of the RF, SVM and Transformer models is evaluated using a fixed versus a non-fixed testing dataset with a training set of 4,500 (i.e. the threshold of the Transformer model). **g-h)** The relationship between actual logP values and percentage prediction error is visualized by a hexagonal fractal plot when evaluated using a fixed versus a non-fixed testing dataset with the same amount of data in the training set, thus capturing the density of points in the dataset of the three combinations of prediction models.

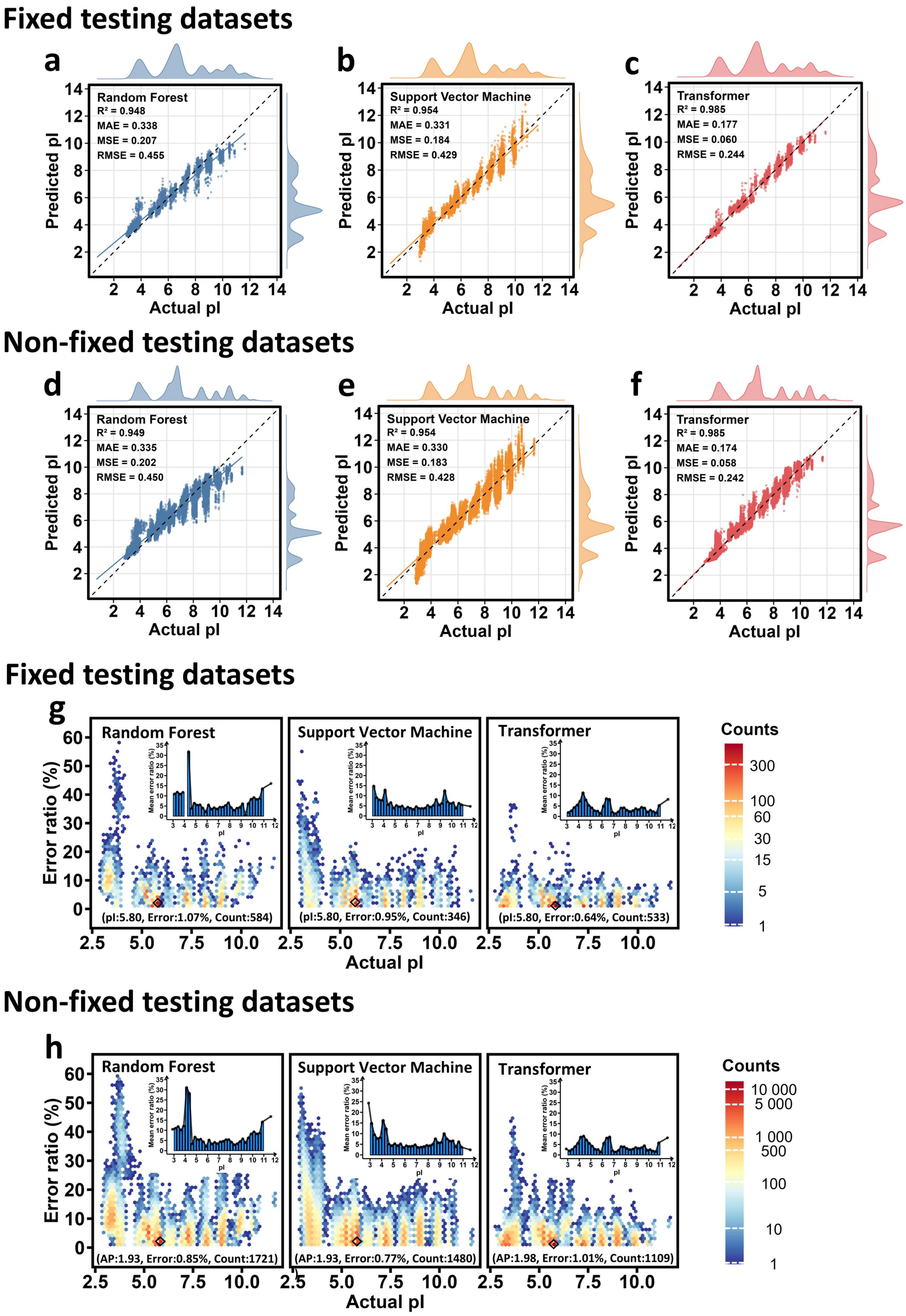


**Figure S14.** Cross-model prediction error analysis of pI. **a-f)** The relationship between the actual and predicted pI values of the RF, SVM and Transformer models is evaluated using a fixed versus a non-fixed testing dataset with a training set of 4,500 (i.e. the threshold of the Transformer model). **g-h)** The relationship between actual pI values and the percentage prediction error is visualized by a hexagonal fractal plot when evaluated using a fixed versus a non-fixed testing dataset with the same amount of data in the training set, thus capturing the density of points in the dataset of the three combinations of prediction models.

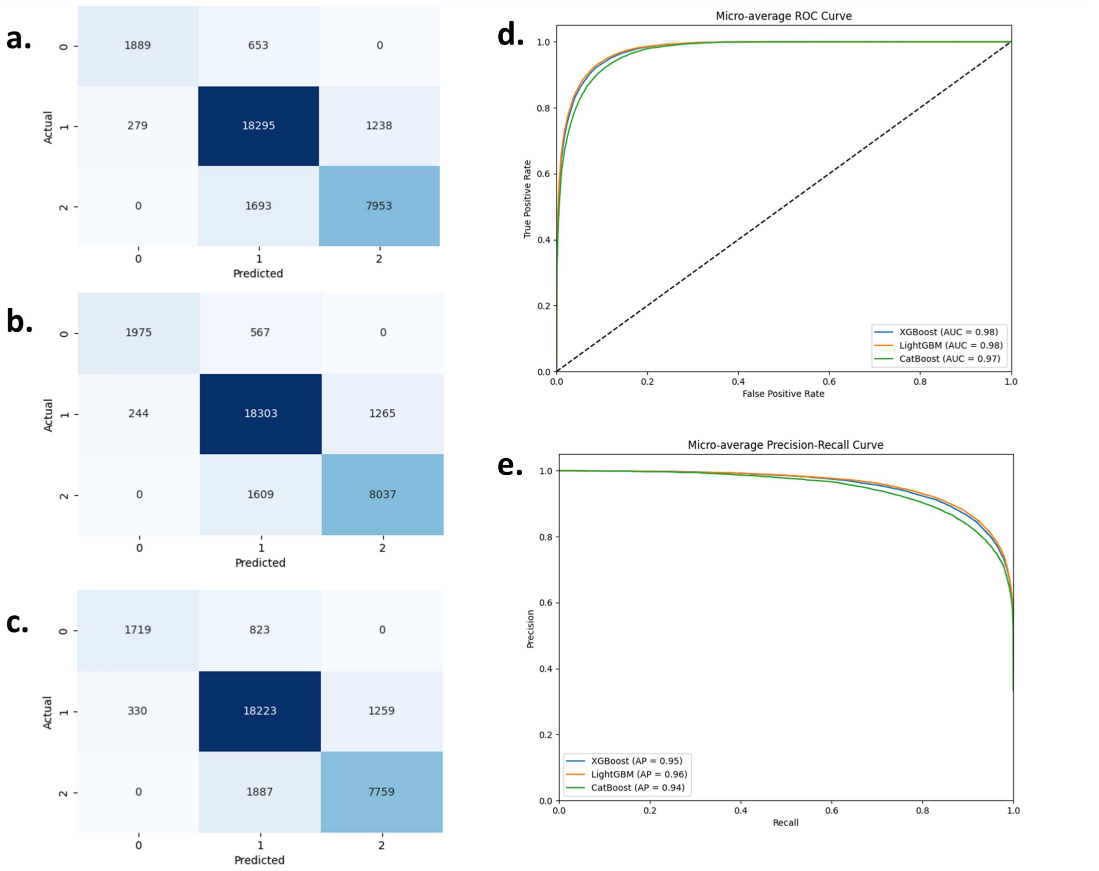


**Figure S15.** Performance Evaluation of Different tree Models for Multiclass classification. **a)** Confusion Matrix for LightGBM. It shows the performance of LightGBM in classifying three classes (0, 1, 2). The matrix indicates the number of true positive, false negative, and false positive predictions for each class. **b)** Confusion Matrix for XGBoost displaying the classification performance across the three classes. **c)** Confusion Matrix for CatBoost, presenting its classification performance for the three classes. **d)** Micro-average ROC Curve for all models, comparing their diagnostic ability. The Area Under the Curve (AUC) values of LightGBM, XGBoost, and CatBoost are 0.98, 0.98, and 0.97 respectively. **e)** Micro-average Precision-Recall Curve for all models. The values are given by LightGBM (0.96), XGBoost (0.95), and CatBoost (0.94).

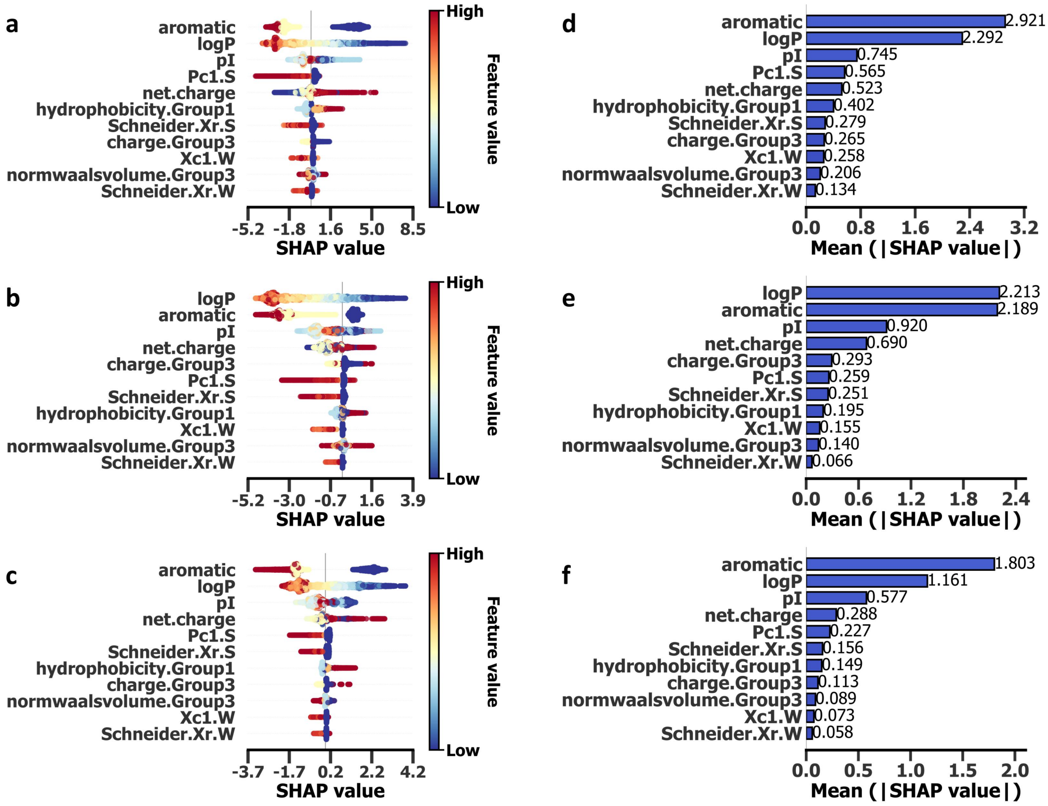


**Figure S16.** Comprehensive SHAP analysis for tetrapeptides' AP values in **Class 0**. **a-c)** Beeswarm plots illustrating the global distribution of features for Class 0 classification across the three models (LightGBM, XGBoost, CatBoost, from top to bottom). Each point represents a sample's SHAP value, with colors indicating feature values (red for high, blue for low). The horizontal position denotes the SHAP value magnitude (negative values indicate negative impact and positive values indicate positive impact), while vertical stacking indicates a higher density of SHAP values in that region. The features are ranked in descending order of importance from top to bottom. **d-f)** The summary bar plot displays the mean absolute SHAP values for 11 features, ranked in descending order of importance. These values represent the average impact of features on predictions, with results corresponding to LightGBM, XGBoost, and CatBoost presented sequentially from top to bottom.

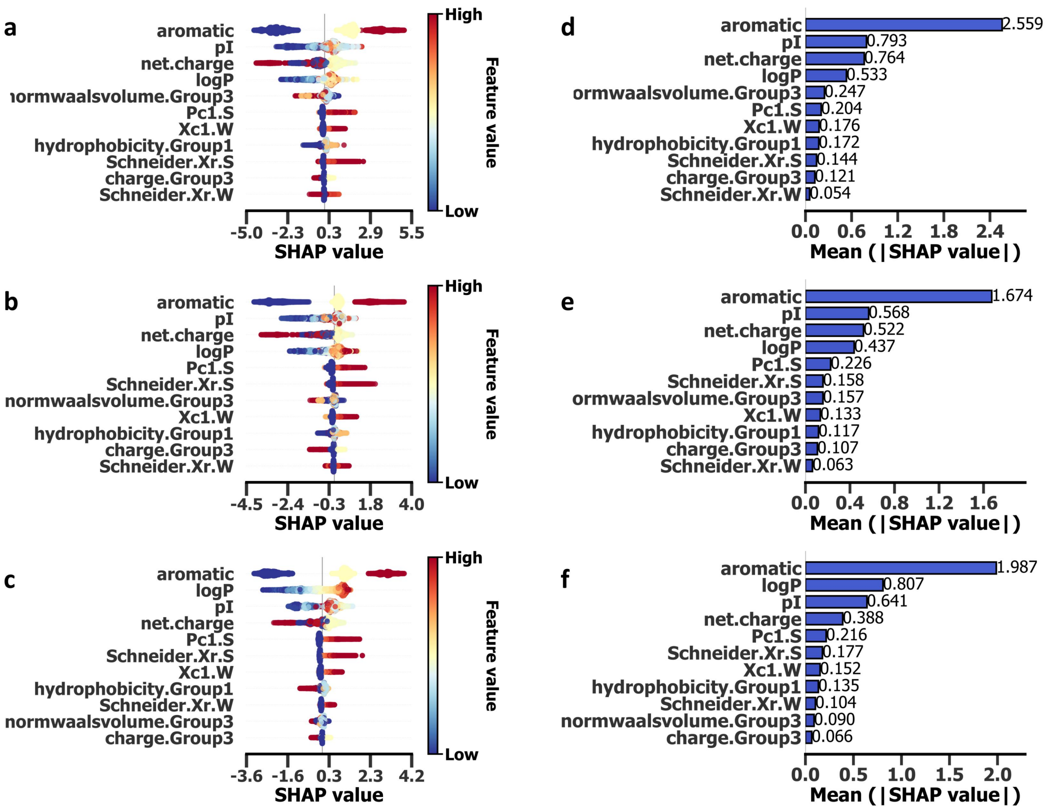


**Figure S17.** Comprehensive SHAP analysis for tetrapeptides' AP values in **Class 2**. **a-c)** Beeswarm plots illustrating the global distribution of features for Class 2 classification across the three models (LightGBM, XGBoost, CatBoost, from top to bottom). Each point represents a sample's SHAP value, with colors indicating feature values (red for high, blue for low). The horizontal position denotes the SHAP value magnitude (negative values indicate negative impact and positive values indicate positive impact), while vertical stacking indicates a higher density of SHAP values in that region. The features are ranked in descending order of importance from top to bottom. d-f) The summary bar plot displays the mean absolute SHAP values for 11 features, ranked in descending order of importance. These values represent the average impact of features on predictions, with results corresponding to LightGBM, XGBoost, and CatBoost presented sequentially from top to bottom.

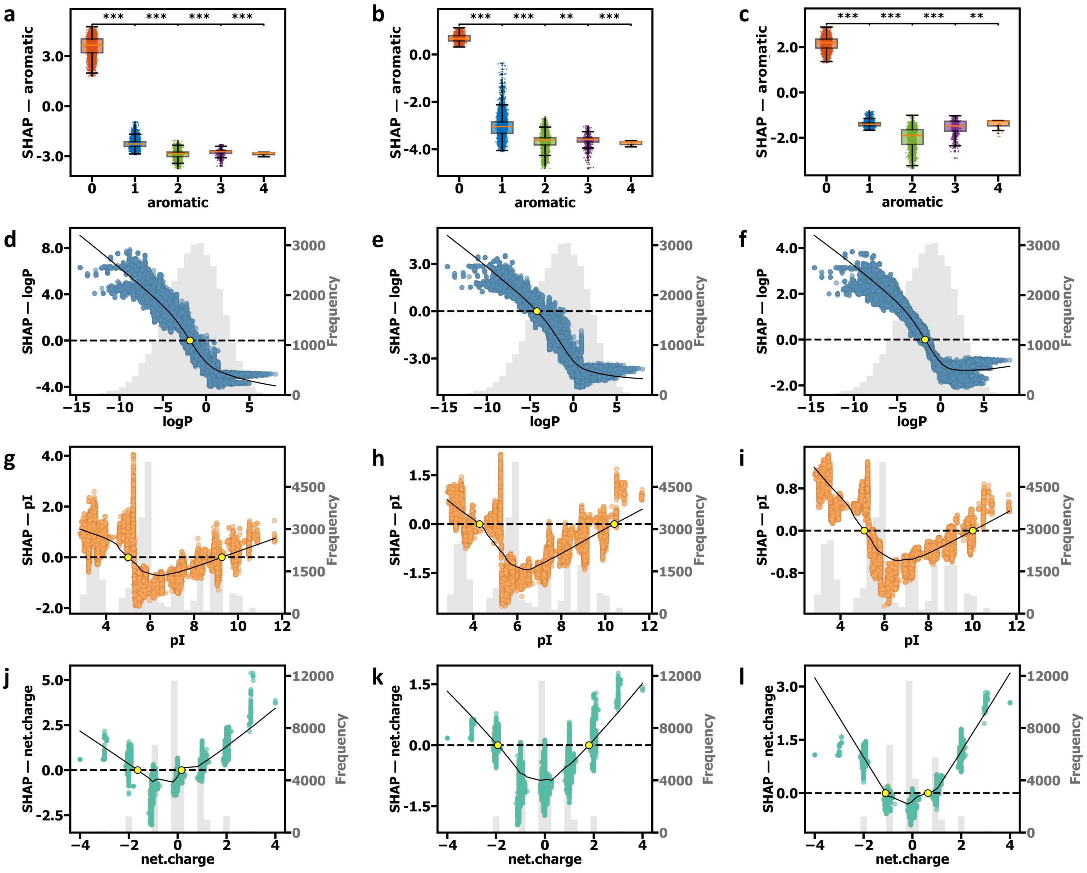


**Figure S18.** SHAP dependence plots for top features in **Class 0** tetrapeptide sequences. **a-c)** Impact of aromatic residue count on Class 0 SHAP values for LightGBM, XGBoost, and CatBoost models. Box plots show SHAP value distributions for different aromatic residue counts (0, 1, 2, 3, and 4). Boxes represent interquartile ranges, middle lines indicate medians, and whiskers extend to 1.5 times the interquartile range. Asterisks or NS (none-significance) above box plots indicate statistical significance levels of SHAP value differences between adjacent groups. Based on Mann-Whitney U tests with Bonferroni correction, significance levels are: *** $p < 0.00025$ (0.001/4), ** $p < 0.0025$ (0.01/4), * $p < 0.0125$ (0.05/4), NS: $p \geq 0.0125$ (not significant). **d-f)** SHAP dependence plots for the logP feature in Class 0 for LightGBM, XGBoost, and CatBoost models. Gray histograms (referring to the right axes) reflect the frequency distribution of logP values in the testing dataset. Black LOWESS (Locally Weighted Scatterplot Smoothing) fitting curves show the overall trend between logP and SHAP values. Black dashed lines indicate the SHAP value of 0 baselines, while yellow markers highlight critical values where the impact of logP on predictions shifts from positive to negative (or vice versa). **g-i)** SHAP dependence plots for the pI feature in Class 0 for LightGBM, XGBoost, and CatBoost models. These plots include scatter points, gray histograms, and black LOWESS curves, similar to d-f. Critical values are marked by yellow points where the impact of pI shifts. **j-l)** SHAP dependence plots for the net charge feature in Class 0 for LightGBM, XGBoost, and CatBoost models. These plots include scatter points, gray histograms, black LOWESS curves, and intersection annotations to comprehensively show how net charge affects model predictions.

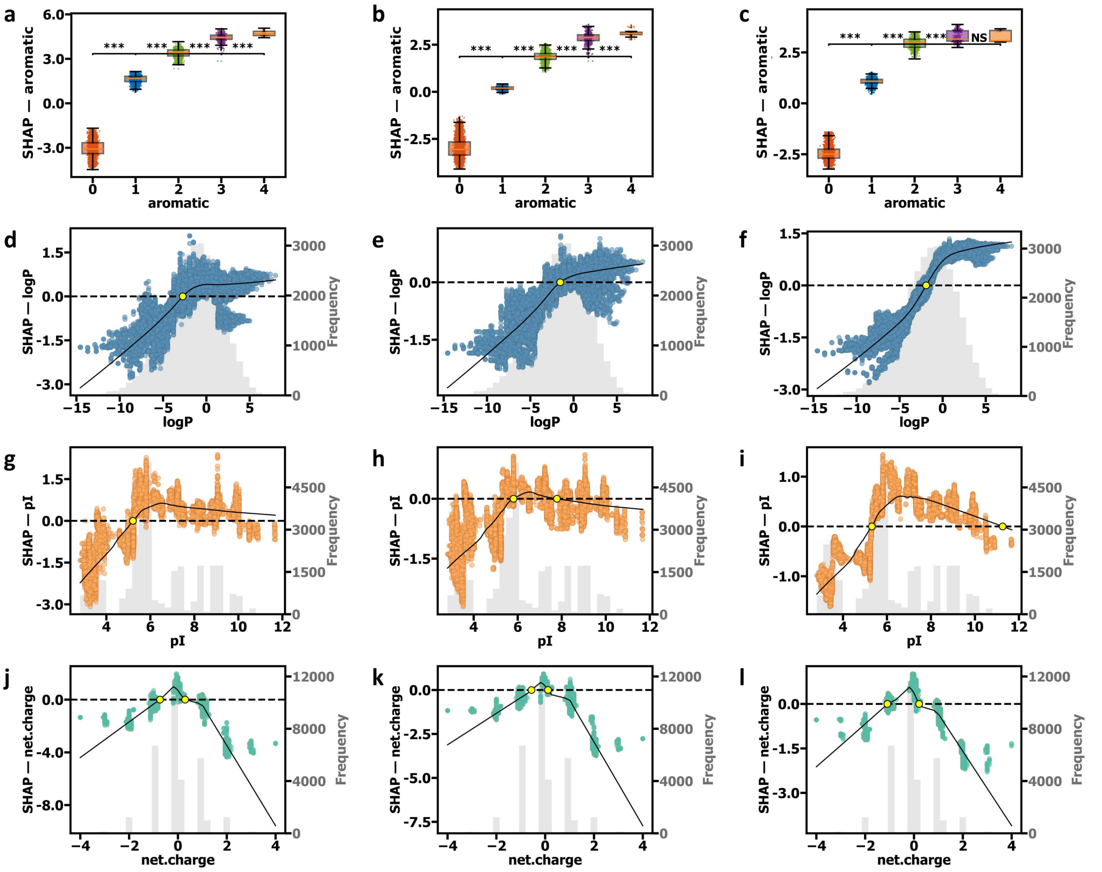

**Figure S19.** SHAP dependence plots for top features in **Class 2** tetrapeptide sequences. **a-c)** Impact of aromatic residue count on Class 2 SHAP values for LightGBM, XGBoost, and CatBoost models. Box plots show SHAP value distributions for different aromatic residue counts (0, 1, 2, 3, and 4). Boxes represent interquartile ranges, middle lines indicate medians, and whiskers extend to 1.5 times the interquartile range. Asterisks or NS (none-significance) above box plots indicate statistical significance levels of SHAP value differences between adjacent groups. Based on Mann-Whitney U tests with Bonferroni correction, significance levels are: *** $p < 0.00025$ (0.001/4), ** $p < 0.0025$ (0.01/4), * $p < 0.0125$ (0.05/4), NS: $p \geq 0.0125$ (not significant). **d-f)** SHAP dependence plots for the logP feature in Class 1 for LightGBM, XGBoost, and CatBoost models. Gray histograms (referring to the right axes) reflect the frequency distribution of logP values in the testing dataset. Black LOWESS (Locally Weighted Scatterplot Smoothing) fitting curves show the overall trend between logP and SHAP values. Black dashed lines indicate the SHAP value of 0 baselines, while yellow markers highlight critical values where the impact of logP on predictions shifts from positive to negative (or vice versa). **g-i)** SHAP dependence plots for the pI feature in Class 2 for LightGBM, XGBoost, and CatBoost models. These plots include scatter points, gray histograms, and black LOWESS curves, similar to d-f. Critical values are marked by yellow points where the impact of pI shifts. **j-l)** SHAP dependence plots for the net charge feature in Class 2 for LightGBM, XGBoost, and CatBoost models. These plots include scatter points, gray histograms, black LOWESS curves, and intersection annotations to comprehensively show how net charge affects model predictions.

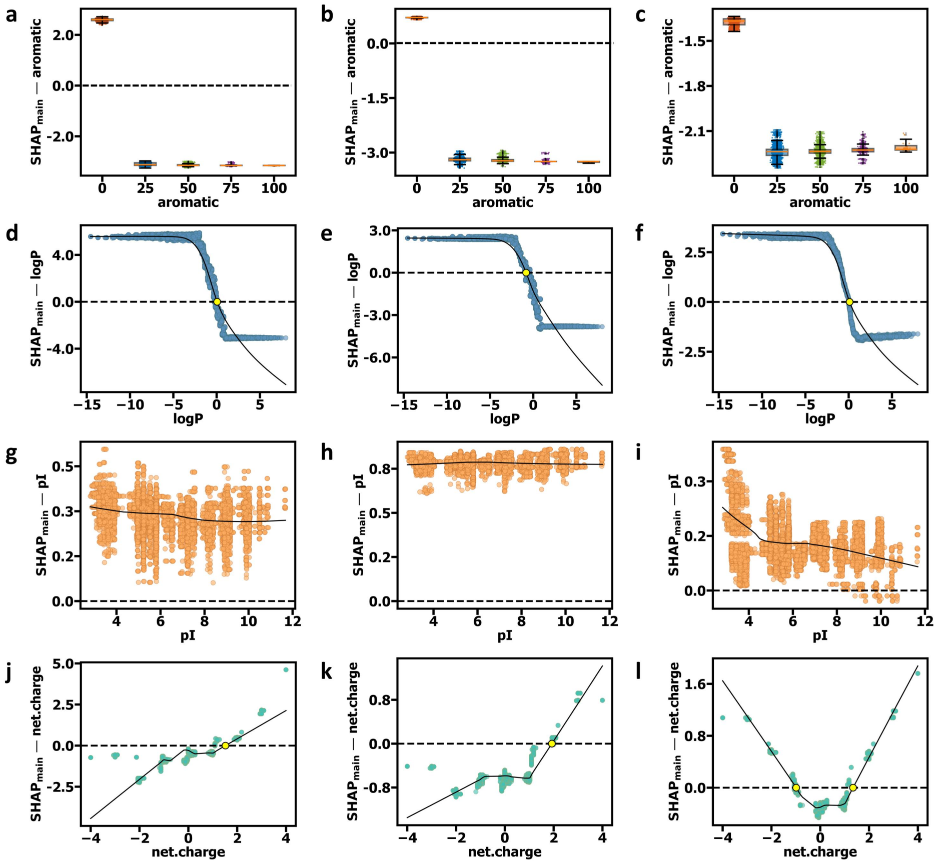


**Figure S20.** Main effect SHAP dependence plots for top features in **Class 0** of AP. **a-c)** SHAP main effect dependence plots for the aromatic property on Class 0 results in the LightGBM, XGBoost, and CatBoost models, from left to right. **d-f)** SHAP main effect dependence plots for the logP feature on Class 0 results in the LightGBM, XGBoost, and CatBoost models, from left to right. **g-i)** SHAP main effect dependence plots for the pI feature on Class 0 results in the LightGBM, XGBoost, and CatBoost models, from left to right. **j-l)** SHAP main effect dependence plots for the net charge feature on Class 0 results. The black LOWESS curve in each Figure Sdepicts the overall trend between different properties and SHAP main effect values.

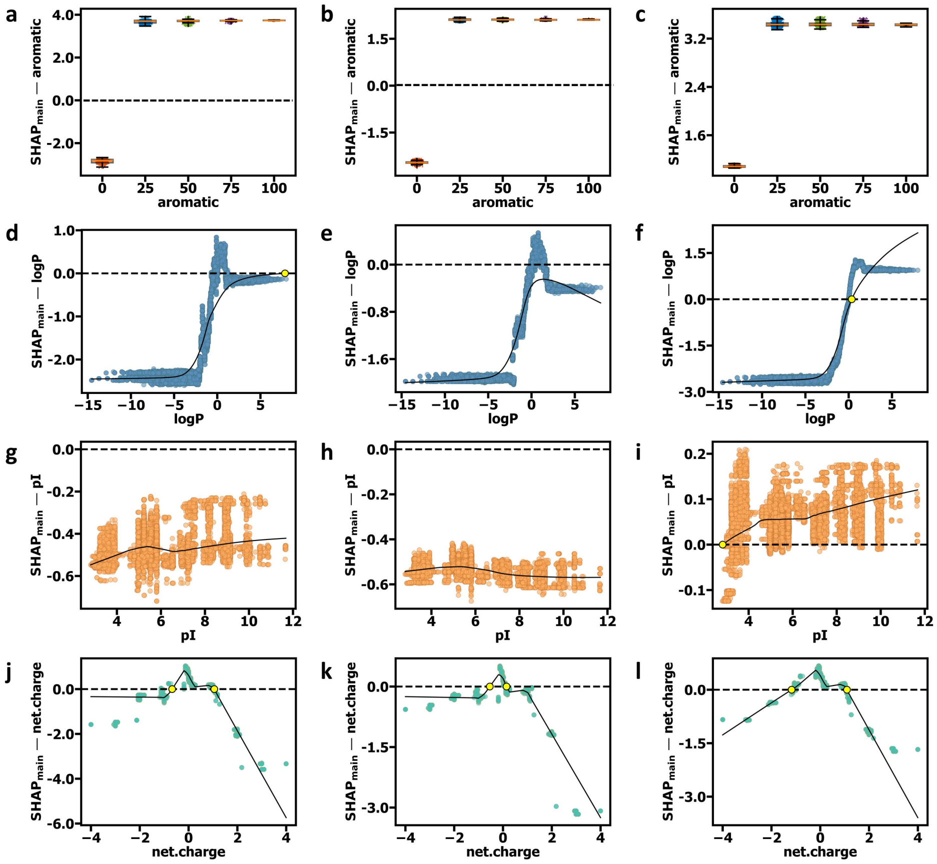

**Figure S21.** Main effect SHAP dependence plots for top features in **Class 2** of AP. **a-c)** SHAP main effect dependence plots for the aromatic property on Class 2 results in the LightGBM, XGBoost, and CatBoost models, from left to right. **d-f)** SHAP main effect dependence plots for the logP feature on Class 2 results in the LightGBM, XGBoost, and CatBoost models, from left to right. **g-i)** SHAP main effect dependence plots for the pI feature on Class 2 results in the LightGBM, XGBoost, and CatBoost models, from left to right. **j-l)** SHAP main effect dependence plots for the net charge feature on Class 2 results. The black LOWESS curve in each Figure Sdepicts the overall trend between different properties and SHAP main effect values.

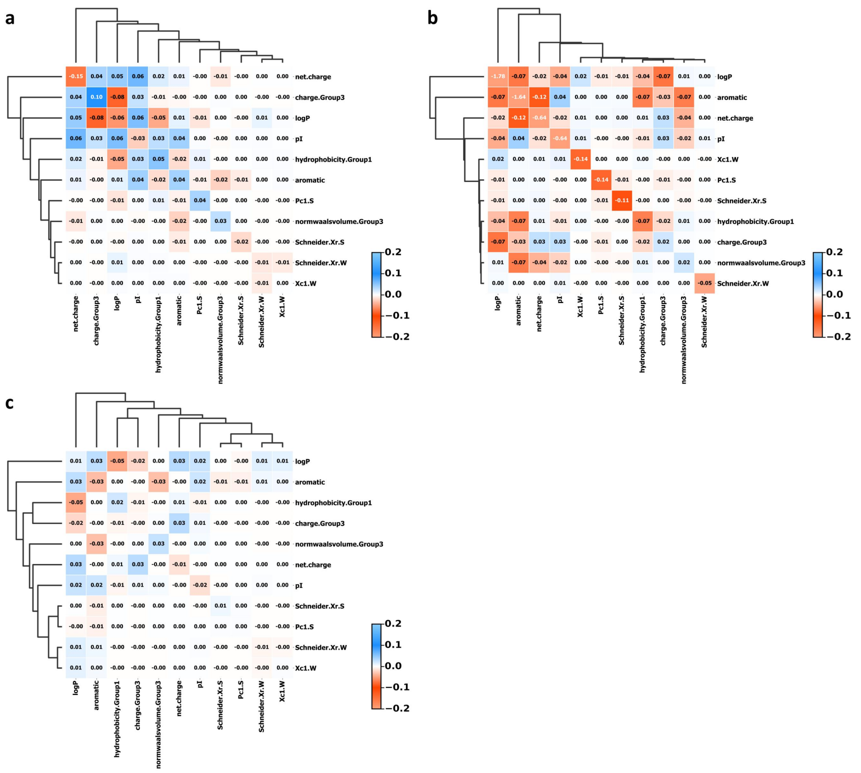


**Figure S22.** Heatmaps of SHAP interaction values for 11 features in **Class 0**. **a-c)** The non-diagonal elements quantitatively represent the strength and direction (positive in blue, negative in orange) of interaction effects between features, with diagonal elements replaced by the main effect values of individual features. The figures are arranged in the following order: LightGBM, XGBoost, and CatBoost.

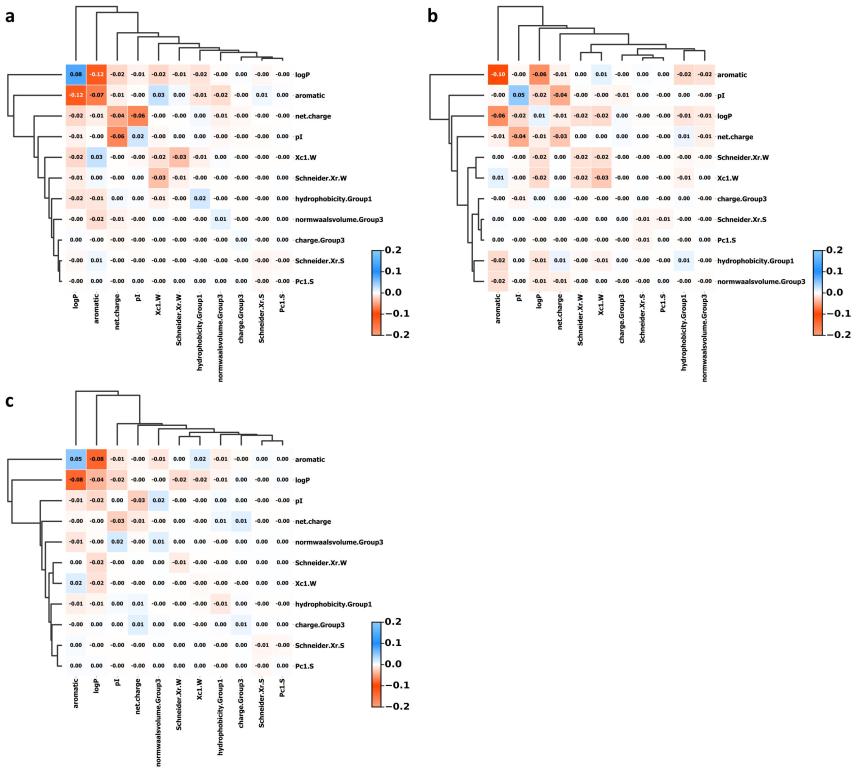

**Figure S23.** Heatmaps of SHAP interaction values for 11 features in **Class 1**. **a-c)** The non-diagonal elements quantitatively represent the strength and direction (positive in blue, negative in orange) of interaction effects between features, with diagonal elements replaced by the main effect values of individual features. The figures are arranged in the following order: LightGBM, XGBoost, and CatBoost.

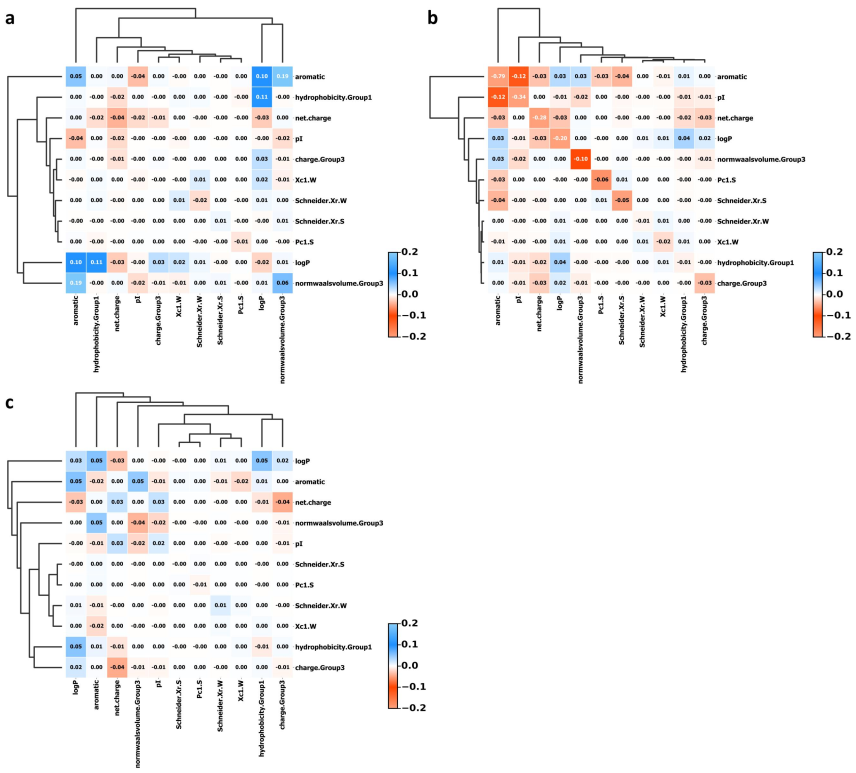

**Figure S24.** Heatmaps of SHAP interaction values for 11 features in **Class 2**. **a-c)** The non-diagonal elements quantitatively represent the strength and direction (positive in blue, negative in orange) of interaction effects between features, with diagonal elements replaced by the main effect values of individual features. The figures are arranged in the following order: LightGBM, XGBoost, and CatBoost.

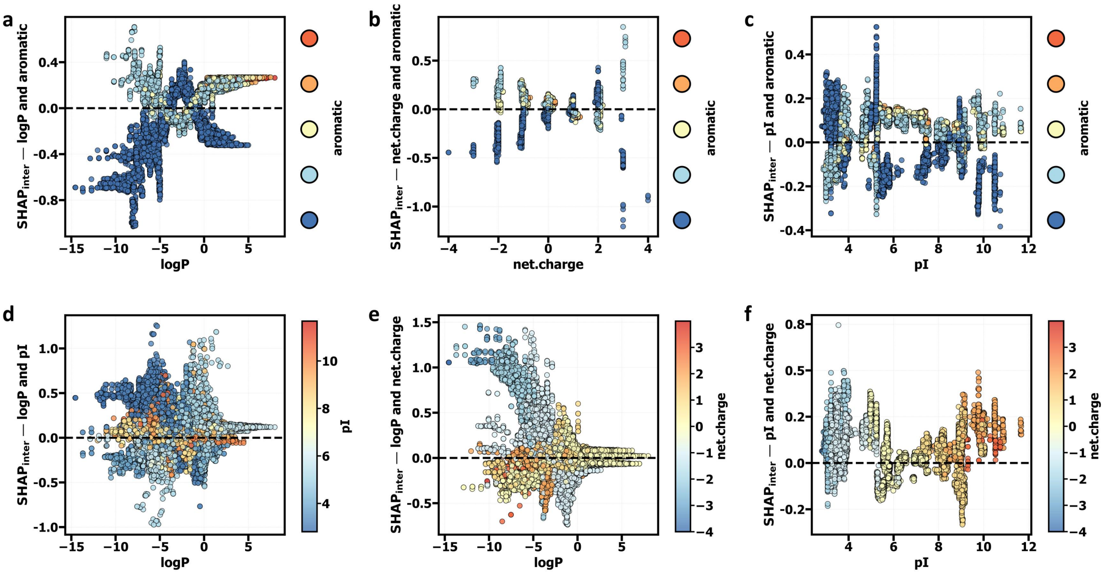


**Figure S25.** SHAP interaction effect dependence plots for the top 4 features (logP, aromatic, pI, net.charge) in **Class 0** for the **LightGBM** model. **a-c)** SHAP interaction effect dependence plots for logP and aromatic (a), net.charge and aromatic (b), and pI and aromatic (c). The color gradient represents the interaction effect magnitude for the aromatic feature. **d-f)** SHAP interaction effect dependence plots for logP and pI (d), logP and net.charge (e), and pI and net.charge (f). The color gradient indicates the interaction effect magnitude for pI (d), net.charge (e), and net.charge (f), respectively. Black dashed lines indicate where SHAP effect values are zero, showing the direction and strength of feature interaction impacts on the model's predictions.

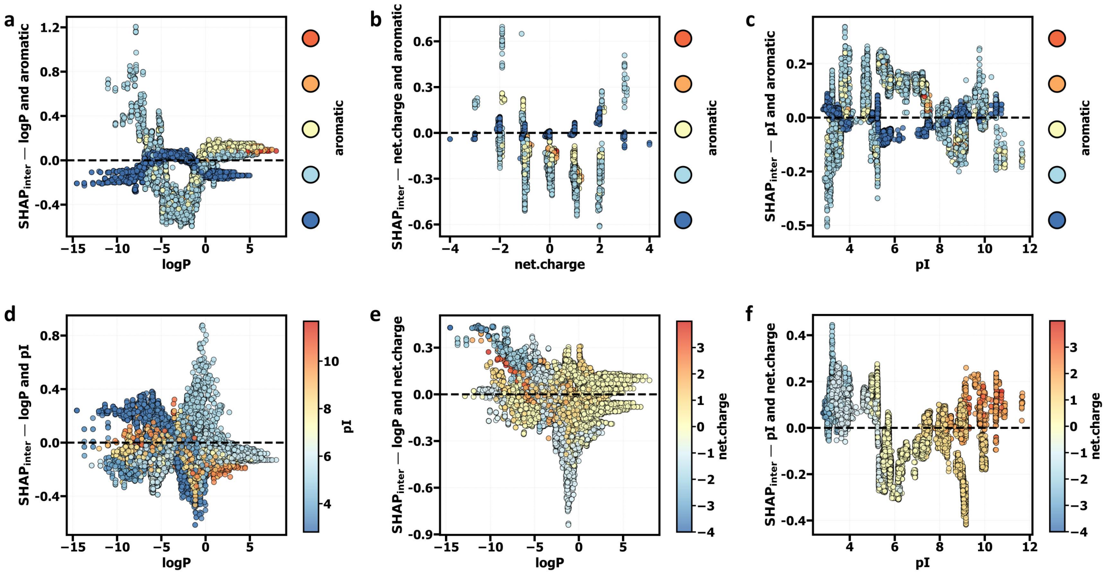


**Figure S26.** SHAP interaction effect dependence plots for the top 4 features (logP, aromatic, pI, net.charge) in **Class 0** for the **XGboost** model. **a-c)** SHAP interaction effect dependence plots for logP and aromatic (a), net.charge and aromatic (b), and pI and aromatic (c). The color gradient represents the interaction effect magnitude for the aromatic feature. **d-f)** SHAP interaction effect dependence plots for logP and pI (d), logP and net.charge (e), and pI and net.charge (f). The color gradient indicates the interaction effect magnitude for pI (d), net.charge (e), and net.charge (f), respectively. Black dashed lines indicate where SHAP effect values are zero, showing the direction and strength of feature interaction impacts on the model's predictions.

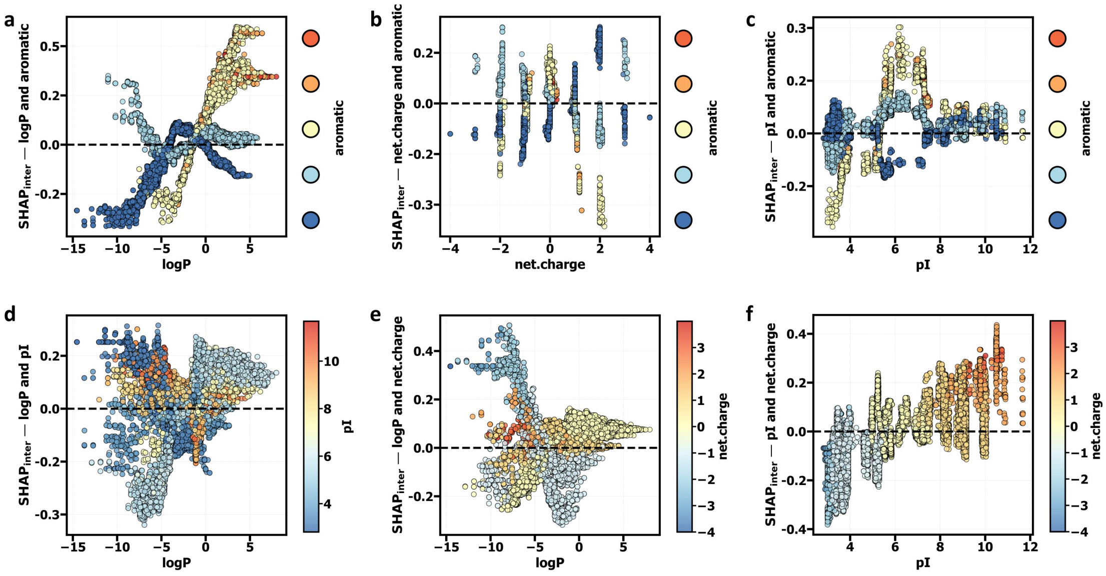


**Figure S27.** SHAP interaction effect dependence plots for the top 4 features (logP, aromatic, pI, net.charge) in **Class 0** for the **CatBoost** model. **a-c)** SHAP interaction effect dependence plots for logP and aromatic (a), net.charge and aromatic (b), and pI and aromatic (c). The color gradient represents the interaction effect magnitude for the aromatic feature. **d-f)** SHAP interaction effect dependence plots for logP and pI (d), logP and net.charge (e), and pI and net.charge (f). The color gradient indicates the interaction effect magnitude for pI (d), net.charge (e), and net.charge (f), respectively. Black dashed lines indicate where SHAP effect values are zero, showing the direction and strength of feature interaction impacts on the model's predictions.

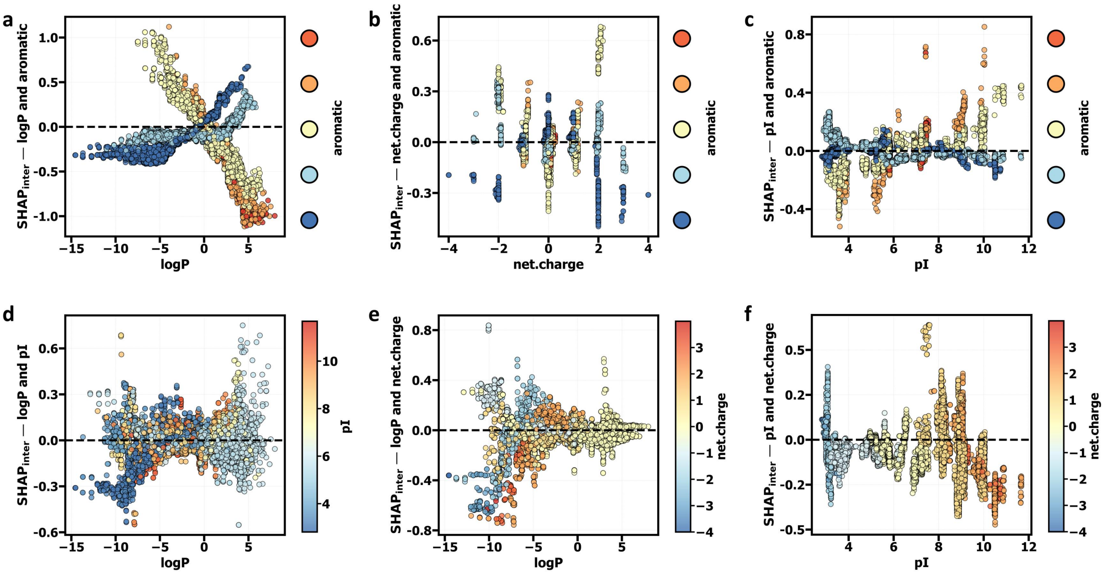


**Figure S28.** SHAP interaction effect dependence plots for the top 4 features (logP, aromatic, pI, net.charge) in **Class 1** for the **LightGBM** model. **a-c)** SHAP interaction effect dependence plots for logP and aromatic (a), net.charge and aromatic (b), and pI and aromatic (c). The color gradient represents the interaction effect magnitude for the aromatic feature. **d-f)** SHAP interaction effect dependence plots for logP and pI (d), logP and net.charge (e), and pI and net.charge (f). The color gradient indicates the interaction effect magnitude for pI (d), net.charge (e), and net.charge (f), respectively. Black dashed lines indicate where SHAP effect values are zero, showing the direction and strength of feature interaction impacts on the model's predictions.

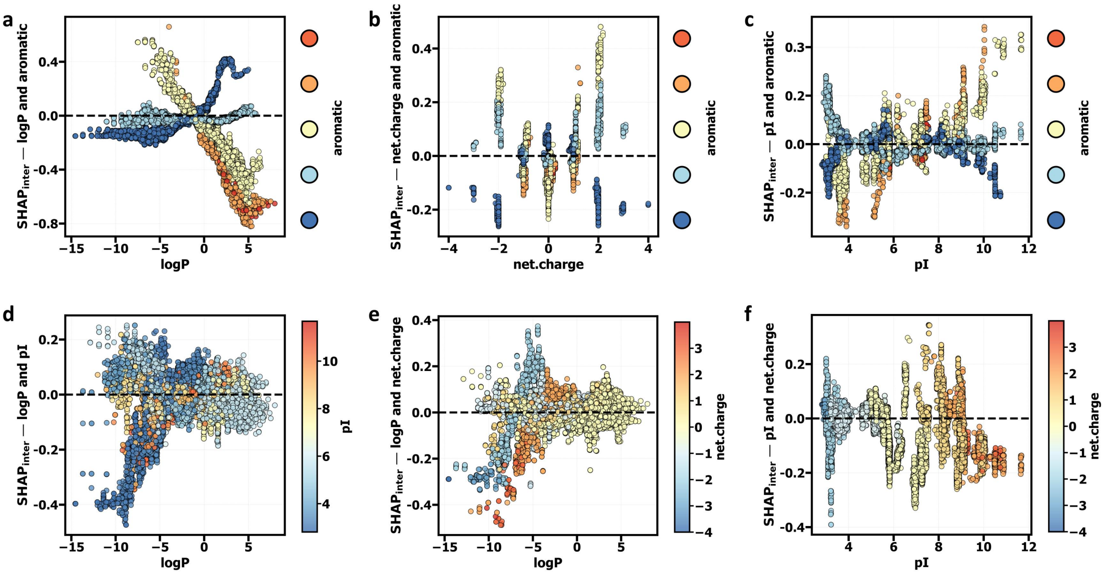


**Figure S29.** SHAP interaction effect dependence plots for the top 4 features (logP, aromatic, pI, net.charge) in **Class 1** for the **XGBoost** model. **a-c)** SHAP interaction effect dependence plots for logP and aromatic (a), net.charge and aromatic (b), and pI and aromatic (c). The color gradient represents the interaction effect magnitude for the aromatic feature. **d-f)** SHAP interaction effect dependence plots for logP and pI (d), logP and net.charge (e), and pI and net.charge (f). The color gradient indicates the interaction effect magnitude for pI (d), net.charge (e), and net.charge (f), respectively. Black dashed lines indicate where SHAP effect values are zero, showing the direction and strength of feature interaction impacts on the model's predictions.

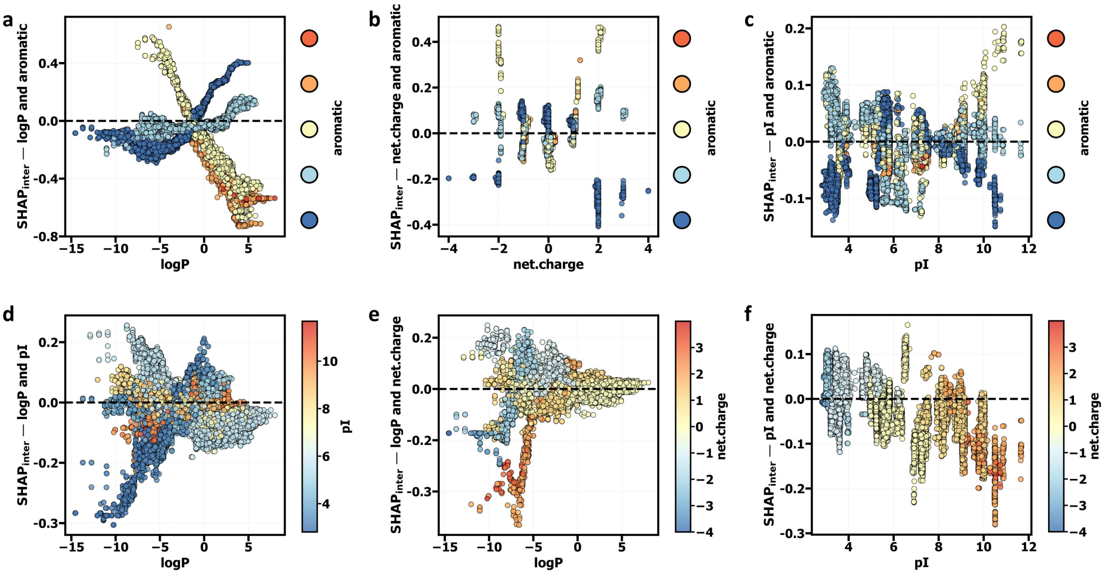


**Figure S30.** SHAP interaction effect dependence plots for the top 4 features (logP, aromatic, pI, net.charge) in **Class 1** for the **CatBoost** model. **a-c)** SHAP interaction effect dependence plots for logP and aromatic (a), net.charge and aromatic (b), and pI and aromatic (c). The color gradient represents the interaction effect magnitude for the aromatic feature. **d-f)** SHAP interaction effect dependence plots for logP and pI (d), logP and net.charge (e), and pI and net.charge (f). The color gradient indicates the interaction effect magnitude for pI (d), net.charge (e), and net.charge (f), respectively. Black dashed lines indicate where SHAP effect values are zero, showing the direction and strength of feature interaction impacts on the model's predictions.

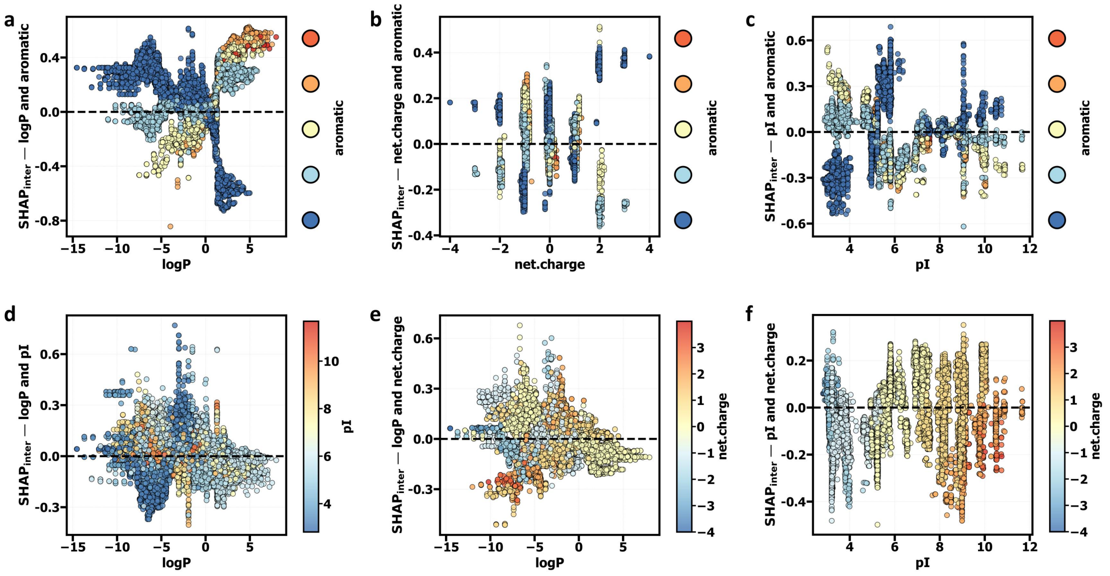


**Figure S31.** SHAP interaction effect dependence plots for the top 4 features (logP, aromatic, pI, net.charge) in **Class 2** for the **LightGBM** model. **a-c)** SHAP interaction effect dependence plots for logP and aromatic (a), net.charge and aromatic (b), and pI and aromatic (c). The color gradient represents the interaction effect magnitude for the aromatic feature. **d-f)** SHAP interaction effect dependence plots for logP and pI (d), logP and net.charge (e), and pI and net.charge (f). The color gradient indicates the interaction effect magnitude for pI (d), net.charge (e), and net.charge (f), respectively. Black dashed lines indicate where SHAP effect values are zero, showing the direction and strength of feature interaction impacts on the model's predictions.

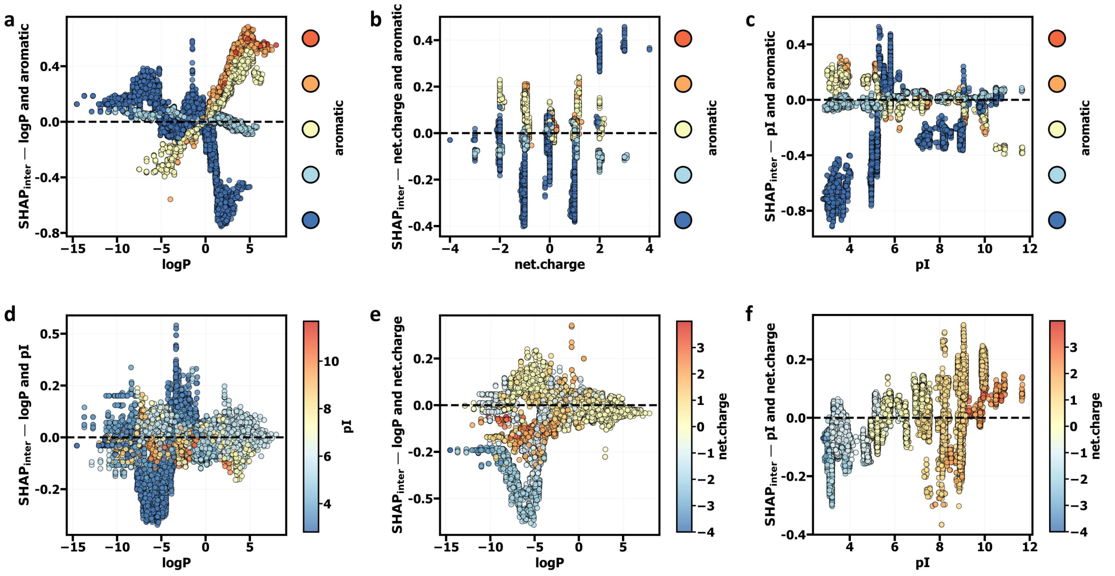


**Figure S32.** SHAP interaction effect dependence plots for the top 4 features (logP, aromatic, pI, net.charge) in **Class 2** for the **XGBoost** model. **a-c)** SHAP interaction effect dependence plots for logP and aromatic (a), net.charge and aromatic (b), and pI and aromatic (c). The color gradient represents the interaction effect magnitude for the aromatic feature. **d-f)** SHAP interaction effect dependence plots for logP and pI (d), logP and net.charge (e), and pI and net.charge (f). The color gradient indicates the interaction effect magnitude for pI (d), net.charge (e), and net.charge (f), respectively. Black dashed lines indicate where SHAP effect values are zero, showing the direction and strength of feature interaction impacts on the model's predictions.

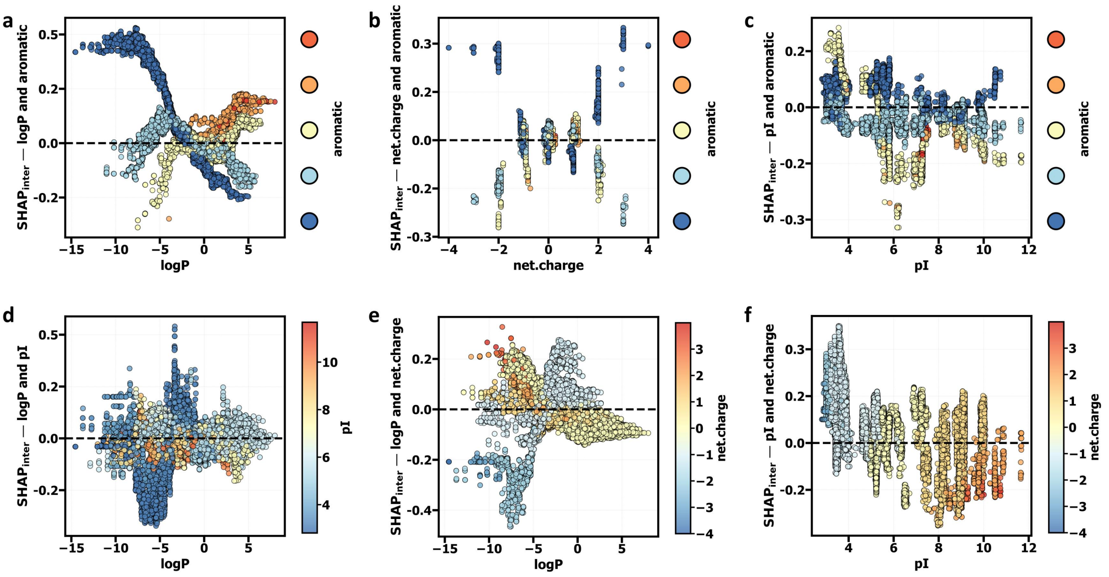


**Figure S33.** SHAP interaction effect dependence plots for the top 4 features (logP, aromatic, pI, net.charge) in **Class 2** for the **CatBoost** model. **a-c)** SHAP interaction effect dependence plots for logP and aromatic (a), net.charge and aromatic (b), and pI and aromatic (c). The color gradient represents the interaction effect magnitude for the aromatic feature. **d-f)** SHAP interaction effect dependence plots for logP and pI (d), logP and net.charge (e), and pI and net.charge (f). The color gradient indicates the interaction effect magnitude for pI (d), net.charge (e), and net.charge (f), respectively. Black dashed lines indicate where SHAP effect values are zero, showing the direction and strength of feature interaction impacts on the model's predictions.

**Table S1.** Performance analysis of different machine learning algorithms (RF, SVM, Transformer) on a fixed testing dataset of 10,000 samples. This table summarizes the training sizes at which the key performance metrics ($R^2$, MAE, MSE, RMSE) reach their best performance points and inflection points during the learning curve.

| algorithm | train_size | value | point_type | properties | metric |
|---|---|---|---|---|---|
| RF | 15000 | 0.821 | Best | AP | R2_TEST |
| RF | 2500 | 0.741 | Inflection | AP | R2_TEST |
| SVM | 15000 | 0.901 | Best | AP | R2_TEST |
| SVM | 10500 | 0.864 | Inflection | AP | R2_TEST |
| Transformer | 16000 | 0.945 | Best | AP | R2_TEST |
| Transformer | 3500 | 0.910 | Inflection | AP | R2_TEST |
| RF | 15000 | 0.077 | Best | AP | MAE_TEST |
| RF | 1500 | 0.107 | Inflection | AP | MAE_TEST |
| SVM | 16000 | 0.061 | Best | AP | MAE_TEST |
| SVM | 10500 | 0.072 | Inflection | AP | MAE_TEST |
| Transformer | 16000 | 0.045 | Best | AP | MAE_TEST |
| Transformer | 3500 | 0.058 | Inflection | AP | MAE_TEST |
| RF | 15000 | 0.104 | Best | AP | RMSE_TEST |
| RF | 2500 | 0.126 | Inflection | AP | RMSE_TEST |
| SVM | 15000 | 0.078 | Best | AP | RMSE_TEST |
| SVM | 10500 | 0.091 | Inflection | AP | RMSE_TEST |
| Transformer | 16000 | 0.058 | Best | AP | RMSE_TEST |
| Transformer | 3500 | 0.074 | Inflection | AP | RMSE_TEST |
| RF | 15000 | 0.011 | Best | AP | MSE_TEST |
| RF | 2500 | 0.016 | Inflection | AP | MSE_TEST |
| SVM | 15000 | 0.006 | Best | AP | MSE_TEST |
| SVM | 10500 | 0.008 | Inflection | AP | MSE_TEST |
| Transformer | 13500 | 0.003 | Best | AP | MSE_TEST |
| Transformer | 14000 | 0.003 | Best | AP | MSE_TEST |
| Transformer | 14500 | 0.003 | Best | AP | MSE_TEST |
| Transformer | 15000 | 0.003 | Best | AP | MSE_TEST |
| Transformer | 15500 | 0.003 | Best | AP | MSE_TEST |
| Transformer | 16000 | 0.003 | Best | AP | MSE_TEST |
| Transformer | 3500 | 0.005 | Inflection | AP | MSE_TEST |
| RF | 16000 | 0.929 | Best | logP | R2_TEST |
| RF | 2000 | 0.853 | Inflection | logP | R2_TEST |
| SVM | 16000 | 1.000 | Best | logP | R2_TEST |
| SVM | 8500 | 1.000 | Inflection | logP | R2_TEST |
| Transformer | 9000 | 0.998 | Best | logP | R2_TEST |
| Transformer | 4500 | 0.991 | Inflection | logP | R2_TEST |
| RF | 16000 | 0.610 | Best | logP | MAE_TEST |
| RF | 2000 | 0.903 | Inflection | logP | MAE_TEST |
| SVM | 11000 | 0.039 | Best | logP | MAE_TEST |
| SVM | 6000 | 0.040 | Inflection | logP | MAE_TEST |
| Transformer | 9000 | 0.099 | Best | logP | MAE_TEST |
| Transformer | 4500 | 0.221 | Inflection | logP | MAE_TEST |

| | | | | | |
|---|---|---|---|---|---|
| RF | 16000 | 0.826 | Best | logP | RMSE_TEST |
| RF | 2000 | 1.191 | Inflection | logP | RMSE_TEST |
| SVM | 16000 | 0.050 | Best | logP | RMSE_TEST |
| SVM | 8500 | 0.061 | Inflection | logP | RMSE_TEST |
| Transformer | 9000 | 0.131 | Best | logP | RMSE_TEST |
| Transformer | 4500 | 0.289 | Inflection | logP | RMSE_TEST |
| RF | 16000 | 0.682 | Best | logP | MSE_TEST |
| RF | 2000 | 1.418 | Inflection | logP | MSE_TEST |
| SVM | 16000 | 0.002 | Best | logP | MSE_TEST |
| SVM | 8500 | 0.004 | Inflection | logP | MSE_TEST |
| Transformer | 9000 | 0.017 | Best | logP | MSE_TEST |
| Transformer | 4500 | 0.084 | Inflection | logP | MSE_TEST |
| RF | 16000 | 0.967 | Best | pI | R2_TEST |
| RF | 2500 | 0.932 | Inflection | pI | R2_TEST |
| SVM | 16000 | 0.974 | Best | pI | R2_TEST |
| SVM | 2000 | 0.935 | Inflection | pI | R2_TEST |
| Transformer | 15000 | 0.993 | Best | pI | R2_TEST |
| Transformer | 4500 | 0.985 | Inflection | pI | R2_TEST |
| RF | 15500 | 0.268 | Best | pI | MAE_TEST |
| RF | 2500 | 0.395 | Inflection | pI | MAE_TEST |
| SVM | 15500 | 0.241 | Best | pI | MAE_TEST |
| SVM | 1500 | 0.445 | Inflection | pI | MAE_TEST |
| Transformer | 15000 | 0.111 | Best | pI | MAE_TEST |
| Transformer | 4500 | 0.177 | Inflection | pI | MAE_TEST |
| RF | 16000 | 0.366 | Best | pI | RMSE_TEST |
| RF | 2500 | 0.524 | Inflection | pI | RMSE_TEST |
| SVM | 16000 | 0.321 | Best | pI | RMSE_TEST |
| SVM | 2000 | 0.511 | Inflection | pI | RMSE_TEST |
| Transformer | 15000 | 0.162 | Best | pI | RMSE_TEST |
| Transformer | 4500 | 0.244 | Inflection | pI | RMSE_TEST |
| RF | 16000 | 0.134 | Best | pI | MSE_TEST |
| RF | 2500 | 0.274 | Inflection | pI | MSE_TEST |
| SVM | 16000 | 0.103 | Best | pI | MSE_TEST |
| SVM | 2000 | 0.261 | Inflection | pI | MSE_TEST |
| Transformer | 15000 | 0.026 | Best | pI | MSE_TEST |
| Transformer | 4500 | 0.060 | Inflection | pI | MSE_TEST |

**Table S2.** Performance analysis of different AI algorithms (RF, SVM, Transformer) on a non-fixed testing dataset. This table summarizes the training sizes at which the key performance metrics ($R^2$, MAE, MSE, RMSE) reach their best performance points and inflection points during the learning curve. The results are derived from evaluations on a varying test set, in contrast to the fixed testing dataset used in Table 1.

| algorithm | train_size | value | point_type | properties | metric |
|---|---|---|---|---|---|
| RF | 16000 | 0.821 | Best | AP | R2_TEST |
| RF | 2500 | 0.744 | Inflection | AP | R2_TEST |
| SVM | 15000 | 0.902 | Best | AP | R2_TEST |
| SVM | 10500 | 0.869 | Inflection | AP | R2_TEST |
| Transformer | 15500 | 0.946 | Best | AP | R2_TEST |
| Transformer | 3500 | 0.910 | Inflection | AP | R2_TEST |
| RF | 15500 | 0.079 | Best | AP | MAE_TEST |
| RF | 16000 | 0.079 | Best | AP | MAE_TEST |
| RF | 2500 | 0.096 | Inflection | AP | MAE_TEST |
| SVM | 15000 | 0.061 | Best | AP | MAE_TEST |
| SVM | 10500 | 0.071 | Inflection | AP | MAE_TEST |
| Transformer | 15500 | 0.045 | Best | AP | MAE_TEST |
| Transformer | 16000 | 0.045 | Best | AP | MAE_TEST |
| Transformer | 3500 | 0.059 | Inflection | AP | MAE_TEST |
| RF | 16000 | 0.106 | Best | AP | RMSE_TEST |
| RF | 2500 | 0.127 | Inflection | AP | RMSE_TEST |
| SVM | 15000 | 0.078 | Best | AP | RMSE_TEST |
| SVM | 10500 | 0.090 | Inflection | AP | RMSE_TEST |
| Transformer | 15500 | 0.058 | Best | AP | RMSE_TEST |
| Transformer | 3500 | 0.075 | Inflection | AP | RMSE_TEST |
| RF | 16000 | 0.011 | Best | AP | MSE_TEST |
| RF | 2500 | 0.016 | Inflection | AP | MSE_TEST |
| SVM | 15000 | 0.006 | Best | AP | MSE_TEST |
| SVM | 15500 | 0.006 | Best | AP | MSE_TEST |
| SVM | 16000 | 0.006 | Best | AP | MSE_TEST |
| SVM | 10500 | 0.008 | Inflection | AP | MSE_TEST |
| Transformer | 15000 | 0.003 | Best | AP | MSE_TEST |
| Transformer | 15500 | 0.003 | Best | AP | MSE_TEST |
| Transformer | 16000 | 0.003 | Best | AP | MSE_TEST |
| Transformer | 3500 | 0.006 | Inflection | AP | MSE_TEST |
| RF | 16000 | 0.930 | Best | logP | R2_TEST |
| RF | 2000 | 0.855 | Inflection | logP | R2_TEST |
| SVM | 16000 | 1.000 | Best | logP | R2_TEST |
| SVM | 8500 | 1.000 | Inflection | logP | R2_TEST |
| Transformer | 9000 | 0.998 | Best | logP | R2_TEST |
| Transformer | 4500 | 0.991 | Inflection | logP | R2_TEST |
| RF | 16000 | 0.615 | Best | logP | MAE_TEST |
| RF | 2000 | 0.906 | Inflection | logP | MAE_TEST |
| SVM | 11000 | 0.039 | Best | logP | MAE_TEST |
| SVM | 6000 | 0.040 | Inflection | logP | MAE_TEST |

| | | | | | |
|---|---|---|---|---|---|
| Transformer | 9000 | 0.099 | Best | logP | MAE_TEST |
| Transformer | 4500 | 0.222 | Inflection | logP | MAE_TEST |
| RF | 16000 | 0.827 | Best | logP | RMSE_TEST |
| RF | 2000 | 1.190 | Inflection | logP | RMSE_TEST |
| SVM | 16000 | 0.050 | Best | logP | RMSE_TEST |
| SVM | 8500 | 0.061 | Inflection | logP | RMSE_TEST |
| Transformer | 5500 | 0.124 | Best | logP | RMSE_TEST |
| Transformer | 4500 | 0.289 | Inflection | logP | RMSE_TEST |
| RF | 16000 | 0.684 | Best | logP | MSE_TEST |
| RF | 2000 | 1.415 | Inflection | logP | MSE_TEST |
| SVM | 16000 | 0.002 | Best | logP | MSE_TEST |
| SVM | 5500 | 0.004 | Inflection | logP | MSE_TEST |
| Transformer | 9000 | 0.017 | Best | logP | MSE_TEST |
| Transformer | 4500 | 0.084 | Inflection | logP | MSE_TEST |
| RF | 16000 | 0.967 | Best | pI | R2_TEST |
| RF | 2500 | 0.932 | Inflection | pI | R2_TEST |
| SVM | 16000 | 0.974 | Best | pI | R2_TEST |
| SVM | 2000 | 0.935 | Inflection | pI | R2_TEST |
| Transformer | 15000 | 0.994 | Best | pI | R2_TEST |
| Transformer | 4500 | 0.985 | Inflection | pI | R2_TEST |
| RF | 15500 | 0.266 | Best | pI | MAE_TEST |
| RF | 2500 | 0.391 | Inflection | pI | MAE_TEST |
| SVM | 16000 | 0.239 | Best | pI | MAE_TEST |
| SVM | 1500 | 0.442 | Inflection | pI | MAE_TEST |
| Transformer | 15000 | 0.110 | Best | pI | MAE_TEST |
| Transformer | 4500 | 0.174 | Inflection | pI | MAE_TEST |
| RF | 16000 | 0.362 | Best | pI | RMSE_TEST |
| RF | 2500 | 0.519 | Inflection | pI | RMSE_TEST |
| SVM | 16000 | 0.320 | Best | pI | RMSE_TEST |
| SVM | 2000 | 0.507 | Inflection | pI | RMSE_TEST |
| Transformer | 15000 | 0.161 | Best | pI | RMSE_TEST |
| Transformer | 4500 | 0.242 | Inflection | pI | RMSE_TEST |
| RF | 16000 | 0.131 | Best | pI | MSE_TEST |
| RF | 2500 | 0.269 | Inflection | pI | MSE_TEST |
| SVM | 16000 | 0.102 | Best | pI | MSE_TEST |
| SVM | 2000 | 0.257 | Inflection | pI | MSE_TEST |
| Transformer | 15000 | 0.026 | Best | pI | MSE_TEST |
| Transformer | 4500 | 0.058 | Inflection | pI | MSE_TEST |

**Table S3.** Performance evaluation of three machine learning models (XGBoost, LightGBM, CatBoost) after feature selection, where 11 key features were identified from an initial pool of 725 variables. The table presents various commonly performance metrics for each model, including accuracy, precision, recall, F1 score, ROC AUC, and PR AUC.

| **Model** | **Accuracy** | **Precision** | **Recall** | **F1 Score** | **ROC AUC** | **PR AUC** | **Parameters** |
|---|---|---|---|---|---|---|---|
| **XGBoost** | 0.879 | 0.879 | 0.879 | 0.878 | 0.969 | 0.934 | 'n_estimators': 400, 'max_depth': 6, 'learning_rate': 0.1 |
| **LightGBM** | 0.885 | 0.884 | 0.885 | 0.884 | 0.971 | 0.941 | 'n_estimators': 400, 'max_depth': 6, 'learning_rate': 0.1 |
| **CatBoost** | 0.866 | 0.865 | 0.866 | 0.864 | 0.962 | 0.91 | 'learning_rate': 0.1, 'iterations': 300, 'depth': 6 |

**Table S4.** Hyperparameter Search Spaces for Random Forest and Support Vector Machine Models.

| **Model** | **Hyperparameter** | **Possible Values** |
|---|---|---|
| **Random Forest (RF)** | n_estimators | [100, 200, 300, 400, 500, 600, 700, 800, 900, 1000] |
| | max_features | ['sqrt', 'log2'] |
| | max_depth | [None, 10, 20, 30, 40, 50] |
| | min_samples_split | [2, 5, 10] |
| | min_samples_leaf | [1, 2, 4] |
| | bootstrap | [True, False] |
| | min_weight_fraction_leaf | [0.0, 0.1, 0.2] |
| | max_leaf_nodes | [None, 10, 20, 30] |
| | min_impurity_decrease | [0.0, 0.01, 0.02] |
| **Support Vector Machine (SVM)** | kernel | ['linear', 'poly', 'rbf'] |
| | C | Uniform(0.001, 100) |
| | degree | randint(1, 5) |
| | gamma | ['scale', 'auto'] |

**Table S5.** Hyperparameter Search Spaces for Tree-based Models (XGBoost, LightGBM, CatBoost).

| Model | Hyperparameter | Possible Values |
|---|---|---|
| | number of trees | [100, 200, 300, 400, 500] |
| **XGBoost** | tree depth | [3, 4, 5, 6, 7] |
| | learning rate | [0.01, 0.02, 0.05, 0.1] |
| | number of trees | [100, 200, 300, 400, 500] |
| **LightGBM** | tree depth | [3, 4, 5, 6, 7] |
| | learning rate | [0.01, 0.02, 0.05, 0.1] |
| | number of trees | [100, 200, 300] |
| **CatBoost** | tree depth | [3, 4, 5, 6, 7] |
| | learning rate | [0.01, 0.02, 0.05, 0.1] |
