## Supplementary material for "Harnessing Uniform Design to Enhance AI-Driven Predictions of Physicochemical Properties of Short Peptides": List of Abbreviations and Acronyms

This document provides a comprehensive list of abbreviations and acronyms used in the main text and supplementary materials of the manuscript.

**Table 1. List of abbreviations and acronyms used in the main text and supplementary materials**

| Abbreviation | Full Term |
|---|---|
| AI | Artificial intelligence |
| AP | Aggregation propensity |
| CatBoost | Categorical Boosting |
| CGMD | Coarse-grained molecular dynamics |
| DL | Deep learning |
| LightGBM | Light Gradient Boosting Machine |
| logP | Hydrophilicity |
| LOWESS | Locally weighted scatterplot smoothing |
| MAE | Mean absolute error |
| MD | Molecular dynamics |
| MD2 | Mixture L2-Discrepancy |
| ML | Machine learning |
| MSE | Mean square error |
| PR AUC | Precision-Recall area under the curve |
| pI | Isoelectric point |
| $R^2$ | Coefficient of determination |
| RF | Random forest |
| RMSE | Root mean square error |
| ROC AUC | Receiver operating characteristic area under the curve |
| SHAP | Shapley additive explanations |
| SOTA | Stochastic optimization adaptive threshold acceptance |
| SVM | Support vector machine |
| TA | Threshold acceptance |
| UD | Uniform design |
| XGBoost | eXtreme Gradient Boosting |

**Table 2. List of variable name abbreviations and descriptions used in Supplementary Data 10**

| Variable | Full Variable Name | Description |
|---|---|---|
| A | Alanine | The fraction of alanine within the tetrapeptide sequence. |
| R | Arginine | The fraction of arginine within the tetrapeptide sequence. |
| N | Asparagine | The fraction of asparagine within the tetrapeptide sequence. |
| D | Aspartic acid | The fraction of aspartic acid within the tetrapeptide sequence. |
| C | Cysteine | The fraction of cysteine within the tetrapeptide sequence. |
| E | Glutamic acid | The fraction of glutamic acid within the tetrapeptide sequence. |
| Q | Glutamine | The fraction of glutamine within the tetrapeptide sequence. |
| G | Glycine | The fraction of glycine within the tetrapeptide sequence. |
| H | Histidine | The fraction of histidine within the tetrapeptide sequence. |
| I | Isoleucine | The fraction of isoleucine within the tetrapeptide sequence. |
| L | Leucine | The fraction of leucine within the tetrapeptide sequence. |
| K | Lysine | The fraction of lysine within the tetrapeptide sequence. |
| M | Methionine | The fraction of methionine within the tetrapeptide sequence. |
| F | Phenylalanine | The fraction of phenylalanine within the |

| | | |
|---|---|---|
| | | tetrapeptide sequence. |
| P | Proline | The fraction of proline within the tetrapeptide sequence. |
| S | Serine | The fraction of serine within the tetrapeptide sequence. |
| T | Threonine | The fraction of threonine within the tetrapeptide sequence. |
| W | Tryptophan | The fraction of tryptophan within the tetrapeptide sequence. |
| Y | Tyrosine | The fraction of tyrosine within the tetrapeptide sequence. |
| V | Valine | The fraction of valine within the tetrapeptide sequence. |
| AA | Alanine-Alanine | The fraction of the dipeptide "Alanine-Alanine" in the sequence. |
| RA | Arginine-Alanine | The fraction of the dipeptide "Arginine-Alanine" in the sequence. |
| NA | Asparagine-Alanine | The fraction of the dipeptide "Asparagine-Alanine" in the sequence. |
| DA | Aspartic acid-Alanine | The fraction of the dipeptide "Aspartic acid-Alanine" in the sequence. |
| CA | Cysteine-Alanine | The fraction of the dipeptide "Cysteine-Alanine" in the sequence. |
| EA | Glutamic acid-Alanine | The fraction of the dipeptide "Glutamic acid-Alanine" in the sequence. |
| QA | Glutamine-Alanine | The fraction of the |

| | | |
|---|---|---|
| | | dipeptide "Glutamine-Alanine" in the sequence. |
| GA | Glycine-Alanine | The fraction of the dipeptide "Glycine-Alanine" in the sequence. |
| HA | Histidine-Alanine | The fraction of the dipeptide "Histidine-Alanine" in the sequence. |
| IA | Isoleucine-Alanine | The fraction of the dipeptide "Isoleucine-Alanine" in the sequence. |
| LA | Leucine-Alanine | The fraction of the dipeptide "Leucine-Alanine" in the sequence. |
| KA | Lysine-Alanine | The fraction of the dipeptide "Lysine-Alanine" in the sequence. |
| MA | Methionine-Alanine | The fraction of the dipeptide "Methionine-Alanine" in the sequence. |
| FA | Phenylalanine-Alanine | The fraction of the dipeptide "Phenylalanine-Alanine" in the sequence. |
| PA | Proline-Alanine | The fraction of the dipeptide "Proline-Alanine" in the sequence. |
| SA | Serine-Alanine | The fraction of the dipeptide "Serine-Alanine" in the sequence. |
| TA | Threonine-Alanine | The fraction of the dipeptide "Threonine-Alanine" in the sequence. |
| WA | Tryptophan-Alanine | The fraction of the |

| | | |
|---|---|---|
| | | dipeptide "Tryptophan-Alanine" in the sequence. |
| YA | Tyrosine-Alanine | The fraction of the dipeptide "Tyrosine-Alanine" in the sequence. |
| VA | Valine-Alanine | The fraction of the dipeptide "Valine-Alanine" in the sequence. |
| AR | Alanine-Arginine | The fraction of the dipeptide "Alanine-Arginine" in the sequence. |
| RR | Arginine-Arginine | The fraction of the dipeptide "Arginine-Arginine" in the sequence. |
| NR | Asparagine-Arginine | The fraction of the dipeptide "Asparagine-Arginine" in the sequence. |
| DR | Aspartic acid-Arginine | The fraction of the dipeptide "Aspartic acid-Arginine" in the sequence. |
| CR | Cysteine-Arginine | The fraction of the dipeptide "Cysteine-Arginine" in the sequence. |
| ER | Glutamic acid-Arginine | The fraction of the dipeptide "Glutamic acid-Arginine" in the sequence. |
| QR | Glutamine-Arginine | The fraction of the dipeptide "Glutamine-Arginine" in the sequence. |
| GR | Glycine-Arginine | The fraction of the dipeptide "Glycine-Arginine" in the sequence. |
| HR | Histidine-Arginine | The fraction of the |

| | | |
|---|---|---|
| | | dipeptide "Histidine-Arginine" in the sequence. |
| IR | Isoleucine-Arginine | The fraction of the dipeptide "Isoleucine-Arginine" in the sequence. |
| LR | Leucine-Arginine | The fraction of the dipeptide "Leucine-Arginine" in the sequence. |
| KR | Lysine-Arginine | The fraction of the dipeptide "Lysine-Arginine" in the sequence. |
| MR | Methionine-Arginine | The fraction of the dipeptide "Methionine-Arginine" in the sequence. |
| FR | Phenylalanine-Arginine | The fraction of the dipeptide "Phenylalanine-Arginine" in the sequence. |
| PR | Proline-Arginine | The fraction of the dipeptide "Proline-Arginine" in the sequence. |
| SR | Serine-Arginine | The fraction of the dipeptide "Serine-Arginine" in the sequence. |
| TR | Threonine-Arginine | The fraction of the dipeptide "Threonine-Arginine" in the sequence. |
| WR | Tryptophan-Arginine | The fraction of the dipeptide "Tryptophan-Arginine" in the sequence. |
| YR | Tyrosine-Arginine | The fraction of the dipeptide "Tyrosine-Arginine" in the sequence. |
| VR | Valine-Arginine | The fraction of the |

| | | |
|---|---|---|
| | | dipeptide "Valine-Arginine" in the sequence. |
| AN | Alanine-Asparagine | The fraction of the dipeptide "Alanine-Asparagine" in the sequence. |
| RN | Arginine-Asparagine | The fraction of the dipeptide "Arginine-Asparagine" in the sequence. |
| NN | Asparagine-Asparagine | The fraction of the dipeptide "Asparagine-Asparagine" in the sequence. |
| DN | Aspartic acid-Asparagine | The fraction of the dipeptide "Aspartic acid-Asparagine" in the sequence. |
| CN | Cysteine-Asparagine | The fraction of the dipeptide "Cysteine-Asparagine" in the sequence. |
| EN | Glutamic acid-Asparagine | The fraction of the dipeptide "Glutamic acid-Asparagine" in the sequence. |
| QN | Glutamine-Asparagine | The fraction of the dipeptide "Glutamine-Asparagine" in the sequence. |
| GN | Glycine-Asparagine | The fraction of the dipeptide "Glycine-Asparagine" in the sequence. |
| HN | Histidine-Asparagine | The fraction of the dipeptide "Histidine-Asparagine" in the sequence. |
| IN | Isoleucine-Asparagine | The fraction of the dipeptide "Isoleucine-Asparagine" in the sequence. |
| LN | Leucine-Asparagine | The fraction of the |

| | | |
|---|---|---|
| | | dipeptide "Leucine-Asparagine" in the sequence. |
| KN | Lysine-Asparagine | The fraction of the dipeptide "Lysine-Asparagine" in the sequence. |
| MN | Methionine-Asparagine | The fraction of the dipeptide "Methionine-Asparagine" in the sequence. |
| FN | Phenylalanine-Asparagi ne | The fraction of the dipeptide "Phenylalanine-Asparagin e" in the sequence. |
| PN | Proline-Asparagine | The fraction of the dipeptide "Proline-Asparagine" in the sequence. |
| SN | Serine-Asparagine | The fraction of the dipeptide "Serine-Asparagine" in the sequence. |
| TN | Threonine-Asparagine | The fraction of the dipeptide "Threonine-Asparagine" in the sequence. |
| WN | Tryptophan-Asparagine | The fraction of the dipeptide "Tryptophan-Asparagine" in the sequence. |
| YN | Tyrosine-Asparagine | The fraction of the dipeptide "Tyrosine-Asparagine" in the sequence. |
| VN | Valine-Asparagine | The fraction of the dipeptide "Valine-Asparagine" in the sequence. |
| AD | Alanine-Aspartic acid | The fraction of the dipeptide "Alanine-Aspartic acid" in the sequence. |
| RD | Arginine-Aspartic acid | The fraction of the |

| | | |
|---|---|---|
| | | dipeptide "Arginine-Aspartic acid" in the sequence. |
| ND | Asparagine-Aspartic acid | The fraction of the dipeptide "Asparagine-Aspartic acid" in the sequence. |
| DD | Aspartic acid-Aspartic acid | The fraction of the dipeptide "Aspartic acid-Aspartic acid" in the sequence. |
| CD | Cysteine-Aspartic acid | The fraction of the dipeptide "Cysteine-Aspartic acid" in the sequence. |
| ED | Glutamic acid-Aspartic acid | The fraction of the dipeptide "Glutamic acid-Aspartic acid" in the sequence. |
| QD | Glutamine-Aspartic acid | The fraction of the dipeptide "Glutamine-Aspartic acid" in the sequence. |
| GD | Glycine-Aspartic acid | The fraction of the dipeptide "Glycine-Aspartic acid" in the sequence. |
| HD | Histidine-Aspartic acid | The fraction of the dipeptide "Histidine-Aspartic acid" in the sequence. |
| ID | Isoleucine-Aspartic acid | The fraction of the dipeptide "Isoleucine-Aspartic acid" in the sequence. |
| LD | Leucine-Aspartic acid | The fraction of the dipeptide "Leucine-Aspartic acid" in the sequence. |
| KD | Lysine-Aspartic acid | The fraction of the dipeptide "Lysine-Aspartic acid" in the sequence. |
| MD | Methionine-Aspartic acid | The fraction of the dipeptide |

| | | |
|---|---|---|
| | | "Methionine-Aspartic acid" in the sequence. |
| FD | Phenylalanine-Aspartic acid | The fraction of the dipeptide "Phenylalanine-Aspartic acid" in the sequence. |
| PD | Proline-Aspartic acid | The fraction of the dipeptide "Proline-Aspartic acid" in the sequence. |
| SD | Serine-Aspartic acid | The fraction of the dipeptide "Serine-Aspartic acid" in the sequence. |
| TD | Threonine-Aspartic acid | The fraction of the dipeptide "Threonine-Aspartic acid" in the sequence. |
| WD | Tryptophan-Aspartic acid | The fraction of the dipeptide "Tryptophan-Aspartic acid" in the sequence. |
| YD | Tyrosine-Aspartic acid | The fraction of the dipeptide "Tyrosine-Aspartic acid" in the sequence. |
| VD | Valine-Aspartic acid | The fraction of the dipeptide "Valine-Aspartic acid" in the sequence. |
| AC | Alanine-Cysteine | The fraction of the dipeptide "Alanine-Cysteine" in the sequence. |
| RC | Arginine-Cysteine | The fraction of the dipeptide "Arginine-Cysteine" in the sequence. |
| NC | Asparagine-Cysteine | The fraction of the dipeptide "Asparagine-Cysteine" in the sequence. |
| DC | Aspartic acid-Cysteine | The fraction of the dipeptide "Aspartic acid-Cysteine" in the sequence. |

| | | |
|---|---|---|
| CC | Cysteine-Cysteine | The fraction of the dipeptide "Cysteine-Cysteine" in the sequence. |
| EC | Glutamic acid-Cysteine | The fraction of the dipeptide "Glutamic acid-Cysteine" in the sequence. |
| QC | Glutamine-Cysteine | The fraction of the dipeptide "Glutamine-Cysteine" in the sequence. |
| GC | Glycine-Cysteine | The fraction of the dipeptide "Glycine-Cysteine" in the sequence. |
| HC | Histidine-Cysteine | The fraction of the dipeptide "Histidine-Cysteine" in the sequence. |
| IC | Isoleucine-Cysteine | The fraction of the dipeptide "Isoleucine-Cysteine" in the sequence. |
| LC | Leucine-Cysteine | The fraction of the dipeptide "Leucine-Cysteine" in the sequence. |
| KC | Lysine-Cysteine | The fraction of the dipeptide "Lysine-Cysteine" in the sequence. |
| MC | Methionine-Cysteine | The fraction of the dipeptide "Methionine-Cysteine" in the sequence. |
| FC | Phenylalanine-Cysteine | The fraction of the dipeptide "Phenylalanine-Cysteine" in the sequence. |
| PC | Proline-Cysteine | The fraction of the dipeptide "Proline-Cysteine" in the sequence. |

| | | |
|---|---|---|
| SC | Serine-Cysteine | The fraction of the dipeptide "Serine-Cysteine" in the sequence. |
| TC | Threonine-Cysteine | The fraction of the dipeptide "Threonine-Cysteine" in the sequence. |
| WC | Tryptophan-Cysteine | The fraction of the dipeptide "Tryptophan-Cysteine" in the sequence. |
| YC | Tyrosine-Cysteine | The fraction of the dipeptide "Tyrosine-Cysteine" in the sequence. |
| VC | Valine-Cysteine | The fraction of the dipeptide "Valine-Cysteine" in the sequence. |
| AE | Alanine-Glutamic acid | The fraction of the dipeptide "Alanine-Glutamic acid" in the sequence. |
| RE | Arginine-Glutamic acid | The fraction of the dipeptide "Arginine-Glutamic acid" in the sequence. |
| NE | Asparagine-Glutamic acid | The fraction of the dipeptide "Asparagine-Glutamic acid" in the sequence. |
| DE | Aspartic acid-Glutamic acid | The fraction of the dipeptide "Aspartic acid-Glutamic acid" in the sequence. |
| CE | Cysteine-Glutamic acid | The fraction of the dipeptide "Cysteine-Glutamic acid" in the sequence. |
| EE | Glutamic acid-Glutamic acid | The fraction of the dipeptide "Glutamic acid-Glutamic acid" in the sequence. |

| | | |
|---|---|---|
| QE | Glutamine-Glutamic acid | The fraction of the dipeptide "Glutamine-Glutamic acid" in the sequence. |
| GE | Glycine-Glutamic acid | The fraction of the dipeptide "Glycine-Glutamic acid" in the sequence. |
| HE | Histidine-Glutamic acid | The fraction of the dipeptide "Histidine-Glutamic acid" in the sequence. |
| IE | Isoleucine-Glutamic acid | The fraction of the dipeptide "Isoleucine-Glutamic acid" in the sequence. |
| LE | Leucine-Glutamic acid | The fraction of the dipeptide "Leucine-Glutamic acid" in the sequence. |
| KE | Lysine-Glutamic acid | The fraction of the dipeptide "Lysine-Glutamic acid" in the sequence. |
| ME | Methionine-Glutamic acid | The fraction of the dipeptide "Methionine-Glutamic acid" in the sequence. |
| FE | Phenylalanine-Glutamic acid | The fraction of the dipeptide "Phenylalanine-Glutamic acid" in the sequence. |
| PE | Proline-Glutamic acid | The fraction of the dipeptide "Proline-Glutamic acid" in the sequence. |
| SE | Serine-Glutamic acid | The fraction of the dipeptide "Serine-Glutamic acid" in the sequence. |
| TE | Threonine-Glutamic acid | The fraction of the dipeptide "Threonine-Glutamic acid" in the sequence. |

| | | |
|---|---|---|
| WE | Tryptophan-Glutamic acid | The fraction of the dipeptide "Tryptophan-Glutamic acid" in the sequence. |
| YE | Tyrosine-Glutamic acid | The fraction of the dipeptide "Tyrosine-Glutamic acid" in the sequence. |
| VE | Valine-Glutamic acid | The fraction of the dipeptide "Valine-Glutamic acid" in the sequence. |
| GE | Glycine-Glutamic acid | The fraction of the dipeptide "Glycine-Glutamic acid" in the sequence. |
| HE | Histidine-Glutamic acid | The fraction of the dipeptide "Histidine-Glutamic acid" in the sequence. |
| IE | Isoleucine-Glutamic acid | The fraction of the dipeptide "Isoleucine-Glutamic acid" in the sequence. |
| LE | Leucine-Glutamic acid | The fraction of the dipeptide "Leucine-Glutamic acid" in the sequence. |
| KE | Lysine-Glutamic acid | The fraction of the dipeptide "Lysine-Glutamic acid" in the sequence. |
| ME | Methionine-Glutamic acid | The fraction of the dipeptide "Methionine-Glutamic acid" in the sequence. |
| FE | Phenylalanine-Glutamic acid | The fraction of the dipeptide "Phenylalanine-Glutamic acid" in the sequence. |
| PE | Proline-Glutamic acid | The fraction of the dipeptide "Proline-Glutamic acid" in the sequence. |

| | | |
|---|---|---|
| SE | Serine-Glutamic acid | The fraction of the dipeptide "Serine-Glutamic acid" in the sequence. |
| TE | Threonine-Glutamic acid | The fraction of the dipeptide "Threonine-Glutamic acid" in the sequence. |
| WE | Tryptophan-Glutamic acid | The fraction of the dipeptide "Tryptophan-Glutamic acid" in the sequence. |
| YE | Tyrosine-Glutamic acid | The fraction of the dipeptide "Tyrosine-Glutamic acid" in the sequence. |
| VE | Valine-Glutamic acid | The fraction of the dipeptide "Valine-Glutamic acid" in the sequence. |
| AQ | Alanine-Glutamine | The fraction of the dipeptide "Alanine-Glutamine" in the sequence. |
| RQ | Arginine-Glutamine | The fraction of the dipeptide "Arginine-Glutamine" in the sequence. |
| NQ | Asparagine-Glutamine | The fraction of the dipeptide "Asparagine-Glutamine" in the sequence. |
| DQ | Aspartic acid-Glutamine | The fraction of the dipeptide "Aspartic acid-Glutamine" in the sequence. |
| CQ | Cysteine-Glutamine | The fraction of the dipeptide "Cysteine-Glutamine" in the sequence. |
| EQ | Glutamic acid-Glutamine | The fraction of the dipeptide "Glutamic acid-Glutamine" in the sequence. |

| | | |
|---|---|---|
| QQ | Glutamine-Glutamine | The fraction of the dipeptide "Glutamine-Glutamine" in the sequence. |
| GQ | Glycine-Glutamine | The fraction of the dipeptide "Glycine-Glutamine" in the sequence. |
| HQ | Histidine-Glutamine | The fraction of the dipeptide "Histidine-Glutamine" in the sequence. |
| IQ | Isoleucine-Glutamine | The fraction of the dipeptide "Isoleucine-Glutamine" in the sequence. |
| LQ | Leucine-Glutamine | The fraction of the dipeptide "Leucine-Glutamine" in the sequence. |
| KQ | Lysine-Glutamine | The fraction of the dipeptide "Lysine-Glutamine" in the sequence. |
| MQ | Methionine-Glutamine | The fraction of the dipeptide "Methionine-Glutamine" in the sequence. |
| FQ | Phenylalanine-Glutamine | The fraction of the dipeptide "Phenylalanine-Glutamine" in the sequence. |
| PQ | Proline-Glutamine | The fraction of the dipeptide "Proline-Glutamine" in the sequence. |
| SQ | Serine-Glutamine | The fraction of the dipeptide "Serine-Glutamine" in the sequence. |
| TQ | Threonine-Glutamine | The fraction of the dipeptide "Threonine-Glutamine" in the sequence. |

| | | |
|---|---|---|
| WQ | Tryptophan-Glutamine | The fraction of the dipeptide "Tryptophan-Glutamine" in the sequence. |
| YQ | Tyrosine-Glutamine | The fraction of the dipeptide "Tyrosine-Glutamine" in the sequence. |
| VQ | Valine-Glutamine | The fraction of the dipeptide "Valine-Glutamine" in the sequence. |
| AG | Alanine-Glycine | The fraction of the dipeptide "Alanine-Glycine" in the sequence. |
| RG | Arginine-Glycine | The fraction of the dipeptide "Arginine-Glycine" in the sequence. |
| NG | Asparagine-Glycine | The fraction of the dipeptide "Asparagine-Glycine" in the sequence. |
| DG | Aspartic acid-Glycine | The fraction of the dipeptide "Aspartic acid-Glycine" in the sequence. |
| CG | Cysteine-Glycine | The fraction of the dipeptide "Cysteine-Glycine" in the sequence. |
| EG | Glutamic acid-Glycine | The fraction of the dipeptide "Glutamic acid-Glycine" in the sequence. |
| QG | Glutamine-Glycine | The fraction of the dipeptide "Glutamine-Glycine" in the sequence. |
| GG | Glycine-Glycine | The fraction of the dipeptide "Glycine-Glycine" in the sequence. |

| | | |
|---|---|---|
| HG | Histidine-Glycine | The fraction of the dipeptide "Histidine-Glycine" in the sequence. |
| IG | Isoleucine-Glycine | The fraction of the dipeptide "Isoleucine-Glycine" in the sequence. |
| LG | Leucine-Glycine | The fraction of the dipeptide "Leucine-Glycine" in the sequence. |
| KG | Lysine-Glycine | The fraction of the dipeptide "Lysine-Glycine" in the sequence. |
| MG | Methionine-Glycine | The fraction of the dipeptide "Methionine-Glycine" in the sequence. |
| FG | Phenylalanine-Glycine | The fraction of the dipeptide "Phenylalanine-Glycine" in the sequence. |
| PG | Proline-Glycine | The fraction of the dipeptide "Proline-Glycine" in the sequence. |
| SG | Serine-Glycine | The fraction of the dipeptide "Serine-Glycine" in the sequence. |
| TG | Threonine-Glycine | The fraction of the dipeptide "Threonine-Glycine" in the sequence. |
| WG | Tryptophan-Glycine | The fraction of the dipeptide "Tryptophan-Glycine" in the sequence. |
| YG | Tyrosine-Glycine | The fraction of the dipeptide "Tyrosine-Glycine" in the sequence. |

| | | |
|---|---|---|
| VG | Valine-Glycine | The fraction of the dipeptide "Valine-Glycine" in the sequence. |
| AH | Alanine-Histidine | The fraction of the dipeptide "Alanine-Histidine" in the sequence. |
| RH | Arginine-Histidine | The fraction of the dipeptide "Arginine-Histidine" in the sequence. |
| NH | Asparagine-Histidine | The fraction of the dipeptide "Asparagine-Histidine" in the sequence. |
| DH | Aspartic acid-Histidine | The fraction of the dipeptide "Aspartic acid-Histidine" in the sequence. |
| CH | Cysteine-Histidine | The fraction of the dipeptide "Cysteine-Histidine" in the sequence. |
| EH | Glutamic acid-Histidine | The fraction of the dipeptide "Glutamic acid-Histidine" in the sequence. |
| QH | Glutamine-Histidine | The fraction of the dipeptide "Glutamine-Histidine" in the sequence. |
| GH | Glycine-Histidine | The fraction of the dipeptide "Glycine-Histidine" in the sequence. |
| HH | Histidine-Histidine | The fraction of the dipeptide "Histidine-Histidine" in the sequence. |
| IH | Isoleucine-Histidine | The fraction of the dipeptide "Isoleucine-Histidine" in the sequence. |

| | | |
|---|---|---|
| LH | Leucine-Histidine | The fraction of the dipeptide "Leucine-Histidine" in the sequence. |
| KH | Lysine-Histidine | The fraction of the dipeptide "Lysine-Histidine" in the sequence. |
| MH | Methionine-Histidine | The fraction of the dipeptide "Methionine-Histidine" in the sequence. |
| FH | Phenylalanine-Histidine | The fraction of the dipeptide "Phenylalanine-Histidine" in the sequence. |
| PH | Proline-Histidine | The fraction of the dipeptide "Proline-Histidine" in the sequence. |
| SH | Serine-Histidine | The fraction of the dipeptide "Serine-Histidine" in the sequence. |
| TH | Threonine-Histidine | The fraction of the dipeptide "Threonine-Histidine" in the sequence. |
| WH | Tryptophan-Histidine | The fraction of the dipeptide "Tryptophan-Histidine" in the sequence. |
| YH | Tyrosine-Histidine | The fraction of the dipeptide "Tyrosine-Histidine" in the sequence. |
| VH | Valine-Histidine | The fraction of the dipeptide "Valine-Histidine" in the sequence. |
| AI | Alanine-Isoleucine | The fraction of the dipeptide "Alanine-Isoleucine" in the sequence. |

| | | |
|---|---|---|
| RI | Arginine-Isoleucine | The fraction of the dipeptide "Arginine-Isoleucine" in the sequence. |
| NI | Asparagine-Isoleucine | The fraction of the dipeptide "Asparagine-Isoleucine" in the sequence. |
| DI | Aspartic acid-Isoleucine | The fraction of the dipeptide "Aspartic acid-Isoleucine" in the sequence. |
| CI | Cysteine-Isoleucine | The fraction of the dipeptide "Cysteine-Isoleucine" in the sequence. |
| EI | Glutamic acid-Isoleucine | The fraction of the dipeptide "Glutamic acid-Isoleucine" in the sequence. |
| QI | Glutamine-Isoleucine | The fraction of the dipeptide "Glutamine-Isoleucine" in the sequence. |
| GI | Glycine-Isoleucine | The fraction of the dipeptide "Glycine-Isoleucine" in the sequence. |
| HI | Histidine-Isoleucine | The fraction of the dipeptide "Histidine-Isoleucine" in the sequence. |
| II | Isoleucine-Isoleucine | The fraction of the dipeptide "Isoleucine-Isoleucine" in the sequence. |
| LI | Leucine-Isoleucine | The fraction of the dipeptide "Leucine-Isoleucine" in the sequence. |
| KI | Lysine-Isoleucine | The fraction of the dipeptide "Lysine-Isoleucine" in the sequence. |

| | | |
|---|---|---|
| MI | Methionine-Isoleucine | The fraction of the dipeptide "Methionine-Isoleucine" in the sequence. |
| FI | Phenylalanine-Isoleucine | The fraction of the dipeptide "Phenylalanine-Isoleucine" in the sequence. |
| PI | Proline-Isoleucine | The fraction of the dipeptide "Proline-Isoleucine" in the sequence. |
| SI | Serine-Isoleucine | The fraction of the dipeptide "Serine-Isoleucine" in the sequence. |
| TI | Threonine-Isoleucine | The fraction of the dipeptide "Threonine-Isoleucine" in the sequence. |
| WI | Tryptophan-Isoleucine | The fraction of the dipeptide "Tryptophan-Isoleucine" in the sequence. |
| YI | Tyrosine-Isoleucine | The fraction of the dipeptide "Tyrosine-Isoleucine" in the sequence. |
| VI | Valine-Isoleucine | The fraction of the dipeptide "Valine-Isoleucine" in the sequence. |
| AL | Alanine-Leucine | The fraction of the dipeptide "Alanine-Leucine" in the sequence. |
| RL | Arginine-Leucine | The fraction of the dipeptide "Arginine-Leucine" in the sequence. |
| NL | Asparagine-Leucine | The fraction of the dipeptide "Asparagine-Leucine" in the sequence. |

| | | |
|---|---|---|
| DL | Aspartic acid-Leucine | The fraction of the dipeptide "Aspartic acid-Leucine" in the sequence. |
| CL | Cysteine-Leucine | The fraction of the dipeptide "Cysteine-Leucine" in the sequence. |
| EL | Glutamic acid-Leucine | The fraction of the dipeptide "Glutamic acid-Leucine" in the sequence. |
| QL | Glutamine-Leucine | The fraction of the dipeptide "Glutamine-Leucine" in the sequence. |
| GL | Glycine-Leucine | The fraction of the dipeptide "Glycine-Leucine" in the sequence. |
| HL | Histidine-Leucine | The fraction of the dipeptide "Histidine-Leucine" in the sequence. |
| IL | Isoleucine-Leucine | The fraction of the dipeptide "Isoleucine-Leucine" in the sequence. |
| LL | Leucine-Leucine | The fraction of the dipeptide "Leucine-Leucine" in the sequence. |
| KL | Lysine-Leucine | The fraction of the dipeptide "Lysine-Leucine" in the sequence. |
| ML | Methionine-Leucine | The fraction of the dipeptide "Methionine-Leucine" in the sequence. |
| FL | Phenylalanine-Leucine | The fraction of the dipeptide "Phenylalanine-Leucine" in the sequence. |

| | | |
|---|---|---|
| PL | Proline-Leucine | The fraction of the dipeptide "Proline-Leucine" in the sequence. |
| SL | Serine-Leucine | The fraction of the dipeptide "Serine-Leucine" in the sequence. |
| TL | Threonine-Leucine | The fraction of the dipeptide "Threonine-Leucine" in the sequence. |
| WL | Tryptophan-Leucine | The fraction of the dipeptide "Tryptophan-Leucine" in the sequence. |
| YL | Tyrosine-Leucine | The fraction of the dipeptide "Tyrosine-Leucine" in the sequence. |
| VL | Valine-Leucine | The fraction of the dipeptide "Valine-Leucine" in the sequence. |
| AK | Alanine-Lysine | The fraction of the dipeptide "Alanine-Lysine" in the sequence. |
| RK | Arginine-Lysine | The fraction of the dipeptide "Arginine-Lysine" in the sequence. |
| NK | Asparagine-Lysine | The fraction of the dipeptide "Asparagine-Lysine" in the sequence. |
| DK | Aspartic acid-Lysine | The fraction of the dipeptide "Aspartic acid-Lysine" in the sequence. |
| CK | Cysteine-Lysine | The fraction of the dipeptide "Cysteine-Lysine" in the sequence. |

| | | |
|---|---|---|
| EK | Glutamic acid-Lysine | The fraction of the dipeptide "Glutamic acid-Lysine" in the sequence. |
| QK | Glutamine-Lysine | The fraction of the dipeptide "Glutamine-Lysine" in the sequence. |
| GK | Glycine-Lysine | The fraction of the dipeptide "Glycine-Lysine" in the sequence. |
| HK | Histidine-Lysine | The fraction of the dipeptide "Histidine-Lysine" in the sequence. |
| IK | Isoleucine-Lysine | The fraction of the dipeptide "Isoleucine-Lysine" in the sequence. |
| LK | Leucine-Lysine | The fraction of the dipeptide "Leucine-Lysine" in the sequence. |
| KK | Lysine-Lysine | The fraction of the dipeptide "Lysine-Lysine" in the sequence. |
| MK | Methionine-Lysine | The fraction of the dipeptide "Methionine-Lysine" in the sequence. |
| FK | Phenylalanine-Lysine | The fraction of the dipeptide "Phenylalanine-Lysine" in the sequence. |
| PK | Proline-Lysine | The fraction of the dipeptide "Proline-Lysine" in the sequence. |
| SK | Serine-Lysine | The fraction of the dipeptide "Serine-Lysine" in the sequence. |
| TK | Threonine-Lysine | The fraction of the dipeptide "Threonine-Lysine" in the |

| | | |
|---|---|---|
| | | sequence. |
| WK | Tryptophan-Lysine | The fraction of the dipeptide "Tryptophan-Lysine" in the sequence. |
| YK | Tyrosine-Lysine | The fraction of the dipeptide "Tyrosine-Lysine" in the sequence. |
| VK | Valine-Lysine | The fraction of the dipeptide "Valine-Lysine" in the sequence. |
| AM | Alanine-Methionine | The fraction of the dipeptide "Alanine-Methionine" in the sequence. |
| RM | Arginine-Methionine | The fraction of the dipeptide "Arginine-Methionine" in the sequence. |
| NM | Asparagine-Methionine | The fraction of the dipeptide "Asparagine-Methionine" in the sequence. |
| DM | Aspartic acid-Methionine | The fraction of the dipeptide "Aspartic acid-Methionine" in the sequence. |
| CM | Cysteine-Methionine | The fraction of the dipeptide "Cysteine-Methionine" in the sequence. |
| EM | Glutamic acid-Methionine | The fraction of the dipeptide "Glutamic acid-Methionine" in the sequence. |
| QM | Glutamine-Methionine | The fraction of the dipeptide "Glutamine-Methionine" in the sequence. |
| GM | Glycine-Methionine | The fraction of the dipeptide "Glycine-Methionine" in the sequence. |

| | | |
|---|---|---|
| HM | Histidine-Methionine | The fraction of the dipeptide "Histidine-Methionine" in the sequence. |
| IM | Isoleucine-Methionine | The fraction of the dipeptide "Isoleucine-Methionine" in the sequence. |
| LM | Leucine-Methionine | The fraction of the dipeptide "Leucine-Methionine" in the sequence. |
| KM | Lysine-Methionine | The fraction of the dipeptide "Lysine-Methionine" in the sequence. |
| MM | Methionine-Methionine | The fraction of the dipeptide "Methionine-Methionine" in the sequence. |
| FM | Phenylalanine-Methionine | The fraction of the dipeptide "Phenylalanine-Methionine" in the sequence. |
| PM | Proline-Methionine | The fraction of the dipeptide "Proline-Methionine" in the sequence. |
| SM | Serine-Methionine | The fraction of the dipeptide "Serine-Methionine" in the sequence. |
| TM | Threonine-Methionine | The fraction of the dipeptide "Threonine-Methionine" in the sequence. |
| WM | Tryptophan-Methionine | The fraction of the dipeptide "Tryptophan-Methionine" in the sequence. |
| YM | Tyrosine-Methionine | The fraction of the dipeptide "Tyrosine-Methionine" in the sequence. |

| | | |
|---|---|---|
| VM | Valine-Methionine | The fraction of the dipeptide "Valine-Methionine" in the sequence. |
| AF | Alanine-Phenylalanine | The fraction of the dipeptide "Alanine-Phenylalanine" in the sequence. |
| RF | Arginine-Phenylalanine | The fraction of the dipeptide "Arginine-Phenylalanine" in the sequence. |
| NF | Asparagine-Phenylalanine | The fraction of the dipeptide "Asparagine-Phenylalanine" in the sequence. |
| DF | Aspartic acid-Phenylalanine | The fraction of the dipeptide "Aspartic acid-Phenylalanine" in the sequence. |
| CF | Cysteine-Phenylalanine | The fraction of the dipeptide "Cysteine-Phenylalanine" in the sequence. |
| EF | Glutamic acid-Phenylalanine | The fraction of the dipeptide "Glutamic acid-Phenylalanine" in the sequence. |
| QF | Glutamine-Phenylalanine | The fraction of the dipeptide "Glutamine-Phenylalanine" in the sequence. |
| GF | Glycine-Phenylalanine | The fraction of the dipeptide "Glycine-Phenylalanine" in the sequence. |
| HF | Histidine-Phenylalanine | The fraction of the dipeptide "Histidine-Phenylalanine" in the sequence. |
| IF | Isoleucine-Phenylalanine | The fraction of the dipeptide "Isoleucine-Phenylalanine" in the sequence. |

| | | |
|---|---|---|
| LF | Leucine-Phenylalanine | The fraction of the dipeptide "Leucine-Phenylalanine" in the sequence. |
| KF | Lysine-Phenylalanine | The fraction of the dipeptide "Lysine-Phenylalanine" in the sequence. |
| MF | Methionine-Phenylalanine | The fraction of the dipeptide "Methionine-Phenylalanine" in the sequence. |
| FF | Phenylalanine-Phenylalanine | The fraction of the dipeptide "Phenylalanine-Phenylalanine" in the sequence. |
| PF | Proline-Phenylalanine | The fraction of the dipeptide "Proline-Phenylalanine" in the sequence. |
| SF | Serine-Phenylalanine | The fraction of the dipeptide "Serine-Phenylalanine" in the sequence. |
| TF | Threonine-Phenylalanine | The fraction of the dipeptide "Threonine-Phenylalanine" in the sequence. |
| WF | Tryptophan-Phenylalanine | The fraction of the dipeptide "Tryptophan-Phenylalanine" in the sequence. |
| YF | Tyrosine-Phenylalanine | The fraction of the dipeptide "Tyrosine-Phenylalanine" in the sequence. |
| VF | Valine-Phenylalanine | The fraction of the dipeptide "Valine-Phenylalanine" in the sequence. |
| AP | Alanine-Proline | The fraction of the dipeptide "Alanine-Proline" in the sequence. |

| | | |
|---|---|---|
| RP | Arginine-Proline | The fraction of the dipeptide "Arginine-Proline" in the sequence. |
| NP | Asparagine-Proline | The fraction of the dipeptide "Asparagine-Proline" in the sequence. |
| DP | Aspartic acid-Proline | The fraction of the dipeptide "Aspartic acid-Proline" in the sequence. |
| CP | Cysteine-Proline | The fraction of the dipeptide "Cysteine-Proline" in the sequence. |
| EP | Glutamic acid-Proline | The fraction of the dipeptide "Glutamic acid-Proline" in the sequence. |
| QP | Glutamine-Proline | The fraction of the dipeptide "Glutamine-Proline" in the sequence. |
| GP | Glycine-Proline | The fraction of the dipeptide "Glycine-Proline" in the sequence. |
| HP | Histidine-Proline | The fraction of the dipeptide "Histidine-Proline" in the sequence. |
| IP | Isoleucine-Proline | The fraction of the dipeptide "Isoleucine-Proline" in the sequence. |
| LP | Leucine-Proline | The fraction of the dipeptide "Leucine-Proline" in the sequence. |
| KP | Lysine-Proline | The fraction of the dipeptide "Lysine-Proline" in the sequence. |
| MP | Methionine-Proline | The fraction of the |

| | | |
|---|---|---|
| | | dipeptide "Methionine-Proline" in the sequence. |
| FP | Phenylalanine-Proline | The fraction of the dipeptide "Phenylalanine-Proline" in the sequence. |
| PP | Proline-Proline | The fraction of the dipeptide "Proline-Proline" in the sequence. |
| SP | Serine-Proline | The fraction of the dipeptide "Serine-Proline" in the sequence. |
| TP | Threonine-Proline | The fraction of the dipeptide "Threonine-Proline" in the sequence. |
| WP | Tryptophan-Proline | The fraction of the dipeptide "Tryptophan-Proline" in the sequence. |
| YP | Tyrosine-Proline | The fraction of the dipeptide "Tyrosine-Proline" in the sequence. |
| VP | Valine-Proline | The fraction of the dipeptide "Valine-Proline" in the sequence. |
| AS | Alanine-Serine | The fraction of the dipeptide "Alanine-Serine" in the sequence. |
| RS | Arginine-Serine | The fraction of the dipeptide "Arginine-Serine" in the sequence. |
| NS | Asparagine-Serine | The fraction of the dipeptide "Asparagine-Serine" in the sequence. |
| DS | Aspartic acid-Serine | The fraction of the dipeptide "Aspartic acid-Serine" in the |

| | | |
|---|---|---|
| | | sequence. |
| CS | Cysteine-Serine | The fraction of the dipeptide "Cysteine-Serine" in the sequence. |
| ES | Glutamic acid-Serine | The fraction of the dipeptide "Glutamic acid-Serine" in the sequence. |
| QS | Glutamine-Serine | The fraction of the dipeptide "Glutamine-Serine" in the sequence. |
| GS | Glycine-Serine | The fraction of the dipeptide "Glycine-Serine" in the sequence. |
| HS | Histidine-Serine | The fraction of the dipeptide "Histidine-Serine" in the sequence. |
| IS | Isoleucine-Serine | The fraction of the dipeptide "Isoleucine-Serine" in the sequence. |
| LS | Leucine-Serine | The fraction of the dipeptide "Leucine-Serine" in the sequence. |
| KS | Lysine-Serine | The fraction of the dipeptide "Lysine-Serine" in the sequence. |
| MS | Methionine-Serine | The fraction of the dipeptide "Methionine-Serine" in the sequence. |
| FS | Phenylalanine-Serine | The fraction of the dipeptide "Phenylalanine-Serine" in the sequence. |
| PS | Proline-Serine | The fraction of the dipeptide "Proline-Serine" in the sequence. |
| SS | Serine-Serine | The fraction of the |

| | | |
|---|---|---|
| | | dipeptide "Serine-Serine" in the sequence. |
| TS | Threonine-Serine | The fraction of the dipeptide "Threonine-Serine" in the sequence. |
| WS | Tryptophan-Serine | The fraction of the dipeptide "Tryptophan-Serine" in the sequence. |
| YS | Tyrosine-Serine | The fraction of the dipeptide "Tyrosine-Serine" in the sequence. |
| VS | Valine-Serine | The fraction of the dipeptide "Valine-Serine" in the sequence. |
| SP | Serine-Proline | The fraction of the dipeptide "Serine-Proline" in the sequence. |
| TP | Threonine-Proline | The fraction of the dipeptide "Threonine-Proline" in the sequence. |
| WP | Tryptophan-Proline | The fraction of the dipeptide "Tryptophan-Proline" in the sequence. |
| YP | Tyrosine-Proline | The fraction of the dipeptide "Tyrosine-Proline" in the sequence. |
| VP | Valine-Proline | The fraction of the dipeptide "Valine-Proline" in the sequence. |
| AS | Alanine-Serine | The fraction of the dipeptide "Alanine-Serine" in the sequence. |
| RS | Arginine-Serine | The fraction of the dipeptide "Arginine-Serine" in the sequence. |
| NS | Asparagine-Serine | The fraction of the |

| | | |
|---|---|---|
| | | dipeptide "Asparagine-Serine" in the sequence. |
| DS | Aspartic acid-Serine | The fraction of the dipeptide "Aspartic acid-Serine" in the sequence. |
| CS | Cysteine-Serine | The fraction of the dipeptide "Cysteine-Serine" in the sequence. |
| ES | Glutamic acid-Serine | The fraction of the dipeptide "Glutamic acid-Serine" in the sequence. |
| QS | Glutamine-Serine | The fraction of the dipeptide "Glutamine-Serine" in the sequence. |
| GS | Glycine-Serine | The fraction of the dipeptide "Glycine-Serine" in the sequence. |
| HS | Histidine-Serine | The fraction of the dipeptide "Histidine-Serine" in the sequence. |
| IS | Isoleucine-Serine | The fraction of the dipeptide "Isoleucine-Serine" in the sequence. |
| LS | Leucine-Serine | The fraction of the dipeptide "Leucine-Serine" in the sequence. |
| KS | Lysine-Serine | The fraction of the dipeptide "Lysine-Serine" in the sequence. |
| MS | Methionine-Serine | The fraction of the dipeptide "Methionine-Serine" in the sequence. |
| FS | Phenylalanine-Serine | The fraction of the dipeptide |

| | | |
|---|---|---|
| | | "Phenylalanine-Serine" in the sequence. |
| PS | Proline-Serine | The fraction of the dipeptide "Proline-Serine" in the sequence. |
| SS | Serine-Serine | The fraction of the dipeptide "Serine-Serine" in the sequence. |
| TS | Threonine-Serine | The fraction of the dipeptide "Threonine-Serine" in the sequence. |
| WS | Tryptophan-Serine | The fraction of the dipeptide "Tryptophan-Serine" in the sequence. |
| YS | Tyrosine-Serine | The fraction of the dipeptide "Tyrosine-Serine" in the sequence. |
| VS | Valine-Serine | The fraction of the dipeptide "Valine-Serine" in the sequence. |
| AT | Alanine-Threonine | The fraction of the dipeptide "Alanine-Threonine" in the sequence. |
| RT | Arginine-Threonine | The fraction of the dipeptide "Arginine-Threonine" in the sequence. |
| NT | Asparagine-Threonine | The fraction of the dipeptide "Asparagine-Threonine" in the sequence. |
| DT | Aspartic acid-Threonine | The fraction of the dipeptide "Aspartic acid-Threonine" in the sequence. |
| CT | Cysteine-Threonine | The fraction of the dipeptide "Cysteine-Threonine" in the sequence. |
| ET | Glutamic | The fraction of the |

| | | |
|---|---|---|
| | acid-Threonine | dipeptide "Glutamic acid-Threonine" in the sequence. |
| QT | Glutamine-Threonine | The fraction of the dipeptide "Glutamine-Threonine" in the sequence. |
| GT | Glycine-Threonine | The fraction of the dipeptide "Glycine-Threonine" in the sequence. |
| HT | Histidine-Threonine | The fraction of the dipeptide "Histidine-Threonine" in the sequence. |
| IT | Isoleucine-Threonine | The fraction of the dipeptide "Isoleucine-Threonine" in the sequence. |
| LT | Leucine-Threonine | The fraction of the dipeptide "Leucine-Threonine" in the sequence. |
| KT | Lysine-Threonine | The fraction of the dipeptide "Lysine-Threonine" in the sequence. |
| MT | Methionine-Threonine | The fraction of the dipeptide "Methionine-Threonine" in the sequence. |
| FT | Phenylalanine-Threonine | The fraction of the dipeptide "Phenylalanine-Threonine" in the sequence. |
| PT | Proline-Threonine | The fraction of the dipeptide "Proline-Threonine" in the sequence. |
| ST | Serine-Threonine | The fraction of the dipeptide "Serine-Threonine" in the sequence. |
| TT | Threonine-Threonine | The fraction of the |

| | | |
|---|---|---|
| | | dipeptide "Threonine-Threonine" in the sequence. |
| WT | Tryptophan-Threonine | The fraction of the dipeptide "Tryptophan-Threonine" in the sequence. |
| YT | Tyrosine-Threonine | The fraction of the dipeptide "Tyrosine-Threonine" in the sequence. |
| VT | Valine-Threonine | The fraction of the dipeptide "Valine-Threonine" in the sequence. |
| AW | Alanine-Tryptophan | The fraction of the dipeptide "Alanine-Tryptophan" in the sequence. |
| RW | Arginine-Tryptophan | The fraction of the dipeptide "Arginine-Tryptophan" in the sequence. |
| NW | Asparagine-Tryptophan | The fraction of the dipeptide "Asparagine-Tryptophan" in the sequence. |
| DW | Aspartic acid-Tryptophan | The fraction of the dipeptide "Aspartic acid-Tryptophan" in the sequence. |
| CW | Cysteine-Tryptophan | The fraction of the dipeptide "Cysteine-Tryptophan" in the sequence. |
| EW | Glutamic acid-Tryptophan | The fraction of the dipeptide "Glutamic acid-Tryptophan" in the sequence. |
| QW | Glutamine-Tryptophan | The fraction of the dipeptide "Glutamine-Tryptophan" in the sequence. |
| GW | Glycine-Tryptophan | The fraction of the |

| | | |
|---|---|---|
| | | dipeptide "Glycine-Tryptophan" in the sequence. |
| HW | Histidine-Tryptophan | The fraction of the dipeptide "Histidine-Tryptophan" in the sequence. |
| IW | Isoleucine-Tryptophan | The fraction of the dipeptide "Isoleucine-Tryptophan" in the sequence. |
| LW | Leucine-Tryptophan | The fraction of the dipeptide "Leucine-Tryptophan" in the sequence. |
| KW | Lysine-Tryptophan | The fraction of the dipeptide "Lysine-Tryptophan" in the sequence. |
| MW | Methionine-Tryptophan | The fraction of the dipeptide "Methionine-Tryptophan" in the sequence. |
| FW | Phenylalanine-Tryptophan | The fraction of the dipeptide "Phenylalanine-Tryptophan" in the sequence. |
| PW | Proline-Tryptophan | The fraction of the dipeptide "Proline-Tryptophan" in the sequence. |
| SW | Serine-Tryptophan | The fraction of the dipeptide "Serine-Tryptophan" in the sequence. |
| TW | Threonine-Tryptophan | The fraction of the dipeptide "Threonine-Tryptophan" in the sequence. |
| WW | Tryptophan-Tryptophan | The fraction of the dipeptide "Tryptophan-Tryptophan" in the sequence. |
| YW | Tyrosine-Tryptophan | The fraction of the |

| | | |
|---|---|---|
| | | dipeptide "Tyrosine-Tryptophan" in the sequence. |
| VW | Valine-Tryptophan | The fraction of the dipeptide "Valine-Tryptophan" in the sequence. |
| AY | Alanine-Tyrosine | The fraction of the dipeptide "Alanine-Tyrosine" in the sequence. |
| RY | Arginine-Tyrosine | The fraction of the dipeptide "Arginine-Tyrosine" in the sequence. |
| NY | Asparagine-Tyrosine | The fraction of the dipeptide "Asparagine-Tyrosine" in the sequence. |
| DY | Aspartic acid-Tyrosine | The fraction of the dipeptide "Aspartic acid-Tyrosine" in the sequence. |
| CY | Cysteine-Tyrosine | The fraction of the dipeptide "Cysteine-Tyrosine" in the sequence. |
| EY | Glutamic acid-Tyrosine | The fraction of the dipeptide "Glutamic acid-Tyrosine" in the sequence. |
| QY | Glutamine-Tyrosine | The fraction of the dipeptide "Glutamine-Tyrosine" in the sequence. |
| GY | Glycine-Tyrosine | The fraction of the dipeptide "Glycine-Tyrosine" in the sequence. |
| HY | Histidine-Tyrosine | The fraction of the dipeptide "Histidine-Tyrosine" in the sequence. |
| IY | Isoleucine-Tyrosine | The fraction of the |

| | | |
|---|---|---|
| | | dipeptide "Isoleucine-Tyrosine" in the sequence. |
| LY | Leucine-Tyrosine | The fraction of the dipeptide "Leucine-Tyrosine" in the sequence. |
| KY | Lysine-Tyrosine | The fraction of the dipeptide "Lysine-Tyrosine" in the sequence. |
| MY | Methionine-Tyrosine | The fraction of the dipeptide "Methionine-Tyrosine" in the sequence. |
| FY | Phenylalanine-Tyrosine | The fraction of the dipeptide "Phenylalanine-Tyrosine" in the sequence. |
| PY | Proline-Tyrosine | The fraction of the dipeptide "Proline-Tyrosine" in the sequence. |
| SY | Serine-Tyrosine | The fraction of the dipeptide "Serine-Tyrosine" in the sequence. |
| TY | Threonine-Tyrosine | The fraction of the dipeptide "Threonine-Tyrosine" in the sequence. |
| WY | Tryptophan-Tyrosine | The fraction of the dipeptide "Tryptophan-Tyrosine" in the sequence. |
| YY | Tyrosine-Tyrosine | The fraction of the dipeptide "Tyrosine-Tyrosine" in the sequence. |
| VY | Valine-Tyrosine | The fraction of the dipeptide "Valine-Tyrosine" in the sequence. |
| AV | Alanine-Valine | The fraction of the |

| | | |
|---|---|---|
| | | dipeptide "Alanine-Valine" in the sequence. |
| RV | Arginine-Valine | The fraction of the dipeptide "Arginine-Valine" in the sequence. |
| NV | Asparagine-Valine | The fraction of the dipeptide "Asparagine-Valine" in the sequence. |
| DV | Aspartic acid-Valine | The fraction of the dipeptide "Aspartic acid-Valine" in the sequence. |
| CV | Cysteine-Valine | The fraction of the dipeptide "Cysteine-Valine" in the sequence. |
| EV | Glutamic acid-Valine | The fraction of the dipeptide "Glutamic acid-Valine" in the sequence. |
| QV | Glutamine-Valine | The fraction of the dipeptide "Glutamine-Valine" in the sequence. |
| GV | Glycine-Valine | The fraction of the dipeptide "Glycine-Valine" in the sequence. |
| HV | Histidine-Valine | The fraction of the dipeptide "Histidine-Valine" in the sequence. |
| IV | Isoleucine-Valine | The fraction of the dipeptide "Isoleucine-Valine" in the sequence. |
| LV | Leucine-Valine | The fraction of the dipeptide "Leucine-Valine" in the sequence. |
| KV | Lysine-Valine | The fraction of the |

| | | |
|---|---|---|
| | | dipeptide "Lysine-Valine" in the sequence. |
| MV | Methionine-Valine | The fraction of the dipeptide "Methionine-Valine" in the sequence. |
| FV | Phenylalanine-Valine | The fraction of the dipeptide "Phenylalanine-Valine" in the sequence. |
| PV | Proline-Valine | The fraction of the dipeptide "Proline-Valine" in the sequence. |
| SV | Serine-Valine | The fraction of the dipeptide "Serine-Valine" in the sequence. |
| TV | Threonine-Valine | The fraction of the dipeptide "Threonine-Valine" in the sequence. |
| WV | Tryptophan-Valine | The fraction of the dipeptide "Tryptophan-Valine" in the sequence. |
| YV | Tyrosine-Valine | The fraction of the dipeptide "Tyrosine-Valine" in the sequence. |
| VV | Valine-Valine | The fraction of the dipeptide "Valine-Valine" in the sequence. |
| | Tyrosine-Tyrosine | The fraction of the dipeptide "Tyrosine-Tyrosine" in the sequence. |
| | Valine-Tyrosine | The fraction of the dipeptide "Valine-Tyrosine" in the sequence. |
| AV | Alanine-Valine | The fraction of the dipeptide "Alanine-Valine" in the sequence. |
| RV | Arginine-Valine | The fraction of the |

| | | |
|---|---|---|
| | | dipeptide "Arginine-Valine" in the sequence. |
| NV | Asparagine-Valine | The fraction of the dipeptide "Asparagine-Valine" in the sequence. |
| DV | Aspartic acid-Valine | The fraction of the dipeptide "Aspartic acid-Valine" in the sequence. |
| CV | Cysteine-Valine | The fraction of the dipeptide "Cysteine-Valine" in the sequence. |
| EV | Glutamic acid-Valine | The fraction of the dipeptide "Glutamic acid-Valine" in the sequence. |
| QV | Glutamine-Valine | The fraction of the dipeptide "Glutamine-Valine" in the sequence. |
| GV | Glycine-Valine | The fraction of the dipeptide "Glycine-Valine" in the sequence. |
| HV | Histidine-Valine | The fraction of the dipeptide "Histidine-Valine" in the sequence. |
| IV | Isoleucine-Valine | The fraction of the dipeptide "Isoleucine-Valine" in the sequence. |
| LV | Leucine-Valine | The fraction of the dipeptide "Leucine-Valine" in the sequence. |
| KV | Lysine-Valine | The fraction of the dipeptide "Lysine-Valine" in the sequence. |
| MV | Methionine-Valine | The fraction of the dipeptide |

| | | |
|---|---|---|
| | | "Methionine-Valine" in the sequence. |
| FV | Phenylalanine-Valine | The fraction of the dipeptide "Phenylalanine-Valine" in the sequence. |
| PV | Proline-Valine | The fraction of the dipeptide "Proline-Valine" in the sequence. |
| MoreauBroto_CIDH920105.lag1 | Moreau-Broto autocorrelation - CIDH920105 - lag 1 | Moreau-Broto autocorrelation of normalized average hydrophobicity scales with lag 1 |
| MoreauBroto_CIDH920105.lag2 | Moreau-Broto autocorrelation - CIDH920105 - lag 2 | Moreau-Broto autocorrelation of normalized average hydrophobicity scales with lag 2 |
| MoreauBroto_CIDH920105.lag3 | Moreau-Broto autocorrelation - CIDH920105 - lag 3 | Moreau-Broto autocorrelation of normalized average hydrophobicity scales with lag 3 |
| MoreauBroto_BHAR880101.lag1 | Moreau-Broto autocorrelation - BHAR880101 - lag 1 | Moreau-Broto autocorrelation of average flexibility indices with lag 1 |
| MoreauBroto_BHAR880101.lag2 | Moreau-Broto autocorrelation - BHAR880101 - lag 2 | Moreau-Broto autocorrelation of average flexibility indices with lag 2 |
| MoreauBroto_BHAR880101.lag3 | Moreau-Broto autocorrelation - BHAR880101 - lag 3 | Moreau-Broto autocorrelation of average flexibility indices with lag 3 |
| MoreauBroto_CHAM820101.lag1 | Moreau-Broto autocorrelation - CHAM820101 - lag 1 | Moreau-Broto autocorrelation of polarizability parameter with lag 1 |
| MoreauBroto_CHAM820101.lag2 | Moreau-Broto autocorrelation - CHAM820101 - lag 2 | Moreau-Broto autocorrelation of polarizability parameter with lag 2 |

| | | |
|---|---|---|
| MoreauBroto_CHAM820101.lag3 | Moreau-Broto autocorrelation - CHAM820101 - lag 3 | Moreau-Broto autocorrelation of polarizability parameter with lag 3 |
| MoreauBroto_CHAM820102.lag1 | Moreau-Broto autocorrelation - CHAM820102 - lag 1 | Moreau-Broto autocorrelation of free energy of solution in water with lag 1 |
| MoreauBroto_CHAM820102.lag2 | Moreau-Broto autocorrelation - CHAM820102 - lag 2 | Moreau-Broto autocorrelation of free energy of solution in water with lag 2 |
| MoreauBroto_CHAM820102.lag3 | Moreau-Broto autocorrelation - CHAM820102 - lag 3 | Moreau-Broto autocorrelation of free energy of solution in water with lag 3 |
| MoreauBroto_CHOC760101.lag1 | Moreau-Broto autocorrelation - CHOC760101 - lag 1 | Moreau-Broto autocorrelation of residue accessible surface area in tripeptide with lag 1 |
| MoreauBroto_CHOC760101.lag2 | Moreau-Broto autocorrelation - CHOC760101 - lag 2 | Moreau-Broto autocorrelation of residue accessible surface area in tripeptide with lag 2 |
| MoreauBroto_CHOC760101.lag3 | Moreau-Broto autocorrelation - CHOC760101 - lag 3 | Moreau-Broto autocorrelation of residue accessible surface area in tripeptide with lag 3 |
| MoreauBroto_BIGC670101.lag1 | Moreau-Broto autocorrelation - BIGC670101 - lag 1 | Moreau-Broto autocorrelation of residue volume with lag 1 |
| MoreauBroto_BIGC670101.lag2 | Moreau-Broto autocorrelation - BIGC670101 - lag 2 | Moreau-Broto autocorrelation of residue volume with lag 2 |
| MoreauBroto_BIGC670101.lag3 | Moreau-Broto autocorrelation - BIGC670101 - lag 3 | Moreau-Broto autocorrelation of residue volume with lag 3 |
| MoreauBroto_CHAM810101.lag1 | Moreau-Broto autocorrelation - CHAM810101 - lag 1 | Moreau-Broto autocorrelation of steric parameter with lag 1 |
| MoreauBroto_CHAM810101.lag2 | Moreau-Broto autocorrelation - CHAM810101 - lag 2 | Moreau-Broto autocorrelation of steric parameter with lag 2 |
| MoreauBroto_CHAM810101 | Moreau-Broto | Moreau-Broto |

| | | |
|---|---|---|
| .lag3 | autocorrelation - CHAM810101 - lag 3 | autocorrelation of steric parameter with lag 3 |
| Moran_CIDH920105.lag1 | Moran autocorrelation - CIDH920105 - lag 1 | Moran autocorrelation of normalized average hydrophobicity scales with lag 1 |
| Moran_CIDH920105.lag2 | Moran autocorrelation - CIDH920105 - lag 2 | Moran autocorrelation of normalized average hydrophobicity scales with lag 2 |
| Moran_CIDH920105.lag3 | Moran autocorrelation - CIDH920105 - lag 3 | Moran autocorrelation of normalized average hydrophobicity scales with lag 3 |
| Moran_BHAR880101.lag1 | Moran autocorrelation - BHAR880101 - lag 1 | Moran autocorrelation of average flexibility indices with lag 1 |
| Moran_BHAR880101.lag2 | Moran autocorrelation - BHAR880101 - lag 2 | Moran autocorrelation of average flexibility indices with lag 2 |
| Moran_BHAR880101.lag3 | Moran autocorrelation - BHAR880101 - lag 3 | Moran autocorrelation of average flexibility indices with lag 3 |
| Moran_CHAM820101.lag1 | Moran autocorrelation - CHAM820101 - lag 1 | Moran autocorrelation of polarizability parameter with lag 1 |
| Moran_CHAM820101.lag2 | Moran autocorrelation - CHAM820101 - lag 2 | Moran autocorrelation of polarizability parameter with lag 2 |
| Moran_CHAM820101.lag3 | Moran autocorrelation - CHAM820101 - lag 3 | Moran autocorrelation of polarizability parameter with lag 3 |
| Moran_CHAM820102.lag1 | Moran autocorrelation - CHAM820102 - lag 1 | Moran autocorrelation of free energy of solution in water with lag 1 |
| Moran_CHAM820102.lag2 | Moran autocorrelation - CHAM820102 - lag 2 | Moran autocorrelation of free energy of solution in water with lag 2 |
| Moran_CHAM820102.lag3 | Moran autocorrelation - CHAM820102 - lag 3 | Moran autocorrelation of free energy of solution in water with lag 3 |
| Moran_CHOC760101.lag1 | Moran autocorrelation - CHOC760101 - lag 1 | Moran autocorrelation of residue accessible surface area in tripeptide with lag |

| | | |
|---|---|---|
| | | 1 |
| Moran_CHOC760101.lag2 | Moran autocorrelation - CHOC760101 - lag 2 | Moran autocorrelation of residue accessible surface area in tripeptide with lag 2 |
| Moran_CHOC760101.lag3 | Moran autocorrelation - CHOC760101 - lag 3 | Moran autocorrelation of residue accessible surface area in tripeptide with lag 3 |
| Moran_BIGC670101.lag1 | Moran autocorrelation - BIGC670101 - lag 1 | Moran autocorrelation of residue volume with lag 1 |
| Moran_BIGC670101.lag2 | Moran autocorrelation - BIGC670101 - lag 2 | Moran autocorrelation of residue volume with lag 2 |
| Moran_BIGC670101.lag3 | Moran autocorrelation - BIGC670101 - lag 3 | Moran autocorrelation of residue volume with lag 3 |
| Moran_CHAM810101.lag1 | Moran autocorrelation - CHAM810101 - lag 1 | Moran autocorrelation of steric parameter with lag 1 |
| Moran_CHAM810101.lag2 | Moran autocorrelation - CHAM810101 - lag 2 | Moran autocorrelation of steric parameter with lag 2 |
| Moran_CHAM810101.lag3 | Moran autocorrelation - CHAM810101 - lag 3 | Moran autocorrelation of steric parameter with lag 3 |
| Geary_CIDH920105.lag1 | Geary autocorrelation - CIDH920105 - lag 1 | Geary autocorrelation of normalized average hydrophobicity scales with lag 1 |
| Geary_CIDH920105.lag2 | Geary autocorrelation - CIDH920105 - lag 2 | Geary autocorrelation of normalized average hydrophobicity scales with lag 2 |
| Geary_CIDH920105.lag3 | Geary autocorrelation - CIDH920105 - lag 3 | Geary autocorrelation of normalized average hydrophobicity scales with lag 3 |
| Geary_BHAR880101.lag1 | Geary autocorrelation - BHAR880101 - lag 1 | Geary autocorrelation of average flexibility indices with lag 1 |
| Geary_BHAR880101.lag2 | Geary autocorrelation - BHAR880101 - lag 2 | Geary autocorrelation of average flexibility indices with lag 2 |
| Geary_BHAR880101.lag3 | Geary autocorrelation - BHAR880101 - lag 3 | Geary autocorrelation of average flexibility indices with lag 3 |
| Geary_CHAM820101.lag1 | Geary autocorrelation - CHAM820101 - lag 1 | Geary autocorrelation of polarizability parameter |

| | | |
|---|---|---|
| | | with lag 1 |
| Geary_CHAM820101.lag2 | Geary autocorrelation - CHAM820101 - lag 2 | Geary autocorrelation of polarizability parameter with lag 2 |
| Geary_CHAM820101.lag3 | Geary autocorrelation - CHAM820101 - lag 3 | Geary autocorrelation of polarizability parameter with lag 3 |
| Geary_CHAM820102.lag1 | Geary autocorrelation - CHAM820102 - lag 1 | Geary autocorrelation of free energy of solution in water with lag 1 |
| Geary_CHAM820102.lag2 | Geary autocorrelation - CHAM820102 - lag 2 | Geary autocorrelation of free energy of solution in water with lag 2 |
| Geary_CHAM820102.lag3 | Geary autocorrelation - CHAM820102 - lag 3 | Geary autocorrelation of free energy of solution in water with lag 3 |
| Geary_CHOC760101.lag1 | Geary autocorrelation - CHOC760101 - lag 1 | Geary autocorrelation of residue accessible surface area in tripeptide with lag 1 |
| Geary_CHOC760101.lag2 | Geary autocorrelation - CHOC760101 - lag 2 | Geary autocorrelation of residue accessible surface area in tripeptide with lag 2 |
| Geary_CHOC760101.lag3 | Geary autocorrelation - CHOC760101 - lag 3 | Geary autocorrelation of residue accessible surface area in tripeptide with lag 3 |
| Geary_BIGC670101.lag1 | Geary autocorrelation - BIGC670101 - lag 1 | Geary autocorrelation of residue volume with lag 1 |
| Geary_BIGC670101.lag2 | Geary autocorrelation - BIGC670101 - lag 2 | Geary autocorrelation of residue volume with lag 2 |
| Geary_BIGC670101.lag3 | Geary autocorrelation - BIGC670101 - lag 3 | Geary autocorrelation of residue volume with lag 3 |
| Geary_CHAM810101.lag1 | Geary autocorrelation - CHAM810101 - lag 1 | Geary autocorrelation of steric parameter with lag 1 |
| Geary_CHAM810101.lag2 | Geary autocorrelation - CHAM810101 - lag 2 | Geary autocorrelation of steric parameter with lag 2 |
| Geary_CHAM810101.lag3 | Geary autocorrelation - CHAM810101 - lag 3 | Geary autocorrelation of steric parameter with lag 3 |
| hydrophobicity.Group1 | Hydrophobicity Group 1 | The fraction of polar amino acids (R, K, E, D, Q, N) within the sequence. |
| hydrophobicity.Group2 | Hydrophobicity Group 2 | The fraction of neutral |

| | | |
|---|---|---|
| | | amino acids (G, A, S, T, P, H, Y) within the sequence. |
| hydrophobicity.Group3 | Hydrophobicity Group 3 | The fraction of hydrophobic amino acids (C, L, V, I, M, F, W) within the sequence. |
| normwaalsvolume.Group1 | Normalized van der Waals Volume Group 1 | The fraction of amino acids with normalized van der Waals volume 0-2.78 (G, A, S, T, P, D, C) within the sequence. |
| normwaalsvolume.Group2 | Normalized van der Waals Volume Group 2 | The fraction of amino acids with normalized van der Waals volume 2.95-4.0 (N, V, E, Q, I, L) within the sequence. |
| normwaalsvolume.Group3 | Normalized van der Waals Volume Group 3 | The fraction of amino acids with normalized van der Waals volume 4.03-8.08 (M, H, K, F, R, Y, W) within the sequence. |
| polarity.Group1 | Polarity Group 1 | The fraction of amino acids with polarity 4.9-6.2 (L, I, F, W, C, M, V, Y) within the sequence. |
| polarity.Group2 | Polarity Group 2 | The fraction of amino acids with polarity 8.0-9.2 (P, A, T, G, S) within the sequence. |
| polarity.Group3 | Polarity Group 3 | The fraction of amino acids with polarity 10.4-13.0 (H, Q, R, K, N, E, D) within the sequence. |
| polarizability.Group1 | Polarizability Group 1 | The fraction of amino acids with polarizability 0-1.08 (G, A, S, D, T) within the sequence. |
| polarizability.Group2 | Polarizability Group 2 | The fraction of amino acids with polarizability 0.128-0.186 (C, P, N, V, E, Q, I, L) within the sequence. |
| polarizability.Group3 | Polarizability Group 3 | The fraction of amino |

| | | |
|---|---|---|
| | | acids with polarizability 0.219-0.409 (K, M, H, F, R, Y, W) within the sequence. |
| charge.Group1 | Charge Group 1 | The fraction of positively charged amino acids (K, R) within the sequence. |
| charge.Group2 | Charge Group 2 | The fraction of neutral amino acids (A, N, C, Q, G, H, I, L, M, F, P, S, T, W, Y, V) within the sequence. |
| charge.Group3 | Charge Group 3 | The fraction of negatively charged amino acids (D, E) within the sequence. |
| solventaccess.Group1 | Solvent Accessibility Group 1 | The fraction of buried amino acids (A, L, F, C, G, I, V, W) within the sequence. |
| solventaccess.Group2 | Solvent Accessibility Group 2 | The fraction of exposed amino acids (R, K, Q, E, N, D) within the sequence. |
| solventaccess.Group3 | Solvent Accessibility Group 3 | The fraction of intermediate solvent accessibility amino acids (M, S, P, T, H, Y) within the sequence. |
| prop1.Tr1221 | Hydrophobicity Transition 1221 | Transition frequency between polar (R, K, E, D, Q, N) and neutral (G, A, S, T, P, H, Y) amino acids |
| prop1.Tr1331 | Hydrophobicity Transition 1331 | Transition frequency between polar (R, K, E, D, Q, N) and hydrophobic (C, L, V, I, M, F, W) amino acids |
| prop1.Tr2332 | Hydrophobicity Transition 2332 | Transition frequency between neutral (G, A, S, T, P, H, Y) and hydrophobic (C, L, V, I, M, F, W) amino acids |
| prop2.Tr1221 | Normalized van der Waals Volume Transition 1221 | Transition frequency between small (0-2.78: G, A, S, T, P, D, C) and |

| | | |
|---|---|---|
| | | medium (2.95-4.0: N, V, E, Q, I, L) volume amino acids |
| prop2.Tr1331 | Normalized van der Waals Volume Transition 1331 | Transition frequency between small (0-2.78: G, A, S, T, P, D, C) and large (4.03-8.08: M, H, K, F, R, Y, W) volume amino acids |
| prop2.Tr2332 | Normalized van der Waals Volume Transition 2332 | Transition frequency between medium (2.95-4.0: N, V, E, Q, I, L) and large (4.03-8.08: M, H, K, F, R, Y, W) volume amino acids |
| prop3.Tr1221 | Polarity Transition 1221 | Transition frequency between low (4.9-6.2: L, I, F, W, C, M, V, Y) and medium (8.0-9.2: P, A, T, G, S) polarity amino acids |
| prop3.Tr1331 | Polarity Transition 1331 | Transition frequency between low (4.9-6.2: L, I, F, W, C, M, V, Y) and high (10.4-13.0: H, Q, R, K, N, E, D) polarity amino acids |
| prop3.Tr2332 | Polarity Transition 2332 | Transition frequency between medium (8.0-9.2: P, A, T, G, S) and high (10.4-13.0: H, Q, R, K, N, E, D) polarity amino acids |
| prop4.Tr1221 | Polarizability Transition 1221 | Transition frequency between low (0-1.08: G, A, S, D, T) and medium (0.128-0.186: C, P, N, V, E, Q, I, L) polarizability amino acids |
| prop4.Tr1331 | Polarizability Transition 1331 | Transition frequency between low (0-1.08: G, A, S, D, T) and high (0.219-0.409: K, M, H, F, R, Y, W) polarizability amino acids |
| prop4.Tr2332 | Polarizability Transition 2332 | Transition frequency between medium (0.128-0.186: C, P, N, V, |

| | | |
|---|---|---|
| | | E, Q, I, L) and high (0.219-0.409: K, M, H, F, R, Y, W) polarizability amino acids |
| prop5.Tr1221 | Charge Transition 1221 | Transition frequency between positive (K, R) and neutral (A, N, C, Q, G, H, I, L, M, F, P, S, T, W, Y, V) charged amino acids |
| prop5.Tr1331 | Charge Transition 1331 | Transition frequency between positive (K, R) and negative (D, E) charged amino acids |
| prop5.Tr2332 | Charge Transition 2332 | Transition frequency between neutral (A, N, C, Q, G, H, I, L, M, F, P, S, T, W, Y, V) and negative (D, E) charged amino acids |
| prop6.Tr1221 | Secondary Structure Transition 1221 | Transition frequency between helix-forming (E, A, L, M, Q, K, R, H) and strand-forming (V, I, Y, C, W, F, T) amino acids |
| prop6.Tr1331 | Secondary Structure Transition 1331 | Transition frequency between helix-forming (E, A, L, M, Q, K, R, H) and coil-forming (G, N, P, S, D) amino acids |
| prop6.Tr2332 | Secondary Structure Transition 2332 | Transition frequency between strand-forming (V, I, Y, C, W, F, T) and coil-forming (G, N, P, S, D) amino acids |
| prop7.Tr1221 | Solvent Accessibility Transition 1221 | Transition frequency between buried (A, L, F, C, G, I, V, W) and exposed (R, K, Q, E, N, D) amino acids |
| prop7.Tr1331 | Solvent Accessibility Transition 1331 | Transition frequency between buried (A, L, F, C, G, I, V, W) and intermediate (M, S, P, T, H, Y) solvent accessibility |

| | | |
|---|---|---|
| | | amino acids |
| prop7.Tr2332 | Solvent Accessibility Transition 2332 | Transition frequency between exposed (R, K, Q, E, N, D) and intermediate (M, S, P, T, H, Y) solvent accessibility amino acids |
| prop1.G1.residue0 | Hydrophobicity Group 1 Residue 0% | Percentage of polar amino acids (R, K, E, D, Q, N) at 0% of the sequence |
| prop1.G1.residue25 | Hydrophobicity Group 1 Residue 25% | Percentage of polar amino acids (R, K, E, D, Q, N) at 25% of the sequence |
| prop1.G1.residue50 | Hydrophobicity Group 1 Residue 50% | Percentage of polar amino acids (R, K, E, D, Q, N) at 50% of the sequence |
| prop1.G1.residue75 | Hydrophobicity Group 1 Residue 75% | Percentage of polar amino acids (R, K, E, D, Q, N) at 75% of the sequence |
| prop1.G1.residue100 | Hydrophobicity Group 1 Residue 100% | Percentage of polar amino acids (R, K, E, D, Q, N) at 100% of the sequence |
| prop1.G2.residue0 | Hydrophobicity Group 2 Residue 0% | Percentage of neutral amino acids (G, A, S, T, P, H, Y) at 0% of the sequence |
| prop1.G2.residue25 | Hydrophobicity Group 2 Residue 25% | Percentage of neutral amino acids (G, A, S, T, P, H, Y) at 25% of the sequence |
| prop1.G2.residue50 | Hydrophobicity Group 2 Residue 50% | Percentage of neutral amino acids (G, A, S, T, P, H, Y) at 50% of the sequence |
| prop1.G2.residue75 | Hydrophobicity Group 2 Residue 75% | Percentage of neutral amino acids (G, A, S, T, P, H, Y) at 75% of the sequence |
| prop1.G2.residue100 | Hydrophobicity Group 2 Residue 100% | Percentage of neutral amino acids (G, A, S, T, P, H, Y) at 100% of the sequence |
| prop1.G3.residue0 | Hydrophobicity Group 3 Residue 0% | Percentage of hydrophobic amino acids (C, L, V, I, M, |

| | | |
|---|---|---|
| | | F, W) at 0% of the sequence |
| prop1.G3.residue25 | Hydrophobicity Group 3 Residue 25% | Percentage of hydrophobic amino acids (C, L, V, I, M, F, W) at 25% of the sequence |
| prop1.G3.residue50 | Hydrophobicity Group 3 Residue 50% | Percentage of hydrophobic amino acids (C, L, V, I, M, F, W) at 50% of the sequence |
| prop1.G3.residue75 | Hydrophobicity Group 3 Residue 75% | Percentage of hydrophobic amino acids (C, L, V, I, M, F, W) at 75% of the sequence |
| prop1.G3.residue100 | Hydrophobicity Group 3 Residue 100% | Percentage of hydrophobic amino acids (C, L, V, I, M, F, W) at 100% of the sequence |
| prop2.G1.residue0 | Van der Waals Volume Group 1 Residue 0% | Percentage of amino acids with normalized van der Waals volume 0-2.78 (G, A, S, T, P, D, C) at 0% of the sequence |
| prop2.G1.residue25 | Van der Waals Volume Group 1 Residue 25% | Percentage of amino acids with normalized van der Waals volume 0-2.78 (G, A, S, T, P, D, C) at 25% of the sequence |
| prop2.G1.residue50 | Van der Waals Volume Group 1 Residue 50% | Percentage of amino acids with normalized van der Waals volume 0-2.78 (G, A, S, T, P, D, C) at 50% of the sequence |
| prop2.G1.residue75 | Van der Waals Volume Group 1 Residue 75% | Percentage of amino acids with normalized van der Waals volume 0-2.78 (G, A, S, T, P, D, C) at 75% of the sequence |
| prop2.G1.residue100 | Van der Waals Volume Group 1 Residue 100% | Percentage of amino acids with normalized van der Waals volume 0-2.78 (G, A, S, T, P, D, C) at 100% of the sequence |
| prop2.G2.residue0 | Van der Waals Volume | Percentage of amino acids |

| | | |
|---|---|---|
| | Group 2 Residue 0% | with normalized van der Waals volume 2.95-4.0 (N, V, E, Q, I, L) at 0% of the sequence |
| prop2.G2.residue25 | Van der Waals Volume Group 2 Residue 25% | Percentage of amino acids with normalized van der Waals volume 2.95-4.0 (N, V, E, Q, I, L) at 25% of the sequence |
| prop2.G2.residue50 | Van der Waals Volume Group 2 Residue 50% | Percentage of amino acids with normalized van der Waals volume 2.95-4.0 (N, V, E, Q, I, L) at 50% of the sequence |
| prop2.G2.residue75 | Van der Waals Volume Group 2 Residue 75% | Percentage of amino acids with normalized van der Waals volume 2.95-4.0 (N, V, E, Q, I, L) at 75% of the sequence |
| prop2.G2.residue100 | Van der Waals Volume Group 2 Residue 100% | Percentage of amino acids with normalized van der Waals volume 2.95-4.0 (N, V, E, Q, I, L) at 100% of the sequence |
| prop2.G3.residue0 | Van der Waals Volume Group 3 Residue 0% | Percentage of amino acids with normalized van der Waals volume 4.03-8.08 (M, H, K, F, R, Y, W) at 0% of the sequence |
| prop2.G3.residue25 | Van der Waals Volume Group 3 Residue 25% | Percentage of amino acids with normalized van der Waals volume 4.03-8.08 (M, H, K, F, R, Y, W) at 25% of the sequence |
| prop2.G3.residue50 | Van der Waals Volume Group 3 Residue 50% | Percentage of amino acids with normalized van der Waals volume 4.03-8.08 (M, H, K, F, R, Y, W) at 50% of the sequence |
| prop2.G3.residue75 | Van der Waals Volume Group 3 Residue 75% | Percentage of amino acids with normalized van der Waals volume 4.03-8.08 (M, H, K, F, R, Y, W) at 75% of the sequence |

| | | |
|---|---|---|
| prop2.G3.residue100 | Van der Waals Volume Group 3 Residue 100% | Percentage of amino acids with normalized van der Waals volume 4.03-8.08 (M, H, K, F, R, Y, W) at 100% of the sequence |
| prop3.G1.residue0 | Polarity Group 1 Residue 0% | Percentage of amino acids with polarity 4.9-6.2 (L, I, F, W, C, M, V, Y) at 0% of the sequence |
| prop3.G1.residue25 | Polarity Group 1 Residue 25% | Percentage of amino acids with polarity 4.9-6.2 (L, I, F, W, C, M, V, Y) at 25% of the sequence |
| prop3.G1.residue50 | Polarity Group 1 Residue 50% | Percentage of amino acids with polarity 4.9-6.2 (L, I, F, W, C, M, V, Y) at 50% of the sequence |
| prop3.G1.residue75 | Polarity Group 1 Residue 75% | Percentage of amino acids with polarity 4.9-6.2 (L, I, F, W, C, M, V, Y) at 75% of the sequence |
| prop3.G1.residue100 | Polarity Group 1 Residue 100% | Percentage of amino acids with polarity 4.9-6.2 (L, I, F, W, C, M, V, Y) at 100% of the sequence |
| prop3.G2.residue0 | Polarity Group 2 Residue 0% | Percentage of amino acids with polarity 8.0-9.2 (P, A, T, G, S) at 0% of the sequence |
| prop3.G2.residue25 | Polarity Group 2 Residue 25% | Percentage of amino acids with polarity 8.0-9.2 (P, A, T, G, S) at 25% of the sequence |
| prop3.G2.residue50 | Polarity Group 2 Residue 50% | Percentage of amino acids with polarity 8.0-9.2 (P, A, T, G, S) at 50% of the sequence |
| prop3.G2.residue75 | Polarity Group 2 Residue 75% | Percentage of amino acids with polarity 8.0-9.2 (P, A, T, G, S) at 75% of the sequence |
| prop3.G2.residue100 | Polarity Group 2 Residue 100% | Percentage of amino acids with polarity 8.0-9.2 (P, A, T, G, S) at 100% of the |

| | | |
|---|---|---|
| | | sequence |
| prop3.G3.residue0 | Polarity Group 3 Residue 0% | Percentage of amino acids with polarity 10.4-13.0 (H, Q, R, K, N, E, D) at 0% of the sequence |
| prop3.G3.residue25 | Polarity Group 3 Residue 25% | Percentage of amino acids with polarity 10.4-13.0 (H, Q, R, K, N, E, D) at 25% of the sequence |
| prop3.G3.residue50 | Polarity Group 3 Residue 50% | Percentage of amino acids with polarity 10.4-13.0 (H, Q, R, K, N, E, D) at 50% of the sequence |
| prop3.G3.residue75 | Polarity Group 3 Residue 75% | Percentage of amino acids with polarity 10.4-13.0 (H, Q, R, K, N, E, D) at 75% of the sequence |
| prop3.G3.residue100 | Polarity Group 3 Residue 100% | Percentage of amino acids with polarity 10.4-13.0 (H, Q, R, K, N, E, D) at 100% of the sequence |
| prop4.G1.residue0 | Polarizability Group 1 Residue 0% | Percentage of amino acids with polarizability 0-1.08 (G, A, S, D, T) at 0% of the sequence |
| prop4.G1.residue25 | Polarizability Group 1 Residue 25% | Percentage of amino acids with polarizability 0-1.08 (G, A, S, D, T) at 25% of the sequence |
| prop4.G1.residue50 | Polarizability Group 1 Residue 50% | Percentage of amino acids with polarizability 0-1.08 (G, A, S, D, T) at 50% of the sequence |
| prop4.G1.residue75 | Polarizability Group 1 Residue 75% | Percentage of amino acids with polarizability 0-1.08 (G, A, S, D, T) at 75% of the sequence |
| prop4.G1.residue100 | Polarizability Group 1 Residue 100% | Percentage of amino acids with polarizability 0-1.08 (G, A, S, D, T) at 100% of the sequence |
| prop4.G2.residue0 | Polarizability Group 2 Residue 0% | Percentage of amino acids with polarizability 0.128-0.186 (C, P, N, V, E, |

| | | |
|---|---|---|
| | | Q, I, L) at 0% of the sequence |
| prop4.G2.residue25 | Polarizability Group 2 Residue 25% | Percentage of amino acids with polarizability 0.128-0.186 (C, P, N, V, E, Q, I, L) at 25% of the sequence |
| prop4.G2.residue50 | Polarizability Group 2 Residue 50% | Percentage of amino acids with polarizability 0.128-0.186 (C, P, N, V, E, Q, I, L) at 50% of the sequence |
| prop4.G2.residue75 | Polarizability Group 2 Residue 75% | Percentage of amino acids with polarizability 0.128-0.186 (C, P, N, V, E, Q, I, L) at 75% of the sequence |
| prop4.G2.residue100 | Polarizability Group 2 Residue 100% | Percentage of amino acids with polarizability 0.128-0.186 (C, P, N, V, E, Q, I, L) at 100% of the sequence |
| prop4.G3.residue0 | Polarizability Group 3 Residue 0% | Percentage of amino acids with polarizability 0.219-0.409 (K, M, H, F, R, Y, W) at 0% of the sequence |
| prop4.G3.residue25 | Polarizability Group 3 Residue 25% | Percentage of amino acids with polarizability 0.219-0.409 (K, M, H, F, R, Y, W) at 25% of the sequence |
| prop4.G3.residue50 | Polarizability Group 3 Residue 50% | Percentage of amino acids with polarizability 0.219-0.409 (K, M, H, F, R, Y, W) at 50% of the sequence |
| prop4.G3.residue75 | Polarizability Group 3 Residue 75% | Percentage of amino acids with polarizability 0.219-0.409 (K, M, H, F, R, Y, W) at 75% of the sequence |
| prop4.G3.residue100 | Polarizability Group 3 Residue 100% | Percentage of amino acids with polarizability |

| | | |
|---|---|---|
| | | 0.219-0.409 (K, M, H, F, R, Y, W) at 100% of the sequence |
| prop5.G1.residue0 | Charge Group 1 Residue 0% | Percentage of positively charged amino acids (K, R) at 0% of the sequence |
| prop5.G1.residue25 | Charge Group 1 Residue 25% | Percentage of positively charged amino acids (K, R) at 25% of the sequence |
| prop5.G1.residue50 | Charge Group 1 Residue 50% | Percentage of positively charged amino acids (K, R) at 50% of the sequence |
| prop5.G1.residue75 | Charge Group 1 Residue 75% | Percentage of positively charged amino acids (K, R) at 75% of the sequence |
| prop5.G1.residue100 | Charge Group 1 Residue 100% | Percentage of positively charged amino acids (K, R) at 100% of the sequence |
| prop5.G2.residue0 | Charge Group 2 Residue 0% | Percentage of neutral amino acids (A, N, C, Q, G, H, I, L, M, F, P, S, T, W, Y, V) at 0% of the sequence |
| prop5.G2.residue25 | Charge Group 2 Residue 25% | Percentage of neutral amino acids (A, N, C, Q, G, H, I, L, M, F, P, S, T, W, Y, V) at 25% of the sequence |
| prop5.G2.residue50 | Charge Group 2 Residue 50% | Percentage of neutral amino acids (A, N, C, Q, G, H, I, L, M, F, P, S, T, W, Y, V) at 50% of the sequence |
| prop5.G2.residue75 | Charge Group 2 Residue 75% | Percentage of neutral amino acids (A, N, C, Q, G, H, I, L, M, F, P, S, T, W, Y, V) at 75% of the sequence |
| prop5.G2.residue100 | Charge Group 2 Residue 100% | Percentage of neutral amino acids (A, N, C, Q, G, H, I, L, M, F, P, S, T, W, Y, V) at 100% of the sequence |

| | | |
|---|---|---|
| prop5.G3.residue0 | Charge Group 3 Residue 0% | Percentage of negatively charged amino acids (D, E) at 0% of the sequence |
| prop5.G3.residue25 | Charge Group 3 Residue 25% | Percentage of negatively charged amino acids (D, E) at 25% of the sequence |
| prop5.G3.residue50 | Charge Group 3 Residue 50% | Percentage of negatively charged amino acids (D, E) at 50% of the sequence |
| prop5.G3.residue75 | Charge Group 3 Residue 75% | Percentage of negatively charged amino acids (D, E) at 75% of the sequence |
| prop5.G3.residue100 | Charge Group 3 Residue 100% | Percentage of negatively charged amino acids (D, E) at 100% of the sequence |
| prop7.G1.residue0 | Solvent Accessibility Group 1 Residue 0% | Percentage of buried amino acids (A, L, F, C, G, I, V, W) at 0% of the sequence |
| prop7.G1.residue25 | Solvent Accessibility Group 1 Residue 25% | Percentage of buried amino acids (A, L, F, C, G, I, V, W) at 25% of the sequence |
| prop7.G1.residue50 | Solvent Accessibility Group 1 Residue 50% | Percentage of buried amino acids (A, L, F, C, G, I, V, W) at 50% of the sequence |
| prop7.G1.residue75 | Solvent Accessibility Group 1 Residue 75% | Percentage of buried amino acids (A, L, F, C, G, I, V, W) at 75% of the sequence |
| prop7.G1.residue100 | Solvent Accessibility Group 1 Residue 100% | Percentage of buried amino acids (A, L, F, C, G, I, V, W) at 100% of the sequence |
| prop7.G2.residue0 | Solvent Accessibility Group 2 Residue 0% | Percentage of exposed amino acids (R, K, Q, E, N, D) at 0% of the sequence |
| prop7.G2.residue25 | Solvent Accessibility Group 2 Residue 25% | Percentage of exposed amino acids (R, K, Q, E, N, D) at 25% of the sequence |

| | | |
|---|---|---|
| prop7.G2.residue50 | Solvent Accessibility Group 2 Residue 50% | Percentage of exposed amino acids (R, K, Q, E, N, D) at 50% of the sequence |
| prop7.G2.residue75 | Solvent Accessibility Group 2 Residue 75% | Percentage of exposed amino acids (R, K, Q, E, N, D) at 75% of the sequence |
| prop7.G2.residue100 | Solvent Accessibility Group 2 Residue 100% | Percentage of exposed amino acids (R, K, Q, E, N, D) at 100% of the sequence |
| prop7.G3.residue0 | Solvent Accessibility Group 3 Residue 0% | Percentage of intermediate amino acids (M, S, P, T, H, Y) at 0% of the sequence |
| prop7.G3.residue25 | Solvent Accessibility Group 3 Residue 25% | Percentage of intermediate amino acids (M, S, P, T, H, Y) at 25% of the sequence |
| prop7.G3.residue50 | Solvent Accessibility Group 3 Residue 50% | Percentage of intermediate amino acids (M, S, P, T, H, Y) at 50% of the sequence |
| prop7.G3.residue75 | Solvent Accessibility Group 3 Residue 75% | Percentage of intermediate amino acids (M, S, P, T, H, Y) at 75% of the sequence |
| prop7.G3.residue100 | Solvent Accessibility Group 3 Residue 100% | Percentage of intermediate amino acids (M, S, P, T, H, Y) at 100% of the sequence |
| Schneider.lag1 | Schneider Sequence-Order-Coupling Number (Lag 1) | The first-rank sequence-order-coupling number based on Schneider's distance matrix between amino acids |
| Schneider.lag2 | Schneider Sequence-Order-Coupling Number (Lag 2) | The second-rank sequence-order-coupling number based on Schneider's distance matrix between amino acids |
| Schneider.lag3 | Schneider Sequence-Order-Coupling Number (Lag 3) | The third-rank sequence-order-coupling number based on Schneider's distance |

| | | |
|---|---|---|
| | | matrix between amino acids |
| Grantham.lag1 | Grantham Sequence-Order-Coupling Number (Lag 1) | The first-rank sequence-order-coupling number based on Grantham's distance matrix between amino acids |
| Grantham.lag2 | Grantham Sequence-Order-Coupling Number (Lag 2) | The second-rank sequence-order-coupling number based on Grantham's distance matrix between amino acids |
| Grantham.lag3 | Grantham Sequence-Order-Coupling Number (Lag 3) | The third-rank sequence-order-coupling number based on Grantham's distance matrix between amino acids |
| Schneider.Xr.A | Schneider Quasi-Sequence-Order for Alanine | Quasi-sequence-order descriptor for Alanine based on Schneider's matrix |
| Schneider.Xr.R | Schneider Quasi-Sequence-Order for Arginine | Quasi-sequence-order descriptor for Arginine based on Schneider's matrix |
| Schneider.Xr.N | Schneider Quasi-Sequence-Order for Asparagine | Quasi-sequence-order descriptor for Asparagine based on Schneider's matrix |
| Schneider.Xr.D | Schneider Quasi-Sequence-Order for Aspartic Acid | Quasi-sequence-order descriptor for Aspartic Acid based on Schneider's matrix |
| Schneider.Xr.C | Schneider Quasi-Sequence-Order for Cysteine | Quasi-sequence-order descriptor for Cysteine based on Schneider's matrix |
| Schneider.Xr.E | Schneider Quasi-Sequence-Order for Glutamic Acid | Quasi-sequence-order descriptor for Glutamic Acid based on Schneider's matrix |

| | | |
|---|---|---|
| Schneider.Xr.Q | Schneider Quasi-Sequence-Order for Glutamine | Quasi-sequence-order descriptor for Glutamine based on Schneider's matrix |
| Schneider.Xr.G | Schneider Quasi-Sequence-Order for Glycine | Quasi-sequence-order descriptor for Glycine based on Schneider's matrix |
| Schneider.Xr.H | Schneider Quasi-Sequence-Order for Histidine | Quasi-sequence-order descriptor for Histidine based on Schneider's matrix |
| Schneider.Xr.I | Schneider Quasi-Sequence-Order for Isoleucine | Quasi-sequence-order descriptor for Isoleucine based on Schneider's matrix |
| Schneider.Xr.L | Schneider Quasi-Sequence-Order for Leucine | Quasi-sequence-order descriptor for Leucine based on Schneider's matrix |
| Schneider.Xr.K | Schneider Quasi-Sequence-Order for Lysine | Quasi-sequence-order descriptor for Lysine based on Schneider's matrix |
| Schneider.Xr.M | Schneider Quasi-Sequence-Order for Methionine | Quasi-sequence-order descriptor for Methionine based on Schneider's matrix |
| Schneider.Xr.F | Schneider Quasi-Sequence-Order for Phenylalanine | Quasi-sequence-order descriptor for Phenylalanine based on Schneider's matrix |
| Schneider.Xr.P | Schneider Quasi-Sequence-Order for Proline | Quasi-sequence-order descriptor for Proline based on Schneider's matrix |
| Schneider.Xr.S | Schneider Quasi-Sequence-Order for Serine | Quasi-sequence-order descriptor for Serine based on Schneider's matrix |
| Schneider.Xr.T | Schneider Quasi-Sequence-Order for Threonine | Quasi-sequence-order descriptor for Threonine based on Schneider's matrix |
| Schneider.Xr.W | Schneider | Quasi-sequence-order |

| | | |
|---|---|---|
| | Quasi-Sequence-Order for Tryptophan | descriptor for Tryptophan based on Schneider's matrix |
| Schneider.Xr.Y | Schneider Quasi-Sequence-Order for Tyrosine | Quasi-sequence-order descriptor for Tyrosine based on Schneider's matrix |
| Schneider.Xr.V | Schneider Quasi-Sequence-Order for Valine | Quasi-sequence-order descriptor for Valine based on Schneider's matrix |
| Grantham.Xr.A | Grantham Quasi-Sequence-Order for Alanine | Quasi-sequence-order descriptor for Alanine based on Grantham's matrix |
| Grantham.Xr.R | Grantham Quasi-Sequence-Order for Arginine | Quasi-sequence-order descriptor for Arginine based on Grantham's matrix |
| Grantham.Xr.N | Grantham Quasi-Sequence-Order for Asparagine | Quasi-sequence-order descriptor for Asparagine based on Grantham's matrix |
| Grantham.Xr.D | Grantham Quasi-Sequence-Order for Aspartic Acid | Quasi-sequence-order descriptor for Aspartic Acid based on Grantham's matrix |
| Grantham.Xr.C | Grantham Quasi-Sequence-Order for Cysteine | Quasi-sequence-order descriptor for Cysteine based on Grantham's matrix |
| Grantham.Xr.E | Grantham Quasi-Sequence-Order for Glutamic Acid | Quasi-sequence-order descriptor for Glutamic Acid based on Grantham's matrix |
| Grantham.Xr.Q | Grantham Quasi-Sequence-Order for Glutamine | Quasi-sequence-order descriptor for Glutamine based on Grantham's matrix |
| Grantham.Xr.G | Grantham Quasi-Sequence-Order for Glycine | Quasi-sequence-order descriptor for Glycine based on Grantham's matrix |
| Grantham.Xr.H | Grantham Quasi-Sequence-Order | Quasi-sequence-order descriptor for Histidine |

| | | |
|---|---|---|
| | for Histidine | based on Grantham's matrix |
| Grantham.Xr.I | Grantham Quasi-Sequence-Order for Isoleucine | Quasi-sequence-order descriptor for Isoleucine based on Grantham's matrix |
| Grantham.Xr.L | Grantham Quasi-Sequence-Order for Leucine | Quasi-sequence-order descriptor for Leucine based on Grantham's matrix |
| Grantham.Xr.K | Grantham Quasi-Sequence-Order for Lysine | Quasi-sequence-order descriptor for Lysine based on Grantham's matrix |
| Grantham.Xr.M | Grantham Quasi-Sequence-Order for Methionine | Quasi-sequence-order descriptor for Methionine based on Grantham's matrix |
| Grantham.Xr.F | Grantham Quasi-Sequence-Order for Phenylalanine | Quasi-sequence-order descriptor for Phenylalanine based on Grantham's matrix |
| Grantham.Xr.P | Grantham Quasi-Sequence-Order for Proline | Quasi-sequence-order descriptor for Proline based on Grantham's matrix |
| Grantham.Xr.S | Grantham Quasi-Sequence-Order for Serine | Quasi-sequence-order descriptor for Serine based on Grantham's matrix |
| Grantham.Xr.T | Grantham Quasi-Sequence-Order for Threonine | Quasi-sequence-order descriptor for Threonine based on Grantham's matrix |
| Grantham.Xr.W | Grantham Quasi-Sequence-Order for Tryptophan | Quasi-sequence-order descriptor for Tryptophan based on Grantham's matrix |
| Grantham.Xr.Y | Grantham Quasi-Sequence-Order for Tyrosine | Quasi-sequence-order descriptor for Tyrosine based on Grantham's matrix |
| Grantham.Xr.V | Grantham Quasi-Sequence-Order for Valine | Quasi-sequence-order descriptor for Valine based on Grantham's matrix |

| | | |
|---|---|---|
| Schneider.Xd.1 | Schneider 1st-rank Quasi-Sequence-Order | Quasi-sequence-order descriptor for 1st rank sequence-order-coupling, based on Schneider-Wrede matrix. |
| Schneider.Xd.2 | Schneider 2nd-rank Quasi-Sequence-Order | Quasi-sequence-order descriptor for 2nd rank sequence-order-coupling, based on Schneider-Wrede matrix. |
| Schneider.Xd.3 | Schneider 3rd-rank Quasi-Sequence-Order | Quasi-sequence-order descriptor for 3rd rank sequence-order-coupling, based on Schneider-Wrede matrix. |
| Grantham.Xd.1 | Grantham 1st-rank Quasi-Sequence-Order | Quasi-sequence-order descriptor for 1st rank sequence-order-coupling, based on Grantham matrix. |
| Grantham.Xd.2 | Grantham 2nd-rank Quasi-Sequence-Order | Quasi-sequence-order descriptor for 2nd rank sequence-order-coupling, based on Grantham matrix. |
| Grantham.Xd.3 | Grantham 3rd-rank Quasi-Sequence-Order | Quasi-sequence-order descriptor for 3rd rank sequence-order-coupling, based on Grantham matrix. |
| Xc1.A | PseAAC for Alanine | Pseudo-amino acid composition for alanine based on hydrophobicity, hydrophilicity, and side chain mass |
| Xc1.R | PseAAC for Arginine | Pseudo-amino acid composition for arginine based on hydrophobicity, hydrophilicity, and side chain mass |
| Xc1.N | PseAAC for Asparagine | Pseudo-amino acid composition for asparagine based on hydrophobicity, |

| | | |
|---|---|---|
| | | hydrophilicity, and side chain mass |
| Xc1.D | PseAAC for Aspartic Acid | Pseudo-amino acid composition for aspartic acid based on hydrophobicity, hydrophilicity, and side chain mass |
| Xc1.C | PseAAC for Cysteine | Pseudo-amino acid composition for cysteine based on hydrophobicity, hydrophilicity, and side chain mass |
| Xc1.E | PseAAC for Glutamic Acid | Pseudo-amino acid composition for glutamic acid based on hydrophobicity, hydrophilicity, and side chain mass |
| Xc1.Q | PseAAC for Glutamine | Pseudo-amino acid composition for glutamine based on hydrophobicity, hydrophilicity, and side chain mass |
| Xc1.G | PseAAC for Glycine | Pseudo-amino acid composition for glycine based on hydrophobicity, hydrophilicity, and side chain mass |
| Xc1.H | PseAAC for Histidine | Pseudo-amino acid composition for histidine based on hydrophobicity, hydrophilicity, and side chain mass |
| Xc1.I | PseAAC for Isoleucine | Pseudo-amino acid composition for isoleucine based on hydrophobicity, hydrophilicity, and side chain mass |
| Xc1.L | PseAAC for Leucine | Pseudo-amino acid composition for leucine based on hydrophobicity, hydrophilicity, and side chain mass |

| | | |
|---|---|---|
| Xc1.K | PseAAC for Lysine | Pseudo-amino acid composition for lysine based on hydrophobicity, hydrophilicity, and side chain mass |
| Xc1.M | PseAAC for Methionine | Pseudo-amino acid composition for methionine based on hydrophobicity, hydrophilicity, and side chain mass |
| Xc1.F | PseAAC for Phenylalanine | Pseudo-amino acid composition for phenylalanine based on hydrophobicity, hydrophilicity, and side chain mass |
| Xc1.P | PseAAC for Proline | Pseudo-amino acid composition for proline based on hydrophobicity, hydrophilicity, and side chain mass |
| Xc1.S | PseAAC for Serine | Pseudo-amino acid composition for serine based on hydrophobicity, hydrophilicity, and side chain mass |
| Xc1.T | PseAAC for Threonine | Pseudo-amino acid composition for threonine based on hydrophobicity, hydrophilicity, and side chain mass |
| Xc1.W | PseAAC for Tryptophan | Pseudo-amino acid composition for tryptophan based on hydrophobicity, hydrophilicity, and side chain mass |
| Xc1.Y | PseAAC for Tyrosine | Pseudo-amino acid composition for tyrosine based on hydrophobicity, hydrophilicity, and side chain mass |
| Xc1.V | PseAAC for Valine | Pseudo-amino acid |

| | | |
|---|---|---|
| | | composition for valine based on hydrophobicity, hydrophilicity, and side chain mass |
| Xc2.lambda.1 | 1st-tier Sequence Coupling | Pseudo-amino acid composition for 1st-rank sequence-order-coupling based on hydrophobicity, hydrophilicity, and side chain mass |
| Xc2.lambda.2 | 2nd-tier Sequence Coupling | Pseudo-amino acid composition for 2nd-rank sequence-order-coupling based on hydrophobicity, hydrophilicity, and side chain mass |
| Xc2.lambda.3 | 3rd-tier Sequence Coupling | Pseudo-amino acid composition for 3rd-rank sequence-order-coupling based on hydrophobicity, hydrophilicity, and side chain mass |
| Pc1.A | Amphiphilic pseudo-amino acid composition for alanine | Reflects the contribution of alanine to the sequence, weighted by hydrophobicity and hydrophilicity correlations. |
| Pc1.R | Amphiphilic pseudo-amino acid composition for arginine | Reflects the contribution of arginine to the sequence, weighted by hydrophobicity and hydrophilicity correlations. |
| Pc1.N | Amphiphilic pseudo-amino acid composition for asparagine | Reflects the contribution of asparagine to the sequence, weighted by hydrophobicity and hydrophilicity correlations. |
| Pc1.D | Amphiphilic pseudo-amino acid composition for aspartic acid | Reflects the contribution of aspartic acid to the sequence, weighted by hydrophobicity and |

| | | |
|---|---|---|
| | | hydrophilicity correlations. |
| Pc1.C | Amphiphilic pseudo-amino acid composition for cysteine | Reflects the contribution of cysteine to the sequence, weighted by hydrophobicity and hydrophilicity correlations. |
| Pc1.E | Amphiphilic pseudo-amino acid composition for glutamic acid | Reflects the contribution of glutamic acid to the sequence, weighted by hydrophobicity and hydrophilicity correlations. |
| Pc1.Q | Amphiphilic pseudo-amino acid composition for glutamine | Reflects the contribution of glutamine to the sequence, weighted by hydrophobicity and hydrophilicity correlations. |
| Pc1.G | Amphiphilic pseudo-amino acid composition for glycine | Reflects the contribution of glycine to the sequence, weighted by hydrophobicity and hydrophilicity correlations. |
| Pc1.H | Amphiphilic pseudo-amino acid composition for histidine | Reflects the contribution of histidine to the sequence, weighted by hydrophobicity and hydrophilicity correlations. |
| Pc1.I | Amphiphilic pseudo-amino acid composition for isoleucine | Reflects the contribution of isoleucine to the sequence, weighted by hydrophobicity and hydrophilicity correlations. |
| Pc1.L | Amphiphilic pseudo-amino acid composition for leucine | Reflects the contribution of leucine to the sequence, weighted by hydrophobicity and hydrophilicity correlations. |

| | | |
|---|---|---|
| Pc1.K | Amphiphilic pseudo-amino acid composition for lysine | Reflects the contribution of lysine to the sequence, weighted by hydrophobicity and hydrophilicity correlations. |
| Pc1.M | Amphiphilic pseudo-amino acid composition for methionine | Reflects the contribution of methionine to the sequence, weighted by hydrophobicity and hydrophilicity correlations. |
| Pc1.F | Amphiphilic pseudo-amino acid composition for phenylalanine | Reflects the contribution of phenylalanine to the sequence, weighted by hydrophobicity and hydrophilicity correlations. |
| Pc1.P | Amphiphilic pseudo-amino acid composition for proline | Reflects the contribution of proline to the sequence, weighted by hydrophobicity and hydrophilicity correlations. |
| Pc1.S | Amphiphilic pseudo-amino acid composition for serine | Reflects the contribution of serine to the sequence, weighted by hydrophobicity and hydrophilicity correlations. |
| Pc1.T | Amphiphilic pseudo-amino acid composition for threonine | Reflects the contribution of threonine to the sequence, weighted by hydrophobicity and hydrophilicity correlations. |
| Pc1.W | Amphiphilic pseudo-amino acid composition for tryptophan | Reflects the contribution of tryptophan to the sequence, weighted by hydrophobicity and hydrophilicity correlations. |
| Pc1.Y | Amphiphilic pseudo-amino acid | Reflects the contribution of tyrosine to the |

| | | |
|---|---|---|
| | composition for tyrosine | sequence, weighted by hydrophobicity and hydrophilicity correlations. |
| Pc1.V | Amphiphilic pseudo-amino acid composition for valine | Reflects the contribution of valine to the sequence, weighted by hydrophobicity and hydrophilicity correlations. |
| Pc2.Hydrophobicity.1 | 1st-rank sequence-order coupling factor for hydrophobicity | Reflects the correlation between adjacent residues based on hydrophobicity. |
| Pc2.Hydrophilicity.1 | 1st-rank sequence-order coupling factor for hydrophilicity | Reflects the correlation between adjacent residues based on hydrophilicity. |
| Pc2.Hydrophobicity.2 | 2nd-rank sequence-order coupling factor for hydrophobicity | Reflects the correlation between residues separated by one position based on hydrophobicity. |
| Pc2.Hydrophilicity.2 | 2nd-rank sequence-order coupling factor for hydrophilicity | Reflects the correlation between residues separated by one position based on hydrophilicity. |
| Pc2.Hydrophobicity.3 | 3rd-rank sequence-order coupling factor for hydrophobicity | Reflects the correlation between residues separated by two positions based on hydrophobicity. |
| Pc2.Hydrophilicity.3 | 3rd-rank sequence-order coupling factor for hydrophilicity | Reflects the correlation between residues separated by two positions based on hydrophilicity. |
| net.charge | Net Charge | This variable represents the theoretical net charge of a peptide sequence, which is calculated using the Henderson-Hasselbalch equation at a defined pH. |
| hydrophobicity.value | Hydrophobicity Index | This variable represents the GRAVY (Grand Average of Hydropathy) hydrophobicity index of an amino acids sequence, |

| | | |
|---|---|---|
| | | computed using one of the 38 scales from different sources. |
| boman.index | Boman Index | This variable represents the potential peptide interaction index proposed by Boman, which estimates the potential of a peptide to bind to membranes or other proteins. |
| tiny | Tiny | This variable represents the composition of tiny amino acids (A + C + G + S + T) in a peptide sequence. |
| small | Small | This variable represents the composition of small amino acids (A + B + C + D + G + N + P + S + T + V) in a peptide sequence. |
| aliphatic | Aliphatic | This variable represents the composition of aliphatic amino acids (A + I + L + V) in a peptide sequence. |
| aromatic | Aromatic | This variable represents the composition of aromatic amino acids (F + H + W + Y) in a peptide sequence. |
| nonpolar | Non-polar | This variable represents the composition of non-polar amino acids (A + C + F + G + I + L + M + P + V + W + Y) in a peptide sequence. |
| polar | Polar | This variable represents the composition of polar amino acids (D + E + H + K + N + Q + R + S + T + Z) in a peptide sequence. |
| charged | Charged | This variable represents the composition of |

| | | |
|---|---|---|
| | | charged amino acids (B + D + E + H + K + R + Z) in a peptide sequence. |
| basic | Basic | This variable represents the composition of basic amino acids (H + K + R) in a peptide sequence. |
| acidic | Acidic | This variable represents the composition of acidic amino acids (B + D + E + Z) in a peptide sequence. |
| molecular_weight | Molecular weight | This variable represents the molecular weight of a peptide sequence. |
